# APOBEC shapes tumor evolution and age at onset of lung cancer in smokers

**DOI:** 10.1101/2024.04.02.587805

**Authors:** Tongwu Zhang, Jian Sang, Phuc H. Hoang, Wei Zhao, Jennifer Rosenbaum, Kofi Ennu Johnson, Leszek J. Klimczak, John McElderry, Alyssa Klein, Christopher Wirth, Erik N. Bergstrom, Marcos Díaz-Gay, Raviteja Vangara, Frank Colon-Matos, Amy Hutchinson, Scott M. Lawrence, Nathan Cole, Bin Zhu, Teresa M. Przytycka, Jianxin Shi, Neil E. Caporaso, Robert Homer, Angela C. Pesatori, Dario Consonni, Marcin Imielinski, Stephen J. Chanock, David C. Wedge, Dmitry A. Gordenin, Ludmil B. Alexandrov, Reuben S. Harris, Maria Teresa Landi

**Author notes:** Corresponding author: Maria Teresa Landi. These authors contributed equally to this work (Tongwu Zhang, Jian Sang).

## Abstract

APOBEC enzymes are part of the innate immunity and are responsible for restricting viruses and retroelements by deaminating cytosine residues^1,2^. Most solid tumors harbor different levels of somatic mutations attributed to the off-target activities of APOBEC3A (A3A) and/or APOBEC3B (A3B)^3–6^. However, how APOBEC3A/B enzymes shape the tumor evolution in the presence of exogenous mutagenic processes is largely unknown. Here, by combining deep whole-genome sequencing with multi-omics profiling of 309 lung cancers from smokers with detailed tobacco smoking information, we identify two subtypes defined by low (*LAS*) and high (*HAS*) APOBEC mutagenesis. LAS are enriched for A3B-like mutagenesis and *KRAS* mutations, whereas HAS for A3A-like mutagenesis and *TP53* mutations. Unlike *APOBEC3A*, *APOBEC3B* expression is strongly associated with an upregulation of the base excision repair pathway. Hypermutation by unrepaired A3A and tobacco smoking mutagenesis combined with *TP53*-induced genomic instability can trigger senescence^7^, apoptosis^8^, and cell regeneration^9^, as indicated by high expression of pulmonary healing signaling pathway, stemness markers and distal cell-of-origin in HAS. The expected association of tobacco smoking variables (e.g., time to first cigarette) with genomic/epigenomic changes are not observed in HAS, a plausible consequence of frequent cell senescence or apoptosis. HAS have more neoantigens, slower clonal expansion, and older age at onset compared to LAS, particularly in heavy smokers, consistent with high proportions of newly generated, unmutated cells and frequent immuno-editing. These findings show how heterogeneity in mutational burden across co-occurring mutational processes and cell types contributes to tumor development, with important clinical implications.

## INTRODUCTION

Somatic mutations in cancer are caused by both exogenous and endogenous mutational processes^10^. Mutational signature analysis has been widely used to reveal the mutagenic processes in cancer genomic studies^4,11,12^. Varying levels of APOBEC mutational signatures (a combination of COSMIC mutational signatures SBS2 and SBS13 or an enrichment with a known APOBEC mutational motif TCW; W=adenine/thymine) have been reported in ∼70% of human cancer types^3–6^ and have been associated with an improved response to immunotherapy in some cancer types^13^. APOBEC mutagenesis is the off-target effect of the activity of the APOBEC family of enzymes that function as cytosine deaminases^14^. This off-target effect has been predominantly attributed to APOBEC3A (A3A) and APOBEC3B (A3B)^15^, which are involved in virus and retroelement restriction^16,17^, response to DNA damage, and other cellular functions^1,21^. Both enzymes have access to the nucleus and are associated with elevated mRNA levels in tumors as compared with matched normal tissue^3,18^. While A3A and A3B imprint similar patterns of mutations on somatic genomes, the two enzymes exhibit enrichments for different motifs. Specifically, an enrichment of YT<u>C</u>A has been associated with APOBEC3A whereas an enrichment of RT<u>C</u>A has been attributed to APOBEC3B (Y=purine; R=pyrimidine)^19^. The occurrence of most APOBEC associated mutations appears to be episodic^20–22^ and this process is likely driven by the APOBEC3A deaminase^6^. APOBEC associated deamination of cytosines can occur exclusively in single-stranded (ss)DNA^1^. APOBEC mutagenesis in tumors is enriched in the lagging strand template and in early-replicating, gene-dense, and active chromatin genome regions^23–25^. APOBEC3-catalyzed mutagenesis can also lead to chromosomal instability^26,27^. Despite considerable information accumulated about off-target mutagenesis by different APOBECs in tumors, their roles in tumor evolution in the context of other mutagenic processes remain largely unknown.

Lung cancer is the second most common cancer type worldwide and the leading cause of cancer death^28^. Tobacco smoking is the foremost risk factor for lung cancer and has strong mutagenic activity leading to high tumor mutational burden (TMB), COSMIC mutational signatures SBS4, DBS2, ID3 and more recently SBS92^4,29,30^, and increased mutation frequency in cancer driver genes^31–33^. Lung cancer also harbors APOBEC signatures in about 35 to 55% of tumors^4,5^, possibly in response to DNA damage and local inflammation (likely induced by tobacco smoking)^2,34^. However, how the co-occurrence of APOBEC and tobacco smoking mutagenesis impact lung cancer evolution has not been investigated.

Here, we present a multi-omics study including newly sequenced deep whole-genome (73x coverage), transcriptome, and epigenomic profiles of 309 paired tumor-normal lung tissue samples from smokers with detailed clinical and smoking history information. We found that the co-occurring of APOBEC and tobacco smoking mutational processes affects the development and age at onset of lung cancer. A similar effect is observed in other tumor types associated with exposure to exogenous mutagens like melanoma.

## RESULTS

We analyzed a multi-omics dataset encompassing a total of 345 smokers with histologically confirmed lung cancer (**Supplementary Table 1**). While the majority of tumors show dominant tobacco smoking-associated signatures SBS4, DBS2, ID3, and few SBS92 as expected (**Supplementary Table 2**), 36 tumors (10.4%) had no mutations assigned to SBS4, even in the subset of 20 (9.2%) samples from the Environment And Genetics in Lung cancer Etiology (EAGLE) study^35^ with very high-confidence and detailed smoking information (**Supplementary Table 3**). Although some of these tumors had DBS2 and ID3 signatures, we excluded the 36 samples lacking signature SBS4 and SBS92 from the subsequent analyses to ensure analyzing only samples from smokers. Thus, the reported analysis includes 309 samples with SBS4 signature, constituted of 286 lung adenocarcinomas (LUAD, 83%), 36 squamous cell carcinomas (LUSC, 11%), and 20 other or mixed subtypes (6%; **Supplementary Fig. 1**). We conducted the analyses across all samples and separately in LUAD only.

### APOBEC mutational signatures define two lung cancer subtypes in smokers

Analysis of mutational signatures, through SigProfilerExtractor and nonnegative matrix factorization^30^, revealed two distinct groups identified by the presence (43.7%) or absence (56.3%) of APOBEC mutational signatures SBS2 and SBS13 (**Fig. 1*a***; **Supplementary Fig. 2**). The absence of APOBEC signatures can be due to technical reasons, including the commonly used 5% of all mutations’ threshold for attributing somatic mutations to a signature within an individual sample^30,36–38^. To confirm this hypothesis, we utilized P-MACD^3,19^, a specialized orthogonal computational approach that focuses purely on detecting APOBEC trinucleotide motifs while providing a minimum estimation of APOBEC mutation load and a sample specific P-value (**Supplementary Table 4**).

**Fig. 1:**
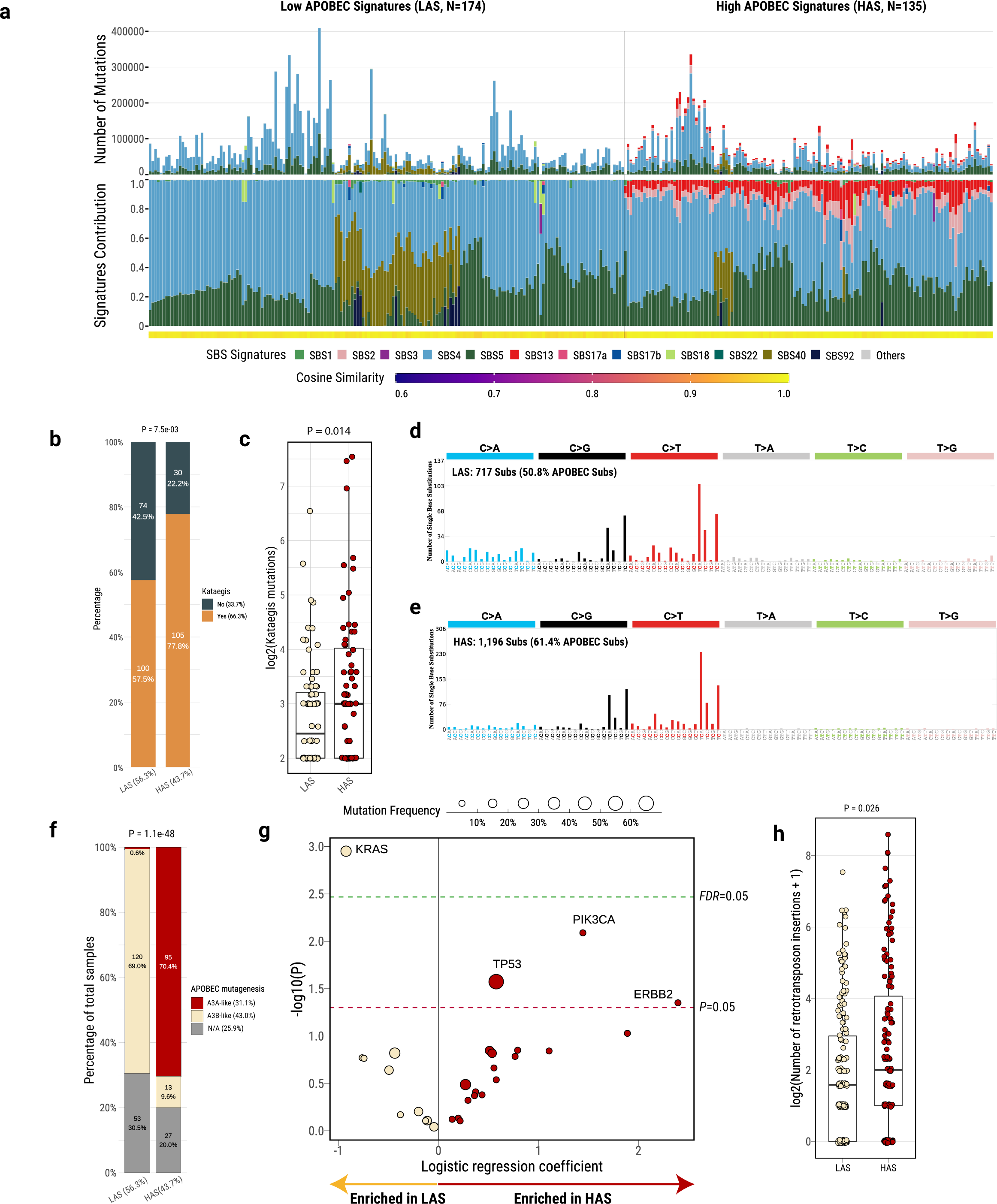
Genomic classification and characterization of lung cancer in smokers based on mutational signatures analyses. **a**, Landscape of SBS mutational processes and identification of two tumor subtypes based on APOBEC mutational signatures. The landscape of mutational signatures includes a bar plot presenting the total number of mutations assigned to each signature, the proportion of signatures assigned to each sample, and the cosine similarity between the original mutation profile and the signature decomposition. **b**, Comparisons of *kataegis* frequency between LAS and HAS tumors. **c,** Number of mutations in *kataegis* between LAS and HAS tumors. **d,e,** Mutational spectrum of total mutations contributing to *kataegis* in LAS and HAS tumors, respectively. **f**, Proportions of A3A-like and A3B-like mutagenesis between LAS and HAS tumors. Tumors not enriched with T<u>C</u>A mutations or without significant differences between RT<u>C</u>A and YT<u>C</u>A mutations are classified as N/A. **g**, Logistic regression analysis between tumor subtypes and nonsynonymous mutation status of driver genes, adjusting for the following covariates: age, sex, histology, TMB, and tumor purity. The significance thresholds *P*<0.05 (red) and *FDR*<0.05 (green) are indicated by the dashed lines. **h**, Number of retrotransposon insertions between LAS and HAS tumors.

Among the “absent” APOBEC signature group, P-MACD identified as APOBEC-positive 158 samples with significant enrichment of APOBEC mutations and 16 samples lacking an enrichment of the APOBEC trinucleotide mutational motifs (**Extended Data Fig. 1**). In addition, P-MACD confirmed all 135 samples with APOBEC signatures identified by the SigProfilerExtractor algorithm (Pearson correlation, R=1; P=0). This suggests that APOBEC mutagenesis operated in all groups, however at different levels. As a further confirmation, excluding the 16 samples with no APOBEC mutations identified by P-MACD did not substantially change the results (data not shown). Moreover, *kataegis* (clusters of localized hypermutation in cancer genomes) or C- or G-coordinated clusters were present in many samples (**Fig. 1*b*; Supplementary Table 4**), including in samples with no detected APOBEC mutations by both methods. Thus, we named the tumors with APOBEC mutations detected by P-MACD or not detected by any method as “*L*ow *A*POBEC *S*ubtype” (LAS) and those identified by both methods as “*H*igh *A*POBEC *S*ubtype” (HAS). Compared to LAS, HAS tumors showed a significantly higher frequency of *kataegis* (P=7.5e-03; **Fig. 1*b***), overall number of mutations within *kataegis* (P=0.014; **Fig. 1*c***), as well as a higher proportion of APOBEC mutations contributing to *kataegis* (61.4% *versus* 50.8%; **Fig. 1*d*-*e***), even in clusters of very low numbers of mutations (41.7%). Importantly, HAS tumors were dominated (70.4%) by the A3A-like mutator phenotype^19^, whereas LAS tumors were dominated (69%) by the A3B-like phenotype (P=1.1e-48; **Fig. 1*f***).

No significant differences were observed between LAS and HAS in terms of stage, histology, sex, tobacco smoking phenotype variables, germline APOBEC3B deletion, tumor purity, TMB, percentage of genome altered (PGA) by copy numbers, number of structural variations (SV), or SBS4 mutation burden (**Supplementary Fig. 3**). The frequency of nonsynonymous mutations in *TP53* (P=0.03; **Fig. 1*g*; Extended Data Fig. 2a**) and a few other driver genes (P<0.05; *e.g.*, *PIK3CA* and *ERBB2*; **Fig. 1*g***; **Supplementary Fig. 4**) is higher in HAS compared to LAS (P=7.28e-03, 3.21e-09, and 4.28e-05 for *TP53*, *PIK3CA*, and *ERBB2*, respectively). We found very similar results when we restricted the cancer driver gene nonsynonymous mutations to those defined as drivers based on the Cancer Genome Interpreter platform (**Methods; Supplementary Fig. 5**). As expected for APOBEC-related mutations, we found a significant enrichment of hotspot C>X mutations (X=any base) at Tp<u>C</u> sites in these genes. Unlike HAS tumors, LAS were enriched with *KRAS* driver mutations (P=1.1e-03; **Fig. 1*h*; Extended Data Fig. 2b**). We confirmed the same findings with *TP53* (P=0.041) and *KRAS* (P=0.00042) enrichments in HAS and LAS tumors, respectively, when we restricted the analyses to LUAD only (**Extended Data Fig. 2c-d**). We further validated these findings in The Cancer Genome Atlas (TCGA) LUAD dataset and observed consistent results with strong enrichment of *TP53* in HAS (P=4.67e-04) and *KRAS* in LAS (P=8.04e-03; **Extended Data Fig. 3**) tumors. In addition, HAS tumors exhibited a significantly higher number of retrotransposon insertions (P=0.026; **Fig. 1*h***), as expected for a tumor subtype with higher genomic instability^39^. In contrast, we found no mutation difference in *SMUG1* and *REV1*, which have been shown to be directly linked to the generation of APOBEC3-mediated mutational signatures^40^.

### Association between *APOBEC3B* and *UNG* expression

Among all AID/APOBEC genes, RNA-seq analyses showed that only *APOBEC3A* and *APOBEC3B* had significantly higher expression in HAS compared to LAS (P=1.95e-03 and 2.27e-04, respectively; **Fig. 2*a*; Supplementary Table 5**). As expected, *APOBEC3A* and *APOBEC3B* did not differ in normal tissue. Notably, only *APOBEC3A* expression was significantly associated with APOBEC mutation burden in HAS tumors, whereas only *APOBEC3B* expression was significantly associated with APOBEC mutations in LAS tumors (R=0.24 and 0.46; P=0.028 and 3.6e-06, respectively; **Extended Data Fig. 4**).

**Fig. 2:**
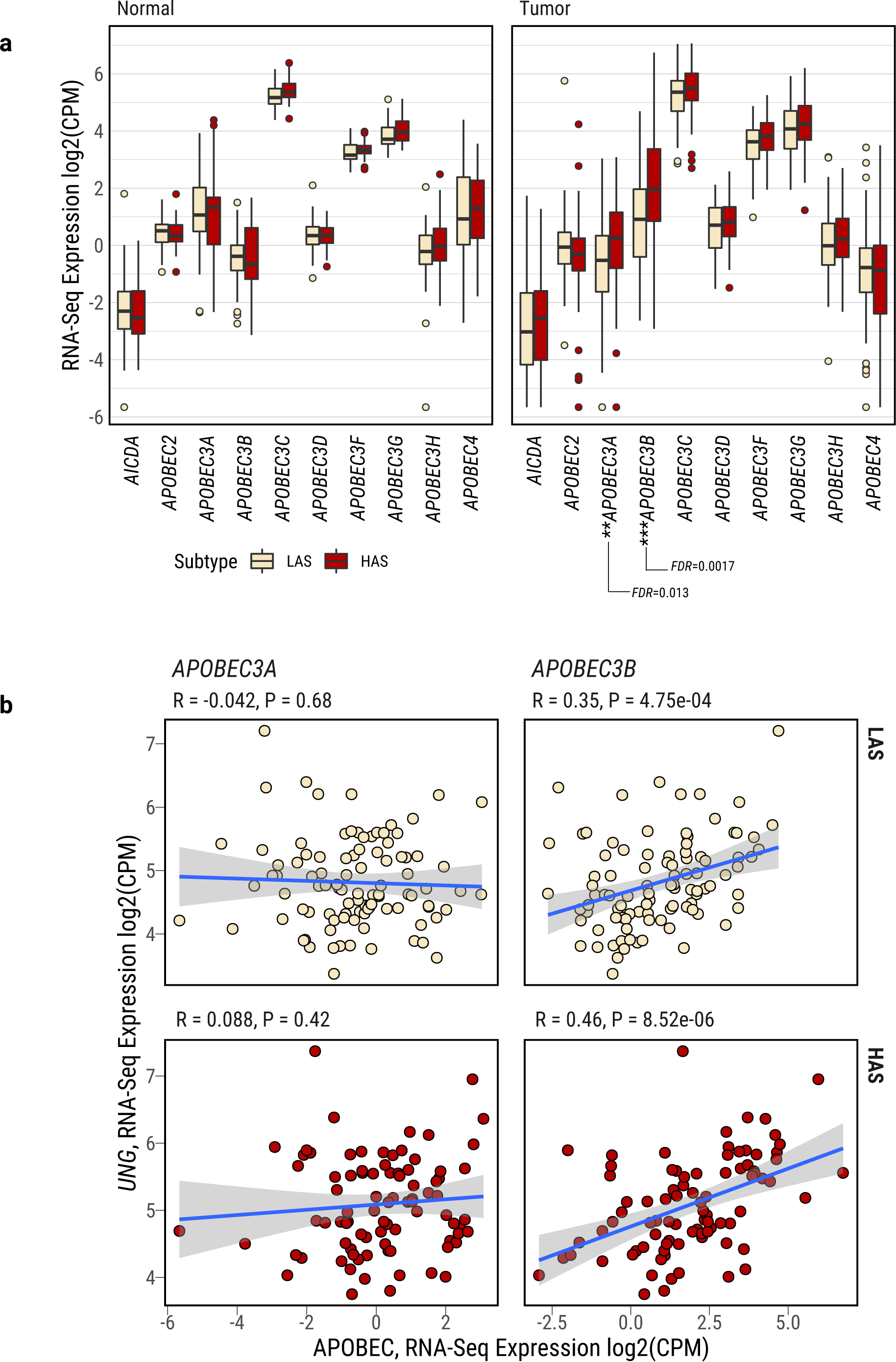
Characterization of *APOBEC3A* and *APOBEC3B* expression in lung cancers from smokers. **a**, Differentially expressed APOBEC family genes between LAS and HAS in both normal and tumor samples. After multiple testing corrections based on the Benjamini–Hochberg method, only *APOBEC3A* and *APOBEC3B* show significant differential expression between LAS and HAS tumors. Of note, *APOBEC1* expression was extremely low across most tumor samples, thus it is not included in the analysis. **b**, Gene expression correlation between *UNG* and *APOBEC3A* (left) or *APOBEC3B* (right), stratified by LAS (top) and HAS (bottom) tumors. Significant P-values and Pearson correlation coefficients are shown on top of each scatter plot.

APOBEC-catalyzed C-to-U lesions can be repaired by base excision repair (BER)^41,42^, primarily by uracil N-DNA glycosylase (UNG, also called UNG2 for the nuclear enzyme). Thus, we investigated the relationship between *UNG* and *APOBEC3A/B* gene expression. *UNG* gene expression was significantly upregulated in HAS compared to LAS in all samples (P=0.0086; **Supplementary Fig. 6a**) and in samples that were copy number neutral at *UNG* genomic location (P=0.02; **Supplementary Fig. 6b**). A significant and positive correlation of gene expression was found between *UNG* and *APOBEC3B* in both tumor subtypes (LAS: R=0.33 and P=8.8e-04; HAS: R=0.45 and P=1.5e-05), but not between *UNG* and *APOBEC3A* (LAS: R=-0.094 and P=0.36; HAS: R=0.046 and P=0.67; **Fig. 2b**). No association was observed between *UNG* and *APOBEC3A* or *APOBEC3B* in normal tissue (**Supplementary Fig. 7a**). Additional genes involved in the BER pathway exhibited a similar pattern, with positive association with *APOBEC3B* but not *APOBEC3A* in tumor samples (**Supplementary Fig. 8**). This finding suggests that *UNG* expression is activated by uracils in DNA generated by APOBEC3B mutagenesis or that *UNG* and *APOBEC3B* share common regulatory mechanisms^43,44^. We observed a similar pattern with *UNG* positively associated with *APOBEC3B*, but not *APOBEC3A* in TCGA LUAD (**Supplementary Fig. 7b-c**) as well as in multiple cancer types in TCGA (*e.g.*, prostate adenocarcinoma; head and neck squamous cell carcinoma; pancreatic ductal adenocarcinoma; lymphoid neoplasm diffuse large B-cell lymphoma; lung squamous cell carcinoma; **Extended Data Fig. 5**). We note that 91.8% of APOBEC mutations were clonal in HAS tumors, although APOBEC mutations were higher than the subclonal mutations due to other mutational processes (**Supplementary Fig. 9a-b**). Similarly, we found that, although the ratio of APOBEC mutations over all mutations was higher for subclonal mutations than clonal mutations, most APOBEC mutations are clonal across different cancer types in the PCAWG study^45^ (**Supplementary Fig. 9c**). The clonality of APOBEC mutations do not support frequent episodic APOBEC mutagenesis as the likely explanation of the lack of association between *APOBEC3A* and *UNG*/BER expression. Moreover, it is unlikely that chronologically discrepant episodic APOBEC3A mutagenesis and UNG/BER induction show similar findings across all cancer types and multiple BER enzymes.

### Tobacco smoking addiction is associated with genomic changes only in LAS tumors

Mutational signature analysis showed that the known tobacco smoking-associated signatures (SBS4, DBS2, and ID3) are positively associated with each other and with the APOBEC signature as well as with other mutational signatures and the overall TMB (**Supplementary Fig. 10**) as previously shown^29^. Among the five tobacco smoking phenotype variables available in 198 subjects from the Environment and Genetics in Lung cancer Etiology (EAGLE) study^35^ (**Supplementary Fig. 11**), we previously found that “time to first cigarette” (TTFC) in the morning, a marker of strong nicotine addiction and high tobacco smoking exposure^46^, showed the only significant association with TMB and tobacco smoking signatures^47^, possibly because of a decreased DNA repair function in those who wake up early to smoke ^48^. However, when we separated the tumors between LAS and HAS in this study, short TTFC was associated with an increased TMB (P_trend_=0.0024, **Fig. 3*a*** and P_trend_=0.0054 in LUAD only, **Supplementary Fig. 12**)**; Methods**) and the other tobacco smoking signatures (SBS4, P_trend_=0.005; DBS2, P_trend_=0.0044; and ID3, P_trend_=0.0029; **Extended Data Fig. 6**) only in LAS samples. Moreover, only LAS tumors showed a significant association between short TTFC and increased mutational burden across different mutation types (**Supplementary Fig. 13**) and in frequently mutated genes known to be associated with tobacco smoking^49^ (*e.g.*, *ZFHX4*; **Extended Data Fig. 7**). Surprisingly, the frequency of genetic variants associated with nicotine addiction^50^ did not vary between HAS and LAS tumors (**Supplementary Fig. 14**), thus this genetic variation does not appear to be the cause of LAS/HAS difference. A more plausible reason for this finding is the different cell composition between HAS and LAS tumors. Strong DNA damage induced by APOBEC3A hypermutation and *TP53*-associated genomic instability can cause more cell senescence and apoptosis^51–54^ in HAS tumors. Apoptosis and senescence can in turn lead to cell regeneration^55,56^. HAS tumor composition likely includes a high number of newly generated or de-differentiated cells, which do not display the expected tobacco smoking-associated patterns that are evident in the more differentiated LAS tumors.

**Fig. 3:**
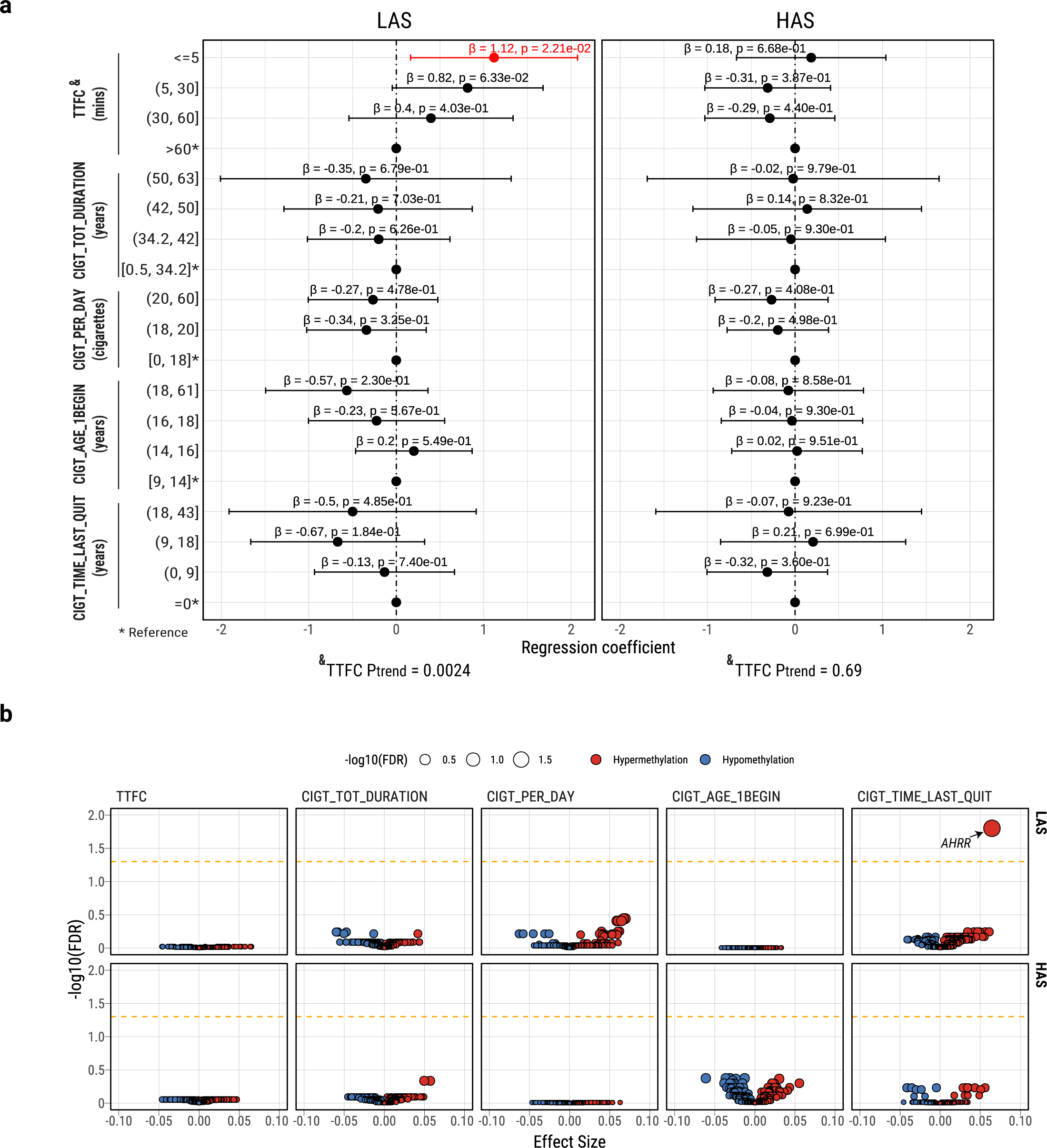
Multivariate regression analysis between five tobacco smoking variables and genomic (n=198) or epigenomic (n=122) features in the EAGLE samples. **a**, Forest plot for the associations between TMB and smoking variables, stratified between LAS and HAS tumors. P-values and regression coefficients with 95% confidence intervals (CIs) are shown for each category of smoking variables. Significant associations are in red ink. Trend test P values (P_trend_) from associations between TTFC and TMB are included below the forest plots. **b,** Volcano plots of the associations between smoking variables and methylation levels of known smoking-related CpG probes in tumors. Association FDR values (adjusted using the Benjamini-Hochberg method) are shown on the y-axis. The orange dashed line indicates the associations with FDR<0.05. The CpG probes associated with tobacco smoking are derived from a study^65^ comparing methylation levels between smokers and never smokers in normal lung tissue. The size and color of each point represent the FDR and association direction, respectively. All association analyses are adjusted for the following covariates: age, sex, histology, and tumor purity.

### Tobacco smoking-induced epigenomic changes can be reversed after quitting smoking only in LAS tumors

Smoking exposure has been shown to induce DNA hypomethylation in the *AHRR* gene^57^. *AHRR* hypomethylation has been proposed to be a marker of tobacco smoking status^58^ and lung cancer mortality^59^; whereas increased methylation at this locus has been used as a marker of successful smoking cessation^60–62^. Altered methylation levels have been seen in other genes following tobacco smoking exposure, although to lesser degrees^63–65^. To understand whether hypermutation-induced cell regeneration in HAS also alters the association between smoking behaviors and DNA methylation changes, we tested associations between smoking variables and methylation levels at known smoking-associated CpG probes (reviewed recently in Ref.^65^) in both tumor and normal datasets of HAS and LAS (**Supplementary Table 6**). As expected, cg05575921 (*AHRR*) was hypomethylated in current smokers *versus* former smokers and according to years from quitting smoking (**Supplementary Fig. 15**). Notably, in normal lung tissue, ‘time since last quitting smoking’ and ‘total smoking duration’ were associated with altered methylation levels at 25 sites (16 genes) and 8 sites (5 genes), respectively (**Extended Data Fig. 8a**). As expected in normal tissue, ‘time since last quitting smoking’ was the only variable associated with any CpG probes, particularly in cg14120703 (*NOTCH1*), cg05575921 (*AHRR*), and cg10420527 (*LRP5*), when both ‘time since quitting’ and ‘smoking duration’ were incllineuded in the same regression model (**Extended Data Fig. 8b**). Importantly, in tumor samples, the association between cg05575921 (*AHRR*) and time since last quitting smoking is observed only in LAS after adjusting for age, sex, tumor histology and tumor purity (P=3.8e-05, **Fig. 3*b*; Supplementary Fig. 15**; P=5.81e-05 in LUAD only). In contrast, *NOTCH1* (**Supplementary Fig. 16**) and other genes (**Supplementary Table 7**) whose methylation levels are not associated with tobacco smoking in tumors showed no difference between LAS and HAS. These findings further support the hypothesis that the dynamic cell composition in HAS tumors can indirectly disrupt the reversion of methylation levels in cg05575921 (*AHRR*) following smoking cessation. We confirmed these results in TCGA LUAD samples (**Supplementary Fig. 17**).

### Cell senescence followed by cell regeneration in HAS tumors

To further confirm the differences in cell composition between HAS and LAS tumors, as a consequence of the enrichment of A3A-like mutagenicity and *TP53*-induced genomic instability in HAS, we analyzed RNA-seq data reflecting the transcriptomic landscape at the time of diagnosis. We tested which gene expression pathways were associated with TTFC in the two APOBEC-based subtypes. The most significantly upregulated pathway associated with short TTFC in HAS tumors was the pulmonary healing signaling pathway (Z-score=3.78; P=3.34e-04; **Supplementary Fig. 18**; **Methods**), which is related to a dynamic process of regeneration of alveolar cells from stem cells. The expression of this pathway was significantly higher in HAS than LAS tumors (Z-score=1.57; P=1.06e-03). These findings suggest that HAS tumors at time of diagnosis have a higher frequency of quiescent stem cells that have exited the quiescent state. In fact, we found a higher gene expression of basal cell markers (*e.g., KRT19, KRT15* and *TP63)*, a sign of lineage infidelity^66^ or squamous differentiation in HAS tumors (**Fig. 4*a***). In contrast, LAS tumors have a higher expression of alveolar type II (AT2) cell markers (*e.g*., *NKX2-1*, *SFTPB,* and *NAPSA),* which are expected in differentiated cells of the alveolar epithelia^67^. These findings were confirmed after adjustment for copy number alterations and tumor purity (**Supplementary Table 8**). We also inferred the cell-of-origin by correlating somatic mutational density with single cell-based lung expression profiles of benign epithelial cell types (**Methods**). Consistent with the RNA-seq results, we observed proximal lung cell types (Club/Basal/Ciliated cells) enriched in HAS and distal lung cell types (AT2) enriched in LAS (P=0.028; OR=2.21; **Supplementary Fig. 19**).

**Fig. 4:**
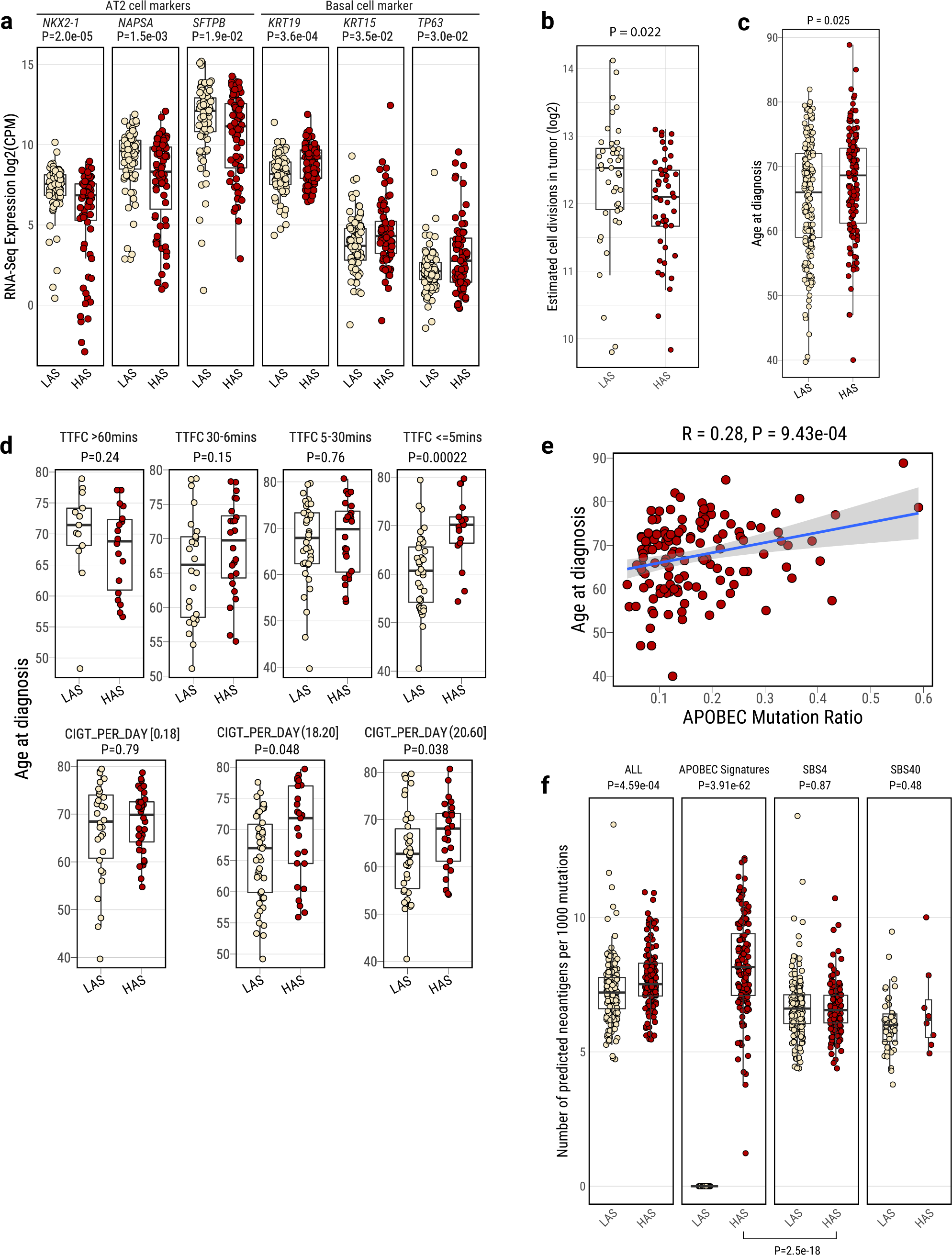
Tumor cell composition and age at onset differences between LAS and HAS tumors. **a**, Boxplots show the differentially expressed gene markers of lung-specific cell types between LAS and HAS LUAD tumors (n=155). **b,** Cumulative number of stem cell division estimates in LAS and HAS tumors based on methylation data. **c**,**d**, Age at diagnosis difference between LAS and HAS tumors overall (**c**), and (**d**) stratified by TTFC [Time to first cigarette in the morning (from the first question of the Fagerstrom test for nicotine dependence: ‘How soon after you wake up do you smoke your first cigarette?’)] or CIGT_PER_DAY (Average intensity of cigarette smoking, measured as the number of cigarettes per day). **e,** Correlation between APOBEC mutation ratio and age at diagnosis in HAS tumor. **f**, Neoantigen prediction for different mutational signatures between LAS and HAS. P-values from Wilcoxon rank-sum tests are labeled for each boxplot. On the bottom, P-value for the different contribution of SBS4 and APOBEC mutational signatures to neoantigen prediction in HAS tumors.

### APOBEC influences age at onset and neoantigens presentation

We hypothesized that recently generated cells in HAS tumors require a long time to accumulate mutations before the tumors become clinically evident. Consistent with this hypothesis, the estimation of methylation-based mitotic rate in HAS tumors showed lower cell division than LAS tumors (P=0.022; **Fig. 4*b***). Importantly, patients with HAS tumors showed a later age at diagnosis (univariate test P=0.025; P=0.022 after adjusting for histology, stage, and sex; **Fig. 4*c***) in comparison to LAS cancers. This difference between LAS and HAS tumors was strongly affected by smoking exposure levels, with LAS showing ∼10 years and ∼5 years younger age at onset in subjects with shorter TTFC (<=5 mins; univariate test P=2.2e-04 and P=0.0034 for LUAD only; multivariate test P=5.91e-04 and P=3.14e-03 for LUAD) or a high number of cigarettes per day (>20; univariate test P=0.038 and P=0.022 for LUAD only; multivariate test P=0.075 and P=0.092 for LUAD), respectively (**Fig. 4*d***). The proportion of APOBEC mutations was positively associated with age at diagnosis in HAS tumors (R=0.28; P=9.43e-04; **Fig. 4*e*;** R=0.39; P=4.21e-05 in LUAD only). We found a later age at onset of the most recent common ancestor (MRCA; **Supplementary Fig. 20a-b**) and no significant difference between HAS and LAS in terms of latency, measured as the difference between the age at diagnosis and the estimated patients’ age at the appearance of the MRCA (**Supplementary Fig. 20c-d**). These findings suggest that APOBEC mutagenesis has a stronger effect on the tumor clonal expansion, precisely when tobacco smoking mutagenesis has its strongest effect^45^, than on the subsequent tumor progression. In contrast, age at diagnosis was not associated with mutation status of major cancer driver genes (i.e., *KRAS* and *TP53*, **Supplementary Fig. 21**).

Notably, the accumulated mutations during HAS evolution exhibited a higher propensity to generate neoantigens than in LAS, adjusting for the non-synonymous mutation burden (P=0.00046; **Fig. 4*f***; P=3.12e-13, **Supplementary Fig. 22** in TCGA LUAD). Within HAS, APOBEC mutations generated more neoantigens than tobacco smoking (P=2.5e-18). The neoantigens in HAS tumors likely increased tumor immune-editing during tumor development^68^, potentially contributing to their late age at diagnosis. Intriguingly, HAS tumors also showed a higher *PD-L1* expression (P=0.020, **Supplementary Fig. 23a**, and P=0.0095 in TCGA HAS LUAD, **Supplementary Fig. 23b**) than LAS tumors. The high neoantigens and *PD-L1* expression suggest that HAS tumors not only are diagnosed later than LAS, but they may also respond better to immunotherapy^13^, a treatment option that could compensate for the APOBEC3A-related resistance to targeted treatments ^69^. Of note, HAS tumors also showed a not-significant improved survival than LAS tumors even though no patients in this study received immune-related treatments (**Supplementary Fig. 24**).

We verified whether an association between APOBEC signature subtypes and age at diagnosis is present across different cancer types in the PCAWG study, taking advantage of the WGS data that allows a precise classification of HAS and LAS subtypes even in tumor types with low mutational burden. Notably, the strongest difference was found in Skin-Melanoma (10 years, P=0.0094), a tumor type associated with a strong exogenous mutagenic exposure (UV radiation), like lung cancer (**Supplementary Fig. 25a**). Since the UVR signatures SBS7a/b have an overlapping context with SBS2, we repeated the analyses separating melanoma HAS and LAS subtypes only using SBS13 and confirmed the results (**Supplementary Fig. 25b**). Thus, while clock-like SBS1 and SBS5 signatures have been shown to increase with age across different tissue types^4,70^, our findings suggest that high APOBEC mutagenesis indirectly influences tumor age at onset in the presence of other strong exogenous mutagenic processes. Of note, we observed no genomic or age at diagnosis differences when we stratified tumors between the presence or absence of other mutational signatures (*e.g.*, SBS40; **Supplementary Fig. 26**).

## DISCUSSION

Our large multi-omics study shows that the co-occurrence of APOBEC and tobacco smoking mutagenesis shapes lung tumor evolution and age at onset of lung cancer from smokers (**Fig. 5**).

**Fig. 5.**
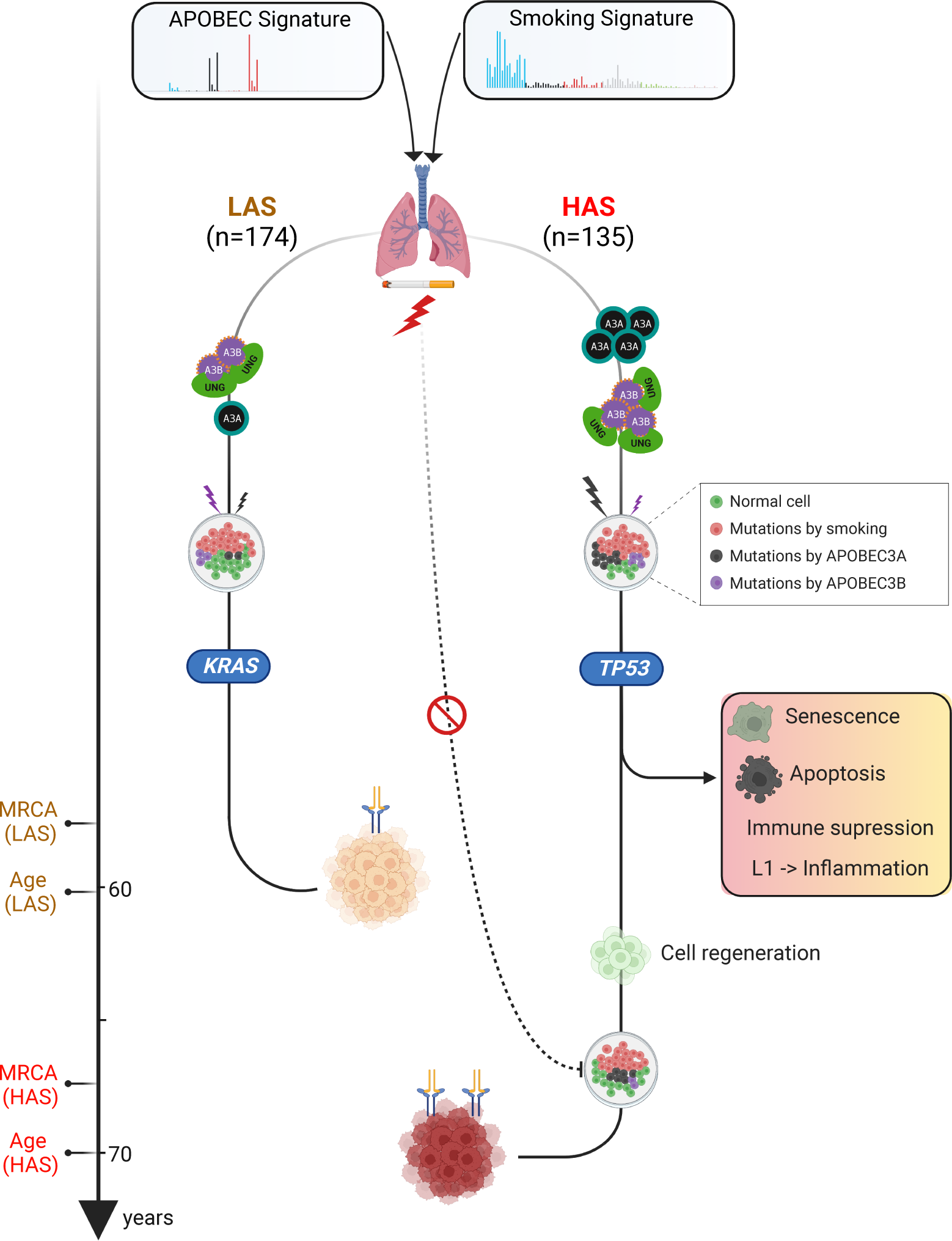
Conceptual diagram of APOBEC shaping tumor development and influencing age at onset of lung cancers from smokers. The schematic was generated using BioRender (https://biorender.com/).

Two distinct tumor subtypes are defined by APOBEC mutagenesis. One is characterized by APOBEC signatures SBS2/13 accounting for more than 5% of mutations in each sample and strong A3A-like mutagenesis (HAS). The other has a lower level of APOBEC mutagenic activity (<5%) and a dominant A3B-like phenotype (LAS). Both *APOBEC3A* and *APOBEC3B* expressions are highest in HAS. Intriguingly, HAS and LAS tumors are enriched with *TP53* and *KRAS* alterations, respectively.

Episodic APOBEC mutagenesis can be enhanced by ssDNA APOBEC substrate formation in response to DNA damage, such as that caused by tobacco carcinogens^71^. A3A mutagenesis, higher in HAS tumors, is known to be stronger than A3B^6,19,72^. Unlike *APOBEC3B*, *APOBEC3A* expression is not associated with mRNA expression of *UNG* and other BER genes in our study, suggesting that A3A mutations are less likely to be repaired by UNG. APOBEC3A has been reported to activate DNA damage response and cause cell-cycle arrest^73^. We hypothesize that the high DNA damage due to unrepaired tobacco smoking and high A3A mutagenesis can trigger cell senescence in HAS tumors. In the presence of *TP53* alterations (enriched in HAS tumors), senescence can be bypassed. Consequently, cells can undergo a crisis and can die through spontaneous mitotic arrest^74^. Furthermore, during cellular senescence, LINE-1 retrotransposable elements (enriched in HAS) have been shown to become transcriptionally derepressed and activate a type-I interferon (IFN-I) response, which in turn contributes to sterile inflammation, a hallmark of aging^75^. Cell senescence^7^, apoptosis^8^ and inflammation^9^ contribute to cell regeneration. Consistent with these observations, the pulmonary healing signaling pathway associated with cell regeneration showed higher expression in HAS *versus* LAS, particularly in heavy smokers. Moreover, HAS tumors had higher markers of lineage infidelity and stemness than LAS. In contrast, the expression of typical LUAD AT2 cells and distal cell-of-origin were higher in LAS tumors, which were enriched with *KRAS* mutations. AT2 cells are the likely cells of origin of lung adenocarcinomas harboring *KRAS* mutations^76^.

Importantly, the dynamic cellular state and composition in HAS could explain why the expected associations of tobacco smoking exposure with genomic or with epigenomic changes are not observed in this subtype. In fact, LAS and HAS tumors have similar overall mutational burdens, copy number alterations and structural variants, although they strongly differ between A3A and A3B mutagenesis and TP53-related genomic instability. Intriguingly, studies on human bronchial epithelial cells showed that tobacco smoking dose-dependent mutation frequency reaches a plateau at around 23 pack-years, followed by a similar mutational burden between heavy smokers and light smokers^77^, possibly a consequence of frequent cell senescence/apoptosis in tumors from heavy smokers.

Understanding how heterogeneity in mutational burden due to co-occuring mutational processes and cell types contributes to clonal evolution is important to understand lung carcinogenesis. Regenerated cells following senescence or apoptosis in HAS are likely to require a long time to accumulate mutations and to undergo subsequent clonal expansion. In addition, hypermutation and related neoantigens can trigger an immune response in the tumor microenvironment, slowing the tumor growth^78,79^. In contrast, LAS tumors, with their lower APOBEC-related mutational burdens, fewer neoantigens, and infrequent LINE 1 retrotransposons, are less likely to undergo cell senescence or apoptosis or to induce a strong immune response. Consequently, they can more steadily accumulate tobacco smoking induced mutations in the progenitor cells, increasing the probability of earlier clonal expansions. This could explain why LAS tumors have an earlier age at diagnosis than HAS tumors, particularly in the presence of strong tobacco smoking exposure, as indicated by the short time to first cigarette in the morning or a high number of cigarettes per day. We observed a similar pattern in melanoma, which is known to be associated with high UV exposure.

In conclusion, APOBEC appears to influence the effect of other mutagenic activities on the cancer genome, with important clinical implications. Currently, investigations into possible cancer treatment strategies inhibiting APOBEC3 mutagenesis are ongoing^40^. Our findings suggest that the mutagenic contexts in which APOBEC3 operates as well as the specific type of APOBEC mutagenesis need to be considered in developing effective treatments.

Finally, this study emphasizes the importance of collecting detailed exposure information and WGS data in cancer studies to better understand the interplay between exogenous and endogenous activities in tumor development.

## METHODS

### Data sources

We originally collected multi-omics datasets from a total of 345 smokers with pathologically confirmed and treatment-naïve lung primary cancers, including 345 tumors with WGS, 196 tumors with mRNA-seq, and 206 tumors with DNA methylation. Among these tumors, 221 were from the EAGLE study^35^, with 218 of these subjects having detailed clinical and smoking information. Briefly, EAGLE tumor samples were histologically confirmed as primary lung cancers with at least 50% tumor nuclei and less than 20% necrosis. Samples were snap-frozen in liquid nitrogen within 20 min of surgical resection and the precise site of tissue sampling was recorded. Detailed information on tumor characteristics, recurrence, treatment, and follow-up data were recovered from patients’ medical records and follow-up visits and hospital admissions were identified by linkage with the region-wide Regional Health Authority database. The study protocol was approved by the Institutional Review Board of the US National Cancer Institute and the involved institutions in Italy. Informed consent was obtained for all subjects prior to study participation. The remaining 124 smokers originated from multiple cancer genomic studies^80–86^ (**Supplementary Table 1**). To investigate the interplay of APOBEC and tobacco smoking mutagenesis, we only included 309 tumors with detected SBS4 signatures in this analysis. The following WGS data inclusion criteria were used for sample selection before downstream somatic analyses: 1) average sequencing depth in tumor samples >40x and normal samples >25x; 2) sample contamination rate <1% measured by both Conpair (v.0.2)^87^ and Somailer (v.0.2.6)^88^; 3) no obvious large copy number detected in normal samples by the Battenberg algorithm (suggesting no tumor and normal sample swapping^66^); 4) tumor exclusions if signature SBS7 (exposure to ultraviolet light, potential metastatic melanoma) or SBS31 (prior chemotherapy treatment with platinum drugs) are identified by mutational signature analysis; 5) Exclusion of samples with low-quality whole genome sequencing data if the number of detected somatic alterations <1000 and NRPCC (number of reads per tumor chromosomal copy, representing the sequencing coverage per haploid genome) <10, or if the number of somatic alterations <100.

### Whole genome sequencing and data preprocessing

Frozen tumor samples and matched normal controls from 221 smokers with lung cancer from the EAGLE study^35^ were selected for whole genome sequencing. DNA extraction and sequencing library construction followed our previously reported protocol^66^. Illumina HiSeq X was used for performing WGS at the Broad Institute following standard Illumina protocols with targeted sequencing depth 80x for tumor and 30x for normal tissue samples. The raw sequencing data in FASTQ format were collected and the GATK best practices workflow was applied to generate the analysis-ready CRAM files using GRCh38 as the human reference genome. For the WGS data from public databases, the preprocessed aligned BAM/CRAM files (including unmapped reads) were first converted back to FASTQ files using Bazam (v.1.0.1)^89^ to retain the sequencing lane and read group information and then processed using the same pipeline as for the EAGLE dataset.

### Genome-wide somatic alteration calling

Genome-wide somatic mutations were called from analysis-ready CRAM files following our previous bioinformatic pipelines^66^. Four different calling algorithms were applied using tumor-normal paired analysis, including Strelka (v.2.9.10), MuTect, MuTect2, and TNscope. Sentieon’s genomics software (v202010.01) was used to run the MuTect (named TNsnv), MuTect2 (named TNhaplotyper), and TNscope algorithms. An ensemble method was applied to merge different callers with additional filtering to reduce false positive calling^66^. The final mutation calls for both SNVs and indels needed to meet the following criteria: 1) read depth >12 in tumor and >6 in normal samples; 2) variant allele frequency <0.02 in normal samples, and 3) overall allele frequency (AF) <0.001 in multiple genetics databases including 1000 Genomes (phase 3 v.5), ExAC (v.0.3.1), and gnomAD exomes (v2.1.1) and genomes (v3.0). For indel calling, only variants called by at least three algorithms were kept, followed by left normalization.

For somatic copy number alterations (SCNAs), the Battenberg algorithm (v.2.2.9)^90^ was used to generate the initial SCNA profile followed by assessment of clonality of each segmentation, as well as tumor purity and tumor ploidy. Battenberg refitting was repeated with a suggested tumor purity and ploidy as input until the final SCNA profile was retained (see below). Recurrent copy number alterations at the gene level were identified using GISTIC v.2.0^91^ based on the major clonal copy number for each segmentation. Two algorithms, Meerkat (v.0.189)^92^ and Manta (v.1.6.0)^93^, were applied with recommended filtering for identifying structural variants (SVs) and the union set of these two callers was merged as the final SV dataset. A window of 50 bp was used when merging SVs. For the identification of putative transposon elements (TE), we used a pipeline called TraFiC-mem (v.1.1.0) (Transposome Finder in Cancer)^94^ and only the TE insertions that passed the default filtering were kept. We define the overall somatic alterations as the summary of SNVs, indels, SVs, and TE insertions.

### Consensus estimation of tumor purity, ploidy, and clonality

For a consensus estimation of tumor purity, we systematically compared three approaches based on different genomic features as previously reported^66^: SCNA profiling using Battenberg; mutation clustering (VAF/CCF distribution) using Bayesian Dirichlet Process (DPClust v.2.2.8)^90,95^; and bayesian mixture models based on both SCNA and mutation clustering using CCUBE^96^. First, the SCNA calling was performed at the sample level based on the Battenberg algorithm followed by the DPClust process. Overall genomic matrices were generated including the purity estimation consistency among Battenberg, DPClust, and CCUBE; ploidy status; superclones or superclusters (CCF>1); percentage of genomic alterations by SCNA (PGA); homozygous deletion size; percentage of segmentation with loss of heterozygosity; etc. Unsupervised hierarchical clustering was performed based on this matrix, which allows the identification of different groups of tumors with similar genomic profiling features and to flag samples for refitting (*e.g.*, good profile, swapped diploid and tetraploid status, over/underestimated purity, incorrect segmentation for copy number alterations). Battenberg was then used to calculate new purity and ploidy values for refitting. This process was repeated on flagged tumor samples until a good genomic profile was retained with consensus tumor purity estimation.

### Mutational signature analysis and identification of the APOBEC mutator phenotype

SigProfilerMatrixGenerator^97^ was used to create the mutational matrix for all types of somatic mutations. The deconvolution of mutational signatures was performed by SigProfilerExtractor (v.1.1.3)^30^ using the WGS setting and the COSMIC mutational signatures as reference (v3.2). Hierarchical clustering of the contribution of mutational signatures was performed using Euclidean distance and Ward’s minimum-variance clustering separated by the presence of signature SBS4 and APOBEC signatures SBS2 and SBS13. SigProfilerClusters (v.1.0.0)^98^ was used to identify *kataegis* and other clustered mutation events (*e.g. Omikli*). Mutational signature analyses were repeated based on the clustered mutations.

To further verify the APOBEC mutational loads, we also analyzed the pattern of mutagenesis by APOBEC cytidine deaminase using P-MACD^3,19^. A mutation annotation file (MAF) was used as input and sample-specific summaries were generated, including “APOBEC_MutLoad_MinEstimate”, which were compared against the number of APOBEC mutations (SBS2+SBS13) estimated by SigProfileExtractor. In addition, we estimated the percentage of hypermutable strand-coordinated ssDNA (scssDNA) per genome, which can represent events of break-induced replication occurring at positions of double-strand breaks that cannot be repaired by canonical DSB repair^99^. Mutation clusters from P-MACD analysis with P≤10^−4^ were considered *bona fide* mutation clusters. Mutation clusters with <10 bp inter-mutational distances were considered as complex events and counted as one mutation, which originated from a single mutagenic event rather than from independent DNA lesions. Mutation clusters with more than 3 mutations containing only mutated Cs or only mutated Gs were used as a proxy for the long persistent regions of scssDNA. These clusters are associated with the enrichment of APOBEC mutation signatures in cancer^3,23,100,101^. The length of a hypermutable ssDNA region was estimated as the distance between mutations bordering a “C” or “G” coordinated cluster.

To distinguish the APOBEC hypermutation signature from APOBEC3B mutagenesis, we followed a previous procedure to calculate Y/RTCA enrichment and identify A3A-like or A3B-like tumors^19^. Given the null hypothesis of random mutagenesis, the expected number of YT<u>C</u>A mutations was calculated based on the fraction of motifs at YT<u>C</u>A from mutations within the T<u>C</u>A context. The expected number of RT<u>C</u>A and NT<u>C</u>A mutations (N=any base) were computed analogously. The χ2 test for goodness of fit was used to identify samples that had a ratio of YT<u>C</u>A to RT<u>C</u>A mutations that differed statistically from random, by comparing observed *vs*. expected mutation counts. P-values were corrected by the Benjamini-Hochberg method with Q<0.05 considered significant.

### Estimation of the age at onset of the most recent common ancestor (MRCA) and tumor latency

We estimated the tumor chronological expansion, including the occurrence of the most recent common ancestor (MRCA) and the tumor latency (calculated as the difference between age at tumor diagnosis and age at the occurrence of the MRCA), using an approach from our previous studies^45,66^. For this analysis, we only included 151 high quality tumors (tumor purity >0.3 and NRPCC (representing the sequencing coverage per haploid genome) >10) with age at diagnosis information. In summary, the number of clock-like CpG>TpG mutations in a NpCpG context (SBS1) was counted for all tumors and a hierarchical bayesian linear regression was fit to relate to age at diagnosis accounting for tumor ploidy, tumor purity and subclonal architecture. The age at the occurrence of the MRCA was estimated for each tumor, adopting an estimated tumor acceleration rate of 5x due to the high mutational burden as previously shown^45^. We subtracted the age at occurrence of the MRCA from the age at tumor diagnosis to estimate the tumor latency or subclonal diversification.

### Cell of origin (COO) inference from genome-wide mutational patterns

COO inference was performed using a recently developed procedure (manuscript under review). In summary, seven benign lung cell type-specific gene expression centroids were derived from the scRNA-seq lung atlas^102–105^, including basal, neuroendocrine, goblet, ciliated, club, alveolar type I and alveolar type II cells. Gamma-Poisson regression was applied to correlate each cell type-specific gene expression profile with somatic SNV profiles from our study and was adjusted for covariates, including replication timing, GC context, average gene expression across all cell-type specific centroids, log intronic fractions, and log exonic fractions. The maximum likelihood regression fit was used to obtain the main effect size (relative size) and 95% confidence interval on the relative risk. A separate regression was fitted for each cell type-specific gene expression profile. The median relative risk was used to compare the COO difference between LAS and HAS tumors. In addition, lung cell-type specific gene expression markers^67^ were used to validate these findings.

### Neoantigen prediction

To predict neoantigens, patient-specific HLA haplotypes were identified using HLA-HD (v.1.2.1)^106^. The software NetMCHpan4.1^107^ was then run on 9-11 neo peptides derived from all nonsynonymous mutations, taking into account the patient’s specific HLA genotypes. NetMCHpan4.1 was included in the NeoPredPipe pipeline (v.1.1)^108^. The decomposed mutation probabilities generated by the SigProfilerExtractor were used to predict the neoantigens derived from specific mutational signatures. For TCGA data, the predicted SNV neoantigen counts per sample were collected from a previous publication ^109^. We used the number of peptides predicted to bind with MHC proteins as the neoantigen load in our analysis.

### Genomic enrichment analyses between LAS and HAS tumors

We identified cancer driver genes using the intOGen pipeline^110^ and 30 genes with a combined q value<0.1 were selected as the driver genes for enrichment. Logistic regression was performed using these genomic alterations (nonsynonymous mutation status) as an outcome and tumor subtype as a variable, adjusting for the following covariates: age, sex, TMB, tumor purity, and histology. In addition, we further restricted the analyses to mutations predicted to be drivers based on the Cancer Genome Interpreter platform^111^ (https://www.cancergenomeinterpreter.org/home). Specifically, the definition of “driver mutation” was based on a machine-learning approach (BoostDM) and a rule-based approach (OncodriveMUT). To be defined as a “driver” the mutation had to be identified as such by both BoostDM and OncodriveMUT.

### Smoking variables association analysis

To investigate associations between genomic data and tobacco smoking variables, we evaluated five variables, including CIGT_AGE_1BEGIN = Age (in years) when subjects started smoking cigarettes regularly for the first time; CIGT_PER_DAY = Average intensity of cigarette smoking, measured as the number of cigarettes per day; CIGT_TIME_LAST_QUIT = Number of years since the subject quitted smoking cigarettes (0 means current smokers); CIGT_TOT_DURATION = Total period (in years) during which the subject smoked cigarettes regularly; TTFC = Time to first cigarette in the morning (from the first question of the Fagerstrom test for nicotine dependence: ‘How soon after you wake up do you smoke your first cigarette?’). Multivariate regression analyses were performed between these categorical smoking variables and genomic data including TMB, SBS4, DBS2, ID3, and number of mutations for each mutation subtype. Regression models were adjusted for the following covariates: age, sex, histology, and tumor purity. In addition, similar multivariate regressions (as trend testing) were performed by transforming the smoking variables from categorical variables into continuous variables.

### Methylation data analysis

Genome-wide DNA methylation was profiled on the Illumina HumanMethylationEPIC BeadChip (Illumina, San Diego, USA). Genomic DNA was extracted and DNA methylation was measured according to Illumina’s standard procedure at the Cancer Genomics Research Laboratory (CGR), National Cancer Institute. Raw DNA methylation data (“.idat” files) were generated for our study, combined with methylation data collected from the public TCGA LUAD study (Illumina Human Methylation 450k BeadChip) (**Supplementary Table 1**), and processed using RnBeads (v.2.0)^112^. Background correction (“enmix.oob”) and beta-mixture quantile normalization (“BMIQ”) were applied. Unreliable probes (Greedycut algorithm with detection P<0.05), cross-reactive probes, and probes mapping to sex chromosomes were removed. Samples with outlier intensities in 450k/EPIC array control probes were removed from the dataset as described in the RnBeads vignette. We performed principal component analysis (PCA) of these samples based on the genotyping probes and removed subjects with Euclidean distance between matched tumor/normal pair > upper quartile + 3*inter-quartile range (IQR). In addition, we performed Combat batch correction separately for tumors and normal samples using the “sva” R package^113^ on M-values for known technical factors including studies, collection sites, sample plates, sample wells, array types (450K and EPIC), chip ID, and positions. We used MBatch (v2.0, https://github.com/MD-Anderson-Bioinformatics/BatchEffectsPackage) to assess for the presence of batch effects. The Dispersion Separability Criteria (DSC) values were <0.5 for all technical factors, suggesting that batch effects were negligible after batch correction. The corrected methylation-level beta values were used for downstream association analyses.

We selected 952 probes that had genome-wide significant associations with smoking variables in a recent study^65^. The P-value and methylation status (hypermethylation vs. hypomethylation) of these probes were included in our visualization of the association results. Linear regression was performed for each smoking variable using the methylation beta value as an outcome and adjusting for the following covariates: age, sex, histology, and tumor purity for the tumor methylation analysis, and age and sex for the methylation analysis of normal tissue. Association P-values were further corrected using the Benjamini-Hochberg method. These analyses were then repeated after separating LAS and HAS tumors. In addition, multivariate regression analyses were also performed including both CIGT_TIME_LAST_QUIT and CIGT_TOT_DURATION as well as age and sex as covariates in the methylation analysis of normal tissue.

In addition, EpiTOC2^114^ were used to estimate the intrinsic stem cell division rate or mitotic clock based on the methylation status of 163 EpiTOC2 CpG probes.

### RNA-Seq data analysis

RNA-Seq data from the EAGLE samples were generated using the Illumina HiSeq platform and Illumina TruSeq Stranded Total RNA-Seq protocol, producing 2×100bp paired-end reads. The RNA-Seq FASTQ files from the EAGLE study we used for this analysis are also available in dbGaP under the accession number phs002346.v1.p1^115^. The RNA-seq FASTQ files from the TCGA-LUAD study were downloaded from the GDC Data Portal^116^. The FASTQ files were aligned to human reference genome GRCh38/hg38 using STAR^117^, annotated using the GENCODE v35, and processed by HTSeq^118^ for gene quantification. The resulting expression data were corrected for batch effects using the R package ComBat-seq^119^ and normalized to Counts Per Million (CPM), followed by log2 transformation using the R package edgeR^120^. Only ‘expressed’ genes defined by CPM >0.1 in at least 10% QC-passed samples were included in the final quantification data.

We performed multivariate analyses to identify genes whose expression was associated with TTFC after adjusting for tumor purity and copy number status. Regression analyses were performed separately within the LAS and HAS subtypes. Genes with an association P<0.05 were selected for pathway analyses using Ingenuity Pathway Analysis (IPA). In addition, we performed differentially expressed gene (DEG) analyzes between LAS and HAS tumors. The batch effect-corrected read counts were used as input for SARTools (v.1.8.0)^121^, and edgeR (v3.36)^120^ was used to identify the DEGs. The top 500 significant DEGs within each comparison were selected for pathway analyses by IPA. Significant pathways were selected based on |Z-score|>1 and FDR<0.05.

### Enrichment and association validation using TCGA and PCAWG data

For TCGA PanCancer data, we downloaded the genomic alterations, gene expression, and clinical information from the cBioPortal for Cancer Genomics (https://www.cbioportal.org/datasets; study: TCGA PanCancer Atlas). For PCAWG PanCancer data, the genomic and clinical information was collected from the ICGC Data Portal (https://dcc.icgc.org/pcawg). The mutational signature decomposition data for both TCGA and PCAWG studies were extracted from a previous publication^4^ based on the SigProfilerExtractor algorithm. A minimum of 50 mutations assigned to SBS2+SBS13 was required for the identification of HAS tumors. Tumor purity was derived from a consensus measurement of purity estimates (CPE) from a previous study^122^. Logistic regression was used for genomic features enrichment analyses between LAS and HAS tumors, adjusting for age, sex, TMB, and tumor purity. The Mann-Whitney (Wilcoxon rank-sum) test was used to compare the age of diagnosis between LAS and HAS with at least 5 patients for each group. In addition, DNA methylation and tobacco smoking data from TCGA LUAD were downloaded from the Genomic Data Commons (GDC) data portal (https://portal.gdc.cancer.gov/).

### Statistical analyses

All statistical analyses were performed using the R software v4.1.2 (https://www.r-project.org/). In general, the Mann-Whitney (Wilcoxon rank sum) test was used for two-group comparisons and the two-sided Fisher exact test was applied for the enrichment analysis of two categorical variables. P[<0.05 was considered statistically significant. If multiple testing was required, we applied the FDR correction based on the Benjamini–Hochberg method. For the survival analyses, a proportional-hazards model was used to investigate the associations between APOBEC subtypes and overall survival, adjusting for age, sex, stage, and histology.

## DATA AVAILABILITY

WGS raw data (CRAM files) and methylation raw data (intensity idat files) used for this work have been deposited in the dbGaP under accession number phs002992.v1.p1. The RNA-Seq raw data (FASTQ files) from EAGLE subjects can be accessed through dbGaP with the accession number phs002346.v1.p1. The access information for the public multi-omics datasets can be found in **Supplementary Table 1**.

## CODE AVAILABILITY

The major bioinformatics pipelines including genome-wide somatic callings based on WGS data can be found at https://github.com/xtmgah/Sherlock-Lung. Battenberg SCNA calling algorithm can be found at https://github.com/Wedge-lab/battenberg. Dirichlet process-based methods for the subclonal reconstruction of tumors can be found at https://github.com/Wedge-lab/dpclust. Mutational signature analysis SigProfilerExtractor can be found at https://github.com/AlexandrovLab/SigProfilerExtractor. The code for P-MACD can be found at https://github.com/NIEHS/P-MACD. The algorithm for dentification of clustered mutations and events is available at https://github.com/AlexandrovLab/SigProfilerClusters.

## ACKNOWLEDGEMENTS

This work has been supported by the Intramural Research Program of the Division of Cancer Epidemiology and Genetics (project ZIACP101231 to M.T.L.), National Cancer Institute, and by the Intramural Research Program of the National Institute of Environmental Health Sciences (project Z1AES103266 to D.A.G), National Institutes of Health (NIH). Cancer research in the Harris lab is supported by NCI P01-CA234228, NCI P50-CA247749, and a CPRIT Established Investigator Recruitment Award. RSH is an Investigator of the Howard Hughes Medical Institute and the Ewing Halsell President’s Council Distinguished Chair at University of Texas Health San Antonio. Research performed for this publication at the Alexandrov Lab was supported by a Packard Fellowship for Science and Engineering as well as by grants from the US National Institutes of Health, including: NIEHS R01ES030993-01A1, NIEHS R01ES032547, and NCI R01CA269919.

We thank Cameron Durfee for helpful comments. This work utilized the computational resources of the NIH HPC Biowulf cluster (https://hpc.nih.gov).

## ETHICS DECLARATIONS

LBA is a compensated consultant and has equity interest in io9, LLC. His spouse is an employee of Biotheranostics, Inc. LBA is also an inventor of a US Patent 10,776,718 for source identification by non-negative matrix factorization. ENB and LBA declare U.S. provisional patent applications with serial numbers 63/289,601 and 63/269,033. LBA also declares U.S. provisional patent applications with serial numbers: 63/366,392; 63/367,846; and 63/412,835. All other authors declare no competing interests.

---

**Extended Data Fig. 1: Characterization of APOBEC mutagenesis based on P-MACD approach between LAS and HAS tumors. a,** Comparison of APOBEC mutational signatures detection between the NMF-based SigProfileExtractor approach and the mutational pattern-based P-MACD approach. APOBEC mutation detection is almost identical between the two approaches in 135 tumors, while P-MACD identifies APOBEC mutations below the detection limit of SigProfileExtractor in 158 tumors. **b,c,** Comparisons of minimal estimated APOBEC mutation load (**b**) and percentage of hypermutable strand-coordinated ssDNA (scssDNA) per genome due to A3A mutagenesis (**c**) between LAS and HAS tumors.

**Extended Data Fig. 2: a-d, Enrichment of *TP53* and *KRAS* mutations between LAS and HAS tumors in all samples (a,b) and LUAD samples only (c-d).** P-values of the Fisher’s exact test are shown on the top of each barplot.

**Extended Data Fig. 3: a, Logistic regression analysis between tumor subtypes and nonsynonymous mutation status of driver genes in TCGA LUAD dataset, adjusting for the following covariates: age, sex, histology, TMB, and tumor purity.** The significance levels P<0.05 (red) and FDR<0.05 (green) are indicated by the dashed lines. Bar plots show *TP53* (**b**) and *KRAS* (**c**) mutation enrichment between LAS and HAS tumors.

**Extended Data Fig. 4: Correlation between minimal estimated APOBEC T<u>C</u>A mutational load from P-MACD and gene expression of *APOBEC3A* and *APOBEC3B*, stratified by LAS and HAS tumors.** Pearson correlation coefficients and P-values are labeled above each plot and in red ink if P<0.05.

**Extended Data Fig. 5: Validation of gene expression correlation between *UNG* and *APOBEC3A* and *APOBEC3B* in all TCGA cancer types. a,** Volcano plot shows the correlations between *UNG* and *APOBEC3A* (blue) and between *UNG* and *APOBEC3B* (yellow). The suggested significance level FDR=0.05 is shown as a dashed red line. **b**, *UNG* gene expression level across all TCGA cancer types sorted by the median *UNG* expression. Cancer type abbreviations from the TCGA study can be found here: https://gdc.cancer.gov/resources-tcga-users/tcga-code-tables/tcga-study-abbreviations.

**Extended Data Fig. 6: Multivariate regression analysis between five smoking variables and the number of mutations assigned to smoking-associated signatures including SBS4 (a), DBS2 (b), and ID3 (c) in EAGLE samples (n=198).** Linear regression analyses performed between LAS and HAS tumors. P-values and regression coefficients with 95% CIs from forest plots are shown for each category of smoking variables. Significant associations are in red ink. Trend test P-values (P_trend_) from associations of TTFC with SBS4, DBS2, and ID3 are included below the forest plots. All association analyses are adjusted for the following covariates: age, sex, histology, and tumor purity.

**Extended Data Fig. 7: Multivariate regression analysis between the smoking variable TTFC and gene mutations in EAGLE samples (n=198). a,** Volcano plot shows the association between each TTFC category and mutation status of commonly mutated genes (Frequency >20%). We performed logistic regression analyses between LAS and HAS tumors, adjusting for the following covariates: age, sex, histology, TMB, and tumor purity. The size of each point on the volcano plot indicates the overall gene mutation frequency. The red and green dashed line indicates the association significance threshold P=0.05, and FDR=0.05, respectively. **b**, Example of an association between each TTFC category and *ZFHX4* mutation frequency stratified by LAS and HAS subtypes. Trend test P-values (P_trend_) are labeled above each subplot.

**Extended Data Fig. 8: Linear regression analysis between five smoking variables and DNA methylation levels of selected CpG probes in normal EAGLE samples (n=122). a,b,** Volcano plots of the associations between methylation levels of each CpG probe and smoking variables independently (**a**) or with CIGT_TIME_LAST_QUIT [Number of years since the subject quitted smoking cigarettes (0 means current smokers)] and CIGT_TOT_DURATION (Total period (in years) during which the subject smoked cigarettes regularly. (**b**) Association FDR values (adjusted using the Benjamini-Hochberg method) are shown on the y-axis of each volcano plot. The orange dashed line indicates the associations with FDR<0.1. The CpG probes associated with tobacco smoking are derived from a study^65^ comparing methylation levels between smokers and never smokers in normal lung tissue. The size and color of each point represent the FDR and association direction, respectively. All association analyses are adjusted for the following covariates: age and sex.

**Supplementary Fig. 1: Summary of multi-omics data and clinical information in this study.**

**Supplementary Fig. 2: Landscape of mutational processes for DBS (a) and ID (b)**. The landscape of mutational signature plots include a bar plot presenting the total number of mutations assigned to each signature, the proportion plot of signatures assigned to each sample, and cosine similarity between the original mutation profile and signature decomposition.

**Supplementary Fig. 3: a-d, Comparison of genomic alterations (a), tobacco smoking exposure (b), clinical information (c), and germline APOBEC3B deletion (d) between LAS and HAS tumors.** The P-values derived from the Wilcoxon sum rank test or Chi-squared test are shown above the plots.

**Supplementary Fig. 4: Genomic characterizations of LAS and HAS tumors.** The subplots from top to bottom: distribution of genomic alteration numbers, most frequently mutated or potential driver genes, oncogenic fusions, significant focal SCNA, and different genomic features. The numbers on the right of each subplot show the overall frequency or median values.

**Supplementary Fig. 5: a,b, Logistic regression analysis between tumor subtypes and driver mutation status of driver genes, adjusting for the following covariates: age, sex, histology, TMB, and tumor purity.** The significance thresholds P<0.05 (red) and FDR<0.05 (green) are indicated by the dashed lines.

**Supplementary Fig. 6: Differentially expressed *UNG* between LAS and HAS tumors based on all available tumors (a) or only in tumors with copy-neutral status for the *UNG* genomic location (b).** Significant P-values from the linear regression are labeled below each boxplot. The linear regression model was adjusted for the following covariates: tumor purity and copy number status in (**a**) and tumor purity only in (**b**).

**ASupplementary Fig. 7: Gene expression correlation between *UNG* and *APOBEC3A* or *APOBEC3B* in normal samples from this study (a), tumor samples from TCGA LUAD (b), and normal samples from TCGA LUAD (c).** Significant P-values and Pearson correlation coefficients are shown on the top of each scatter plot.

**ASupplementary Fig. 8: Volcano plots of the expression correlations between genes in the base excision repair pathway and *APOBEC3A* (a) or *APOBEC3B* (b) in tumors, and between genes in the base excision repair pathway and *APOBEC3A* (c) or *APOBEC3B* (d) in normal samples.** The red dashed line indicates the significance threshold FDR=0.05.

**Supplementary Fig. 9: APOBEC mutation clonality. a,** Percentage of APOBEC subclonal mutations in each tumor from this study. **b**, Fold changes between relative proportions of subclonal and clonal mutations attributed to individual mutational signatures. Box plots demarcate the first and third quartiles of the distribution, with the median shown in the center and whiskers covering data within 1.5× the IQR from the box. **c**, Percentage of APOBEC subclonal mutations in all cancer types with clonality data from the PCAWG study. For each cancer type, the x-axis shows the number of samples with APOBEC mutations over the total number of samples with clonality data.

**Supplementary Fig. 10: Pairwise Pearson correlation between the number of mutations assigned to two observed mutational signatures or overall TMB.** The color and size of the point indicate the correlation coefficients. *P<0.05, [[P<0.01 and [[[P<0.001.

**Supplementary Fig. 11: Distributions of the values of each smoking variable in the EAGLE samples (n=198).**

**Supplementary Fig. 12: Forest plot for the associations between TMB and smoking variables, stratified between LAS and HAS tumors in LUAD samples only.** Forest plot for the associations between TMB and smoking variables, stratified between LAS and HAS tumors. P-values and regression coefficients with 95% confidence intervals (CIs) are shown for each category of smoking variables. Significant associations are in red ink. Trend test P values (P_trend_) from associations between TTFC and TMB are included below the forest plots.

**Supplementary Fig. 13: Linear regression between the smoking variable TTFC and the number of mutations among SBS, DBS, and ID in LAS and HAS tumors (n=198).** TTFC >60 mins is used as the reference for the associations, adjusting for the following covariates: age, sex, histology, and tumor purity. The associations are performed by separating the LAS (circles) and HAS (triangles) tumors. The orange dashed line indicates the significance threshold of P=0.05.

**Supplementary Fig. 14: Comparison of germline variants from a GWAS of nicotine addiction between LAS and HAS tumors.** The P-values derived from the Chi-squared test are shown above the plots.

**Supplementary Fig. 15: Multivariate** regression analysis between the DNA methylation level of CpG probe cg05575921 within *AHRR* and the smoking variable CIGT_TIME_LAST_QUIT [Number of years since the subject quitted smoking cigarettes (0 means current smokers)] (a) and smoking status (b) in tumor and normal EAGLE tissue samples (n=122). The association analyses are performed on all tumors and separately between LAS and HAS tumor subtypes. Trend test P-values (P_trend_) are labeled above each subplot. The linear regression model was adjusted for the following covariates in tumor: age, sex, histology, and tumor purity; and in normal tissue: age and sex.

**Supplementary Fig. 16: Multivariate** regression analysis between the smoking variable CIGT_TIME_LAST_QUIT [Number of years since the subject quitted smoking cigarettes (0 means current smokers)] and DNA methylation level of CpG probe cg14120703 within *NOTCH1* in tumor or normal EAGLE tissue samples (n=122). The association analyses are performed on all subjects and separately between LAS and HAS tumor. Trend test P-values (P_trend_) are labeled above each subplot. The linear regression model was adjusted for the following covariates in tumor: age, sex, histology, and tumor purity; and in normal tissue: age and sex.

**ASupplementary Fig. 17: Association between smoking status and DNA methylation levels of the *AHRR* CpG probe cg05575921 and cg14120703 from *NOTCH1* in the TCGA LUAD dataset.** The linear regression analyses are performed separately between LAS and HAS tumor. Trend test P-values (P_trend_) are labeled above each subplot.

**Supplementary Fig. 18: Clustering of pathways significantly associated with short TTFC in LUAD tumors from EAGLE (n=105).** Pathway analyses using RNA-Seq data from lung tumor tissue separately between LAS and HAS tumors. Only pathways with absolute Z-score>1 and FDR<0.05 are included.

**Supplementary Fig. 19: Inferring patient-specific LUAD cell-of-origin from somatic mutational patterns. a,** Hierarchically clustered heatmap showing correlations between scRNA-seq expression signature and somatic SNV densities across individual WGS samples. Estimated relative risk from the maximum likelihood regression fit is plotted for each tumor sample and lung cell type combination. **b,** Pie charts show the percentage of each inferred cell-of-origin in LAS and HAS tumors.

**Supplementary Fig. 20: Comparison of the estimated age at the occurrence of the most recent common ancestor (MRCA) in all tumors (a) and in tumors with TTFC<=5 mins (b), and latency in all tumors (c) and in tumors with TTFC <=5 mins (d) between LAS and HAS tumors.** Tumor cell latency is calculated as the difference between age at diagnosis and the estimated age at the occurrence of MRCA based on 5x acceleration rate. The P-value from the Wilcoxon sum rank test is shown above the boxplot.

**Supplementary Fig. 21: Age at diagnosis differences in patients based on *TP53* mutation (a) status and *RAS* mutation status (b).**

**Supplementary Fig. 22: Neo**antigen burden between LAS and HAS from the TCGA LUAD dataset. P-values from Wilcoxon rank-sum tests are labeled on the top of the boxplot.

**Supplementary Fig. 23: *PD-L1* expression difference between LAS and HAS tumors from this study (a) and the TCGA LUAD study (b).**

**Supplementary Fig. 24: Kaplan–Meier survival curves for overall survival stratified by tumor subtype (LAS vs. HAS).**

**Supplementary Fig. 25: Comparison of age at diagnosis between LAS and HAS tumors from the PCAWG study across multiple cancer types using SBS2 and SBS13 to define HAS and LAS subtypes (a) and skin-melanoma only using SBS13 to define LAS and HAS subtypes (b).** Only cancer types with sufficient tumors in both APOBEC LAS and HAS subtypes for comparison are included. P-values from the Wilcoxon sum rank test are shown above the boxplot.

**Supplementary Fig. 26: Comparisons of tumors stratified based on the detection of SBS40 signatures with regard to *TP53* mutation frequency (a), *KRAS* mutation frequency (b) and age at diagnosis (c).** P-values of the Fisher’s exact test or Wilcoxon sum rank test are shown on the top of each bar plot.

## TABLES

**Supplementary Table 1:** Overall data summary and genomic features included in this study.

**Supplementary Table 2:** Mutational signature deconvolution for SBS, DBS, and ID mutation types using the COSMIC mutational signature catalog as reference.

**Supplementary Table 3:** Smoking history information for the EAGLE study subjects.

**Supplementary Table 4:** Distinct patterns of mutagenesis by APOBEC cytidine deaminases using the P-MACD algorithm.

**Supplementary Table 5:** RNA-Seq expression data quantified as log2CPM for genes involved in the major analyses.

**Supplementary Table 6:** Methylation beta value for 952 CpG probes that were found to have genome-wide significant associations with smoking variables in a previous study^65^.

**Supplementary Table 7:** Associations between smoking variables and methylation levels of known smoking-related CpG probes in both normal and tumor tissue samples.

**Supplementary Table 8:** Differentially expressed gene markers of lung-specific cell types between LAS and HAS LUAD tumors (n=155) adjusting for copy number alterations and tumor purity.

## Notes

### Summary of Updates

Correcting some encoding errors detected in the submission system.

## REFERENCE

1. Mertz, T. M., Collins, C. D., Dennis, M., Coxon, M. & Roberts, S. A. APOBEC-Induced Mutagenesis in Cancer. Annu. Rev. Genet. (2022) doi:10.1146/annurev-genet-072920-035840.

2. Pecori, R., Di Giorgio, S., Paulo Lorenzo, J. & Nina Papavasiliou, F. Functions and consequences of AID/APOBEC-mediated DNA and RNA deamination. Nat. Rev. Genet. (2022) doi:10.1038/s41576-022-00459-8.

3. Roberts, S. A. et al. An APOBEC cytidine deaminase mutagenesis pattern is widespread in human cancers. Nat. Genet. 45, 970–976 (2013).

4. Alexandrov, L. B. et al. The repertoire of mutational signatures in human cancer. Nature 578, 94–101 (2020).

5. Degasperi, A. et al. Substitution mutational signatures in whole-genome-sequenced cancers in the UK population. Science 376, (2022).

6. Petljak, M. et al. Mechanisms of APOBEC3 mutagenesis in human cancer cells. Nature 1–9 (2022).

7. Da Silva-Álvarez, S. et al. Cell senescence contributes to tissue regeneration in zebrafish. Aging Cell 19, e13052 (2020).

8. Bergmann, A. & Steller, H. Apoptosis, stem cells, and tissue regeneration. Sci. Signal. 3, re8 (2010).

9. Eming, S. A., Wynn, T. A. & Martin, P. Inflammation and metabolism in tissue repair and regeneration. Science 356, 1026–1030 (2017).

10. Stratton, M. R., Campbell, P. J. & Futreal, P. A. The cancer genome. Nature 458, 719–724 (2009).

11. Alexandrov, L. B. & Stratton, M. R. Mutational signatures: the patterns of somatic mutations hidden in cancer genomes. Curr. Opin. Genet. Dev. 24, 52–60 (2014).

12. Koh, G., Degasperi, A., Zou, X., Momen, S. & Nik-Zainal, S. Mutational signatures: emerging concepts, caveats and clinical applications. Nat. Rev. Cancer 21, 619–637 (2021).

13. Wang, S., Jia, M., He, Z. & Liu, X.-S. APOBEC3B and APOBEC mutational signature as potential predictive markers for immunotherapy response in non-small cell lung cancer. Oncogene 37, 3924–3936 (2018).

14. Langenbucher, A. et al. An extended APOBEC3A mutation signature in cancer. Nat. Commun. 12, 1602 (2021).

15. Granadillo Rodríguez, M., Flath, B. & Chelico, L. The interesting relationship between APOBEC3 deoxycytidine deaminases and cancer: a long road ahead. Open Biol. 10, 200188 (2020).

16. Harris, R. S. & Dudley, J. P. APOBECs and virus restriction. Virology 479-480, 131–145 (2015).

17. Cheng, A. Z. et al. APOBECs and Herpesviruses. Viruses 13, (2021).

18. Burns, M. B., Temiz, N. A. & Harris, R. S. Evidence for APOBEC3B mutagenesis in multiple human cancers. Nat. Genet. 45, 977–983 (2013).

19. Chan, K. et al. An APOBEC3A hypermutation signature is distinguishable from the signature of background mutagenesis by APOBEC3B in human cancers. Nat. Genet. 47, 1067–1072 (2015).

20. Petljak, M. et al. Characterizing Mutational Signatures in Human Cancer Cell Lines Reveals Episodic APOBEC Mutagenesis. Cell 176, 1282–1294.e20 (2019).

21. Petljak, M. & Maciejowski, J. Molecular origins of APOBEC-associated mutations in cancer. DNA Repair 94, 102905 (2020).

22. Burgess, D. J. Switching APOBEC mutation signatures. Nature reviews. Genetics vol. 20 253 (2019).

23. Kazanov, M. D. et al. APOBEC-Induced Cancer Mutations Are Uniquely Enriched in Early-Replicating, Gene-Dense, and Active Chromatin Regions. Cell Rep. 13, 1103–1109 (2015).

24. Haradhvala, N. J. et al. Mutational Strand Asymmetries in Cancer Genomes Reveal Mechanisms of DNA Damage and Repair. Cell 164, 538–549 (2016).

25. Hoopes, J. I. et al. APOBEC3A and APOBEC3B Preferentially Deaminate the Lagging Strand Template during DNA Replication. Cell Rep. 14, 1273–1282 (2016).

26. Venkatesan, S. et al. Induction of APOBEC3 Exacerbates DNA Replication Stress and Chromosomal Instability in Early Breast and Lung Cancer Evolution. Cancer Discov. 11, 2456–2473 (2021).

27. Jakobsdottir, G. M., Brewer, D. S., Cooper, C., Green, C. & Wedge, D. C. APOBEC3 mutational signatures are associated with extensive and diverse genomic instability across multiple tumour types. BMC Biol. 20, 117 (2022).

28. Siegel, R. L., Miller, K. D., Fuchs, H. E. & Jemal, A. Cancer statistics, 2022. CA Cancer J. Clin. 72, 7–33 (2022).

29. Alexandrov, L. B. et al. Mutational signatures associated with tobacco smoking in human cancer. Science 354, 618–622 (2016).

30. Islam, S. M. A. et al. Uncovering novel mutational signatures by de novo extraction with SigProfilerExtractor. Cell Genom 2, None (2022).

31. Nagahashi, M. et al. Common driver mutations and smoking history affect tumor mutation burden in lung adenocarcinoma. J. Surg. Res. 230, 181–185 (2018).

32. Wang, X. et al. Association between Smoking History and Tumor Mutation Burden in Advanced Non-Small Cell Lung Cancer. Cancer Res. 81, 2566–2573 (2021).

33. Wang, R., Li, S., Wen, W. & Zhang, J. Multi-Omics Analysis of the Effects of Smoking on Human Tumors. Front Mol Biosci 8, 704910 (2021).

34. Swanton, C., McGranahan, N., Starrett, G. J. & Harris, R. S. APOBEC Enzymes: Mutagenic Fuel for Cancer Evolution and Heterogeneity. Cancer Discov. 5, 704–712 (2015).

35. Landi, M. T. et al. Environment And Genetics in Lung cancer Etiology (EAGLE) study: an integrative population-based case-control study of lung cancer. BMC Public Health 8, 203 (2008).

36. Alexandrov, L. B., Nik-Zainal, S., Wedge, D. C., Campbell, P. J. & Stratton, M. R. Deciphering signatures of mutational processes operative in human cancer. Cell Rep. 3, 246–259 (2013).

37. Rosenthal, R., McGranahan, N., Herrero, J., Taylor, B. S. & Swanton, C. DeconstructSigs: delineating mutational processes in single tumors distinguishes DNA repair deficiencies and patterns of carcinoma evolution. Genome Biol. 17, 31 (2016).

38. Degasperi, A. et al. A practical framework and online tool for mutational signature analyses show inter-tissue variation and driver dependencies. Nat Cancer 1, 249–263 (2020).

39. Symer, D. E. et al. Human l1 retrotransposition is associated with genetic instability in vivo. Cell 110, 327–338 (2002).

40. Petljak, M., Green, A. M., Maciejowski, J. & Weitzman, M. D. Addressing the benefits of inhibiting APOBEC3-dependent mutagenesis in cancer. Nat. Genet. (2022) doi:10.1038/s41588-022-01196-8.

41. Krokan, H. E. et al. Error-free versus mutagenic processing of genomic uracil--relevance to cancer. DNA Repair 19, 38–47 (2014).

42. Lindahl, T. Suppression of spontaneous mutagenesis in human cells by DNA base excision-repair. Mutat. Res. 462, 129–135 (2000).

43. Serebrenik, A. A. et al. The deaminase APOBEC3B triggers the death of cells lacking uracil DNA glycosylase. Proc. Natl. Acad. Sci. U. S. A. 116, 22158–22163 (2019).

44. Roelofs, P. A. et al. Characterization of the mechanism by which the RB/E2F pathway controls expression of the cancer genomic DNA deaminase APOBEC3B. Elife 9, (2020).

45. Gerstung, M. et al. The evolutionary history of 2,658 cancers. Nature 578, 122–128 (2020).

46. Gu, F. et al. Time to smoke first morning cigarette and lung cancer in a case-control study. J. Natl. Cancer Inst. 106, dju118 (2014).

47. Zhang, T. et al. Time to first cigarette and its impact on lung tumorigenesis. bioRxiv 2022.11.07.515434 (2022) doi:10.1101/2022.11.07.515434.

48. Sancar, A. et al. Circadian clock control of the cellular response to DNA damage. FEBS Lett. 584, 2618–2625 (2010).

49. Wang, R., Li, S., Wen, W. & Zhang, J. Multi-Omics Analysis of the Effects of Smoking on Human Tumors. Front Mol Biosci 8, 704910 (2021).

50. Quach, B. C. et al. Expanding the genetic architecture of nicotine dependence and its shared genetics with multiple traits. Nat. Commun. 11, 5562 (2020).

51. Wang, J. Y. DNA damage and apoptosis. Cell Death Differ. 8, 1047–1048 (2001).

52. Benchimol, S. p53-dependent pathways of apoptosis. Cell Death Differ. 8, 1049–1051 (2001).

53. d’Adda di Fagagna, F. Living on a break: cellular senescence as a DNA-damage response. Nat. Rev. Cancer 8, 512–522 (2008).

54. Matt, S. & Hofmann, T. G. The DNA damage-induced cell death response: a roadmap to kill cancer cells. Cell. Mol. Life Sci. 73, 2829–2850 (2016).

55. Serrano, M. Senescence helps regeneration. Developmental cell vol. 31 671–672 (2014).

56. Antelo-Iglesias, L., Picallos-Rabina, P., Estévez-Souto, V., Da Silva-Álvarez, S. & Collado, M. The role of cellular senescence in tissue repair and regeneration. Mech. Ageing Dev. 198, 111528 (2021).

57. Zeilinger, S. et al. Tobacco smoking leads to extensive genome-wide changes in DNA methylation. PLoS One 8, e63812 (2013).

58. Philibert, R. A., Beach, S. R. H. & Brody, G. H. Demethylation of the aryl hydrocarbon receptor repressor as a biomarker for nascent smokers. Epigenetics 7, 1331–1338 (2012).

59. Bojesen, S. E., Timpson, N., Relton, C., Davey Smith, G. & Nordestgaard, B. G. AHRR (cg05575921) hypomethylation marks smoking behaviour, morbidity and mortality. Thorax 72, 646–653 (2017).

60. Fasanelli, F. et al. Hypomethylation of smoking-related genes is associated with future lung cancer in four prospective cohorts. Nat. Commun. 6, 1–9 (2015).

61. Guida, F. et al. Dynamics of smoking-induced genome-wide methylation changes with time since smoking cessation. Hum. Mol. Genet. 24, 2349–2359 (2015).

62. Philibert, R. et al. Reversion of AHRR Demethylation Is a Quantitative Biomarker of Smoking Cessation. Front. Psychiatry 7, 55 (2016).

63. Joehanes, R. et al. Epigenetic Signatures of Cigarette Smoking. Circ. Cardiovasc. Genet. 9, 436–447 (2016).

64. Lee, M. K., Hong, Y., Kim, S.-Y., London, S. J. & Kim, W. J. DNA methylation and smoking in Korean adults: epigenome-wide association study. Clin. Epigenetics 8, 103 (2016).

65. Christiansen, C. et al. Novel DNA methylation signatures of tobacco smoking with trans-ethnic effects. Clin. Epigenetics 13, 36 (2021).

66. Zhang, T. et al. Genomic and evolutionary classification of lung cancer in never smokers. Nat. Genet. 53, 1348–1359 (2021).

67. Wang, Z. et al. Deciphering cell lineage specification of human lung adenocarcinoma with single-cell RNA sequencing. Nat. Commun. 12, 6500 (2021).

68. Shankaran, V. et al. IFNgamma and lymphocytes prevent primary tumour development and shape tumour immunogenicity. Nature 410, 1107–1111 (2001).

69. Isozaki, H. et al. Therapy-induced APOBEC3A drives evolution of persistent cancer cells. Nature 620, 393–401 (2023).

70. Alexandrov, L. B. et al. Clock-like mutational processes in human somatic cells. Nat. Genet. 47, 1402–1407 (2015).

71. Saini, N. & Gordenin, D. A. Hypermutation in single-stranded DNA. DNA Repair 91-92, 102868 (2020).

72. Jarvis, M. et al. Mutational impact of APOBEC3B and APOBEC3A in a human cell line. bioRxiv 2022.04.26.489523 (2022) doi:10.1101/2022.04.26.489523.

73. Landry, S., Narvaiza, I., Linfesty, D. C. & Weitzman, M. D. APOBEC3A can activate the DNA damage response and cause cell-cycle arrest. EMBO Rep. 12, 444–450 (2011).

74. Hayashi, M. T., Cesare, A. J., Rivera, T. & Karlseder, J. Cell death during crisis is mediated by mitotic telomere deprotection. Nature 522, 492–496 (2015).

75. De Cecco, M. et al. L1 drives IFN in senescent cells and promotes age-associated inflammation. Nature 566, 73–78 (2019).

76. Xu, X. et al. Evidence for type II cells as cells of origin of K-Ras–induced distal lung adenocarcinoma. Proceedings of the National Academy of Sciences 109, 4910–4915 (2012).

77. Huang, Z. et al. Single-cell analysis of somatic mutations in human bronchial epithelial cells in relation to aging and smoking. Nat. Genet. (2022) doi:10.1038/s41588-022-01035-w.

78. DiMarco, A. V. et al. APOBEC Mutagenesis Inhibits Breast Cancer Growth through Induction of T cell-Mediated Antitumor Immune Responses. Cancer Immunol Res 10, 70–86 (2022).

79. Robertson, A. G. et al. Comprehensive Molecular Characterization of Muscle-Invasive Bladder Cancer. Cell 171, 540–556.e25 (2017).

80. Imielinski, M. et al. Mapping the hallmarks of lung adenocarcinoma with massively parallel sequencing. Cell 150, 1107–1120 (2012).

81. Campbell, J. D. et al. Distinct patterns of somatic genome alterations in lung adenocarcinomas and squamous cell carcinomas. Nat. Genet. 48, 607–616 (2016).

82. Lee, J.-K. et al. Clonal History and Genetic Predictors of Transformation Into Small-Cell Carcinomas From Lung Adenocarcinomas. J. Clin. Oncol. 35, 3065–3074 (2017).

83. Leong, T. L. et al. Deep multi-region whole-genome sequencing reveals heterogeneity and gene-by-environment interactions in treatment-naive, metastatic lung cancer. Oncogene 38, 1661–1675 (2019).

84. Lee, J. J.-K. et al. Tracing Oncogene Rearrangements in the Mutational History of Lung Adenocarcinoma. Cell 177, 1842–1857.e21 (2019).

85. Carrot-Zhang, J. et al. Whole-genome characterization of lung adenocarcinomas lacking the RTK/RAS/RAF pathway. Cell Rep. 34, 108707 (2021).

86. Cancer Genome Atlas Research Network. Comprehensive molecular profiling of lung adenocarcinoma. Nature 511, 543–550 (2014).

87. Bergmann, E. A., Chen, B.-J., Arora, K., Vacic, V. & Zody, M. C. Conpair: concordance and contamination estimator for matched tumor–normal pairs. Bioinformatics 32, 3196–3198 (2016).

88. Pedersen, B. S. et al. Somalier: rapid relatedness estimation for cancer and germline studies using efficient genome sketches. Genome Med. 12, 62 (2020).

89. Sadedin, S. P. & Oshlack, A. Bazam: a rapid method for read extraction and realignment of high-throughput sequencing data. Genome Biol. 20, 78 (2019).

90. Nik-Zainal, S. et al. The life history of 21 breast cancers. Cell 149, 994–1007 (2012).

91. Mermel, C. H. et al. GISTIC2.0 facilitates sensitive and confident localization of the targets of focal somatic copy-number alteration in human cancers. Genome Biol. 12, R41 (2011).

92. Yang, L. et al. Diverse mechanisms of somatic structural variations in human cancer genomes. Cell 153, 919–929 (2013).

93. Chen, X. et al. Manta: rapid detection of structural variants and indels for germline and cancer sequencing applications. Bioinformatics 32, 1220–1222 (2016).

94. Tubio, J. M. C. et al. Mobile DNA in cancer. Extensive transduction of nonrepetitive DNA mediated by L1 retrotransposition in cancer genomes. Science 345, 1251343 (2014).

95. Dentro, S. C., Wedge, D. C. & Van Loo, P. Principles of Reconstructing the Subclonal Architecture of Cancers. Cold Spring Harb. Perspect. Med. 7, (2017).

96. Yuan, K., Macintyre, G., Liu, W., PCAWG-11 working group & Markowetz, F. Ccube: A fast and robust method for estimating cancer cell fractions. bioRxiv 484402 (2018) doi:10.1101/484402.

97. Bergstrom, E. N. et al. SigProfilerMatrixGenerator: a tool for visualizing and exploring patterns of small mutational events. BMC Genomics 20, 685 (2019).

98. Bergstrom, E. N., Kundu, M., Tbeileh, N. & Alexandrov, L. B. Examining clustered somatic mutations with SigProfilerClusters. bioRxiv 2022.02.11.480117 (2022) doi:10.1101/2022.02.11.480117.

99. Sakofsky, C. J. et al. Repair of multiple simultaneous double-strand breaks causes bursts of genome-wide clustered hypermutation. PLoS Biol. 17, e3000464 (2019).

100. Davis, C. F. et al. The somatic genomic landscape of chromophobe renal cell carcinoma. Cancer Cell 26, 319–330 (2014).

101. Gerhauser, C. et al. Molecular Evolution of Early-Onset Prostate Cancer Identifies Molecular Risk Markers and Clinical Trajectories. Cancer Cell 34, 996–1011.e8 (2018).

102. Lambrechts, D. et al. Phenotype molding of stromal cells in the lung tumor microenvironment. Nature Medicine vol. 24 1277–1289 Preprint at 10.1038/s41591-018-0096-5 (2018).

103. Raredon, M. S. B. et al. Single-cell connectomic analysis of adult mammalian lungs. Sci Adv 5, eaaw3851 (2019).

104. Laughney, A. M. et al. Regenerative lineages and immune-mediated pruning in lung cancer metastasis. Nat. Med. 26, 259–269 (2020).

105. Lukassen, S. et al. SARS-CoV-2 receptor ACE2 and TMPRSS2 are primarily expressed in bronchial transient secretory cells. EMBO J. 39, e105114 (2020).

106. Kawaguchi, S., Higasa, K., Shimizu, M., Yamada, R. & Matsuda, F. HLA-HD: An accurate HLA typing algorithm for next-generation sequencing data. Hum. Mutat. 38, 788–797 (2017).

107. Reynisson, B., Alvarez, B., Paul, S., Peters, B. & Nielsen, M. NetMHCpan-4.1 and NetMHCIIpan-4.0: improved predictions of MHC antigen presentation by concurrent motif deconvolution and integration of MS MHC eluted ligand data. Nucleic Acids Res. 48, W449–W454 (2020).

108. Schenck, R. O., Lakatos, E., Gatenbee, C., Graham, T. A. & Anderson, A. R. A. NeoPredPipe: high-throughput neoantigen prediction and recognition potential pipeline. BMC Bioinformatics 20, 264 (2019).

109. Thorsson, V. et al. The Immune Landscape of Cancer. Immunity 48, 812–830.e14 (2018).

110. Martínez-Jiménez, F. et al. A compendium of mutational cancer driver genes. Nat. Rev. Cancer 20, 555–572 (2020).

111. Muiños, F., Martínez-Jiménez, F., Pich, O., Gonzalez-Perez, A. & Lopez-Bigas, N. In silico saturation mutagenesis of cancer genes. Nature 596, 428–432 (2021).

112. Müller, F. et al. RnBeads 2.0: comprehensive analysis of DNA methylation data. Genome Biol. 20, 55 (2019).

113. Leek, J. T., Johnson, W. E., Parker, H. S., Jaffe, A. E. & Storey, J. D. The sva package for removing batch effects and other unwanted variation in high-throughput experiments. Bioinformatics 28, 882–883 (2012).

114. Teschendorff, A. E. A comparison of epigenetic mitotic-like clocks for cancer risk prediction. Genome Med. 12, 56 (2020).

115. Zhao, W. et al. Clinical Implications of Inter- and Intratumor Heterogeneity of Immune Cell Markers in Lung Cancer. JNCI: Journal of the National Cancer Institute vol. 114 280–289 Preprint at 10.1093/jnci/djab157 (2022).

116. Grossman, R. L. et al. Toward a Shared Vision for Cancer Genomic Data. N. Engl. J. Med. 375, 1109–1112 (2016).

117. Dobin, A. et al. STAR: ultrafast universal RNA-seq aligner. Bioinformatics 29, 15–21 (2013).

118. Putri, G. H., Anders, S., Pyl, P. T., Pimanda, J. E. & Zanini, F. Analysing high-throughput sequencing data in Python with HTSeq 2.0. Bioinformatics vol. 38 2943–2945 Preprint at 10.1093/bioinformatics/btac166 (2022).

119. Zhang, Y., Parmigiani, G. & Johnson, W. E. ComBat-seq: batch effect adjustment for RNA-seq count data. NAR Genom Bioinform 2, lqaa078 (2020).

120. Robinson, M. D., McCarthy, D. J. & Smyth, G. K. edgeR: a Bioconductor package for differential expression analysis of digital gene expression data. Bioinformatics 26, 139–140 (2010).

121. Varet, H., Brillet-Guéguen, L., Coppée, J.-Y. & Dillies, M.-A. SARTools: A DESeq2- and EdgeR-Based R Pipeline for Comprehensive Differential Analysis of RNA-Seq Data. PLoS One 11, e0157022 (2016).

122. Aran, D., Sirota, M. & Butte, A. J. Systematic pan-cancer analysis of tumour purity. Nat. Commun. 6, 8971 (2015).

