## Extended_Figures for "APOBEC shapes tumor evolution and age at onset of lung cancer in smokers"

#### Extended Data Fig. 1

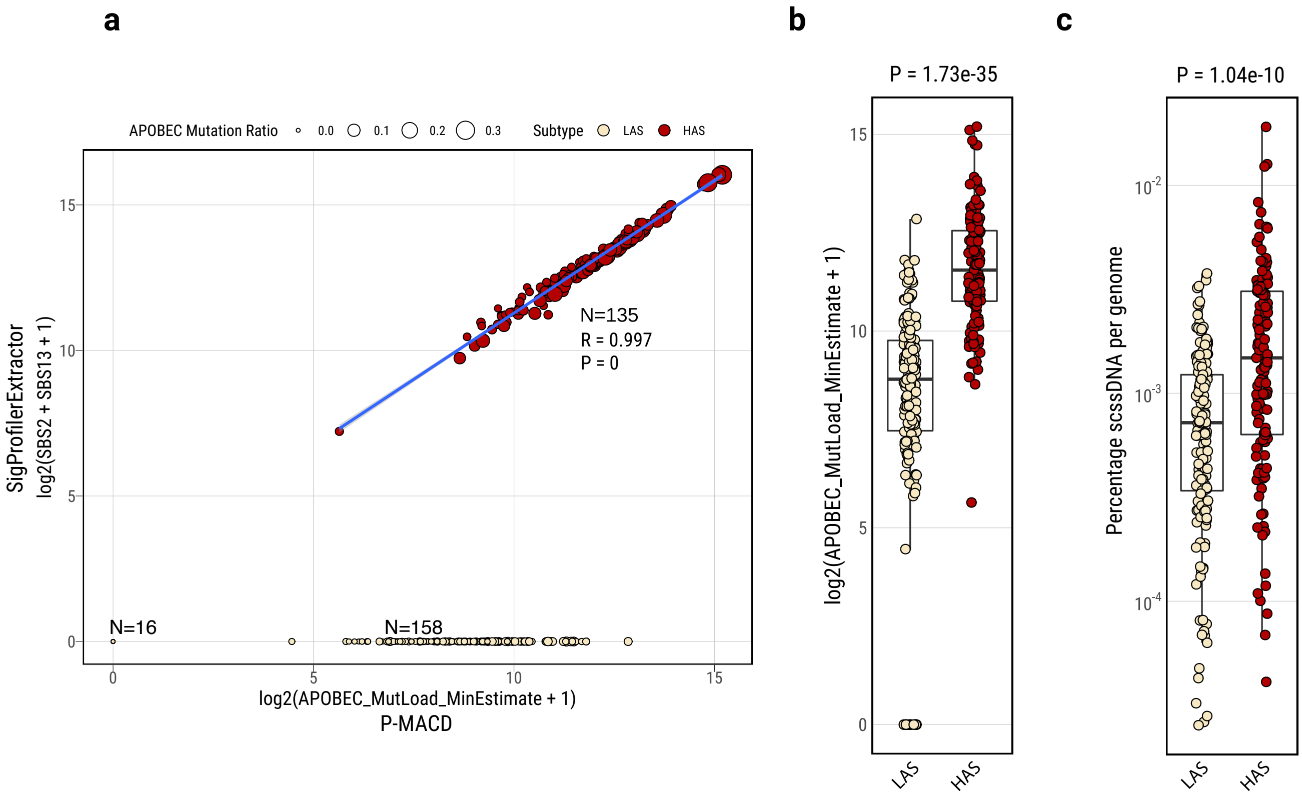

**Extended Data Fig. 2**

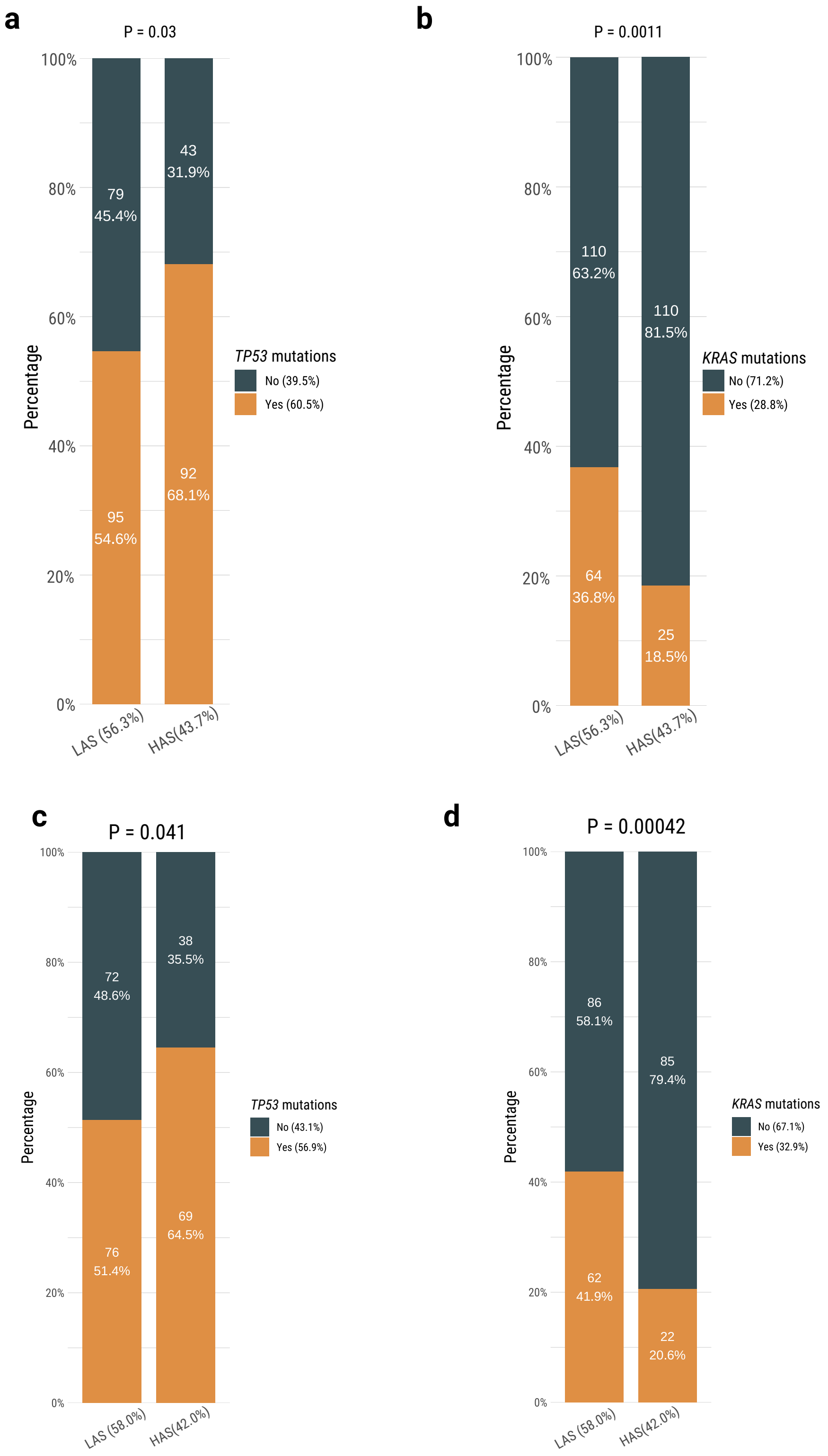

**Extended Data Fig. 3**

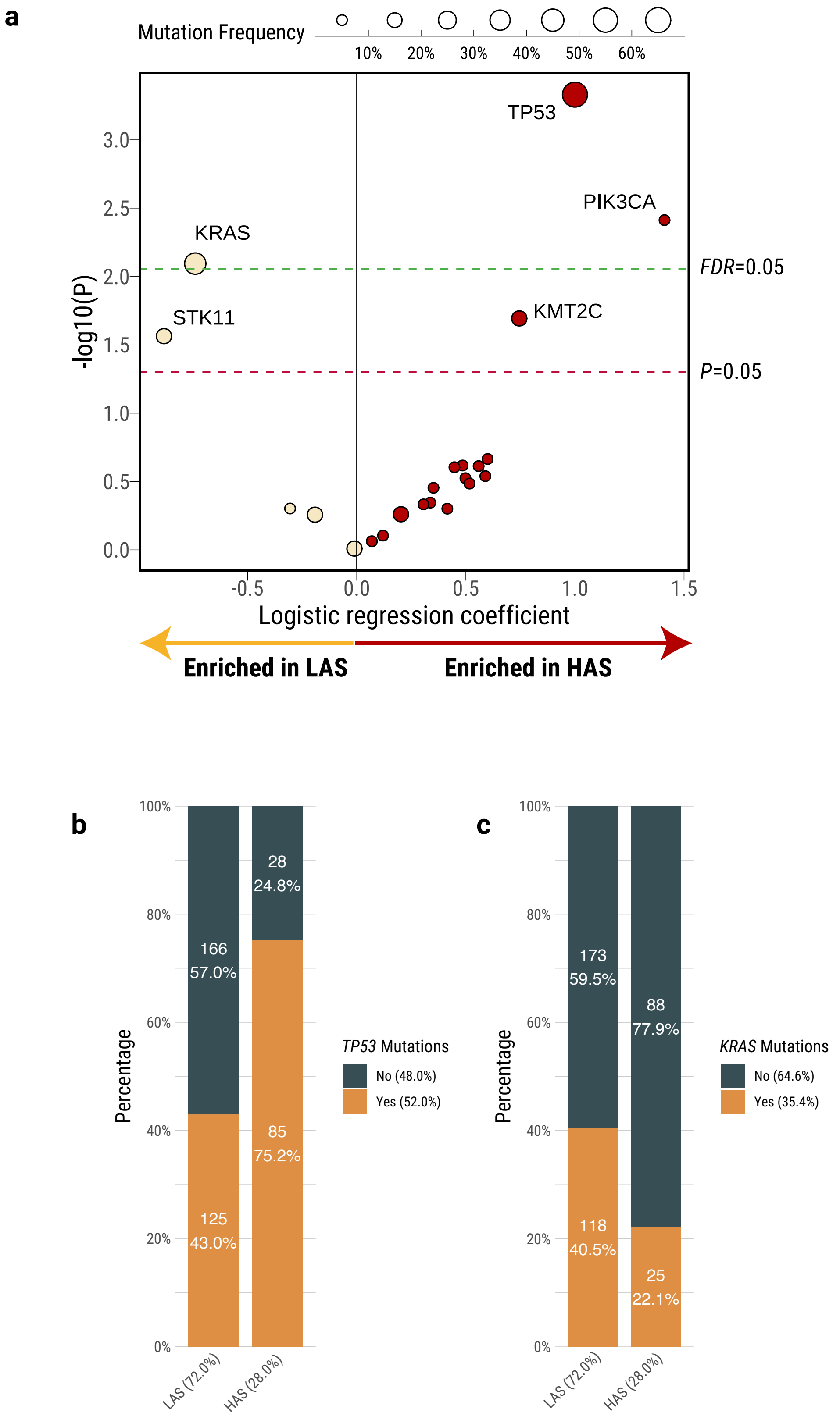

**Extended Data Fig. 4**

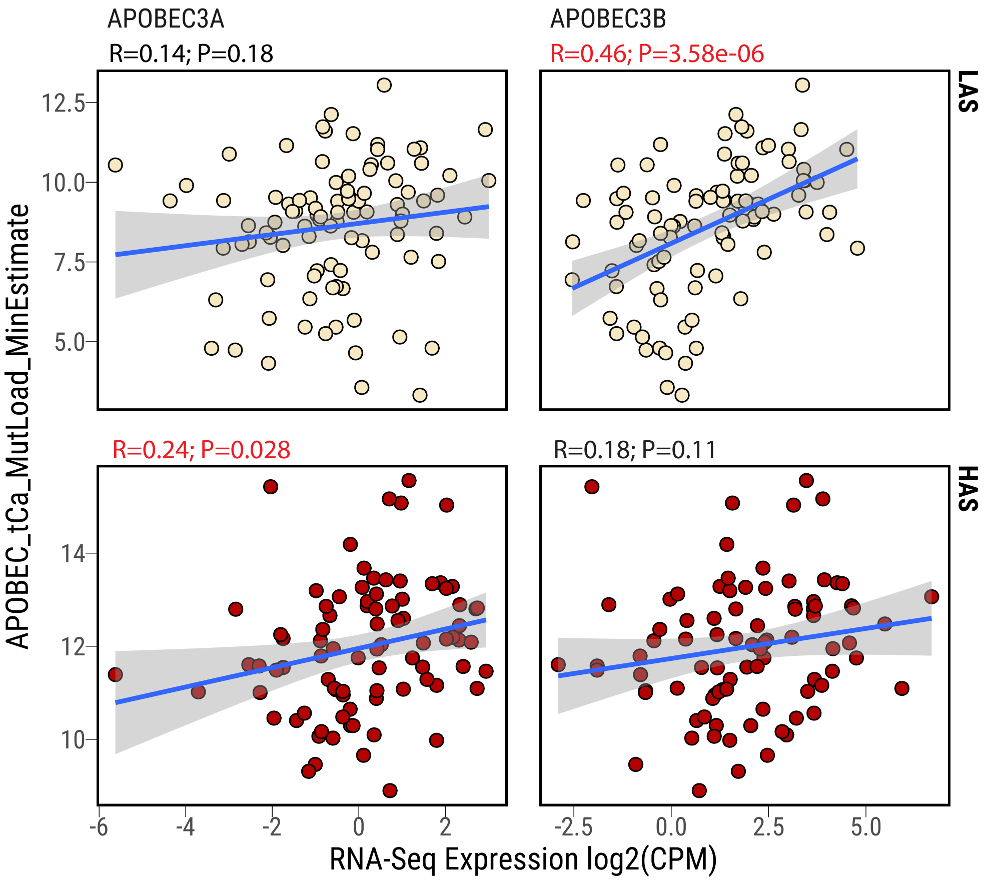

#### Extended Data Fig. 5

**a**

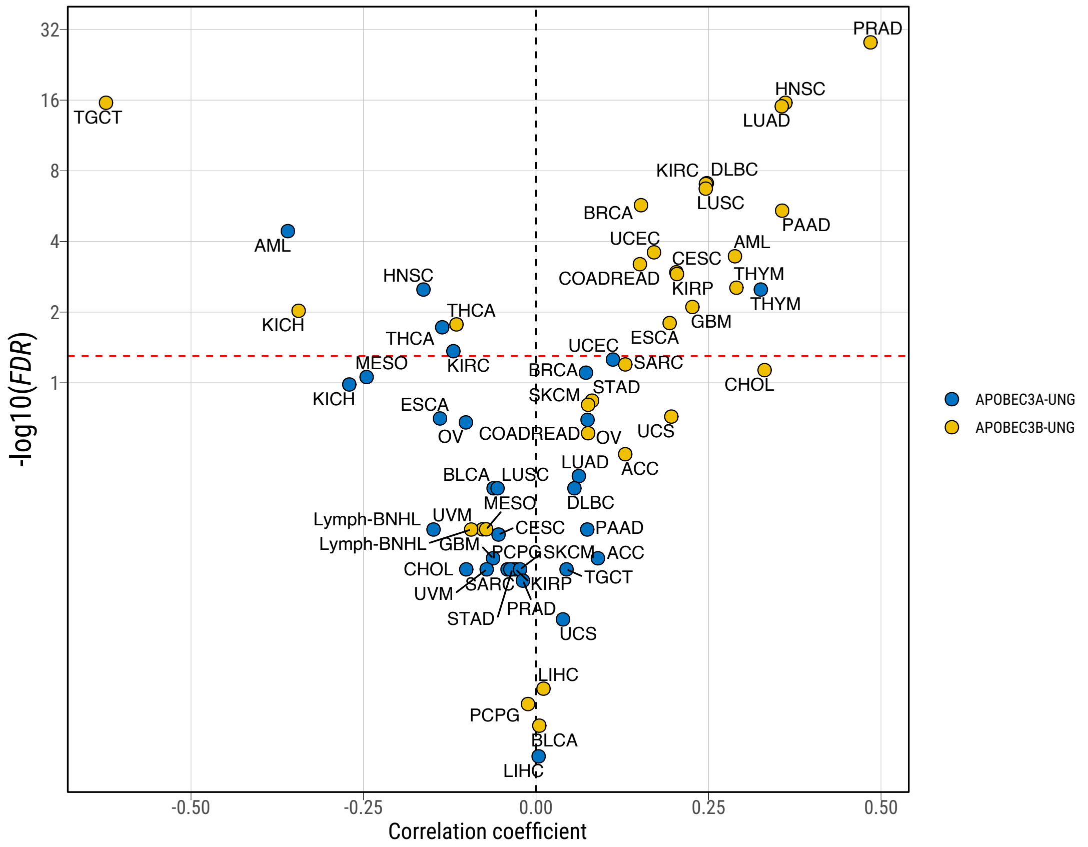


**b**

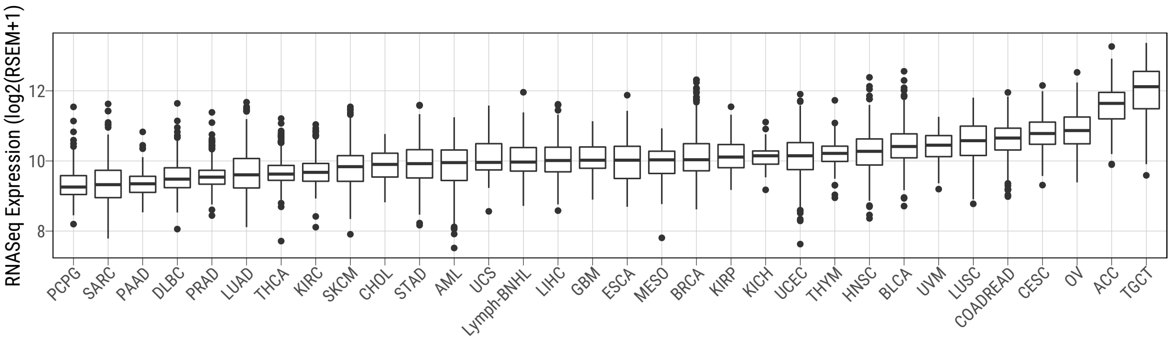

### Extended Data Fig. 6

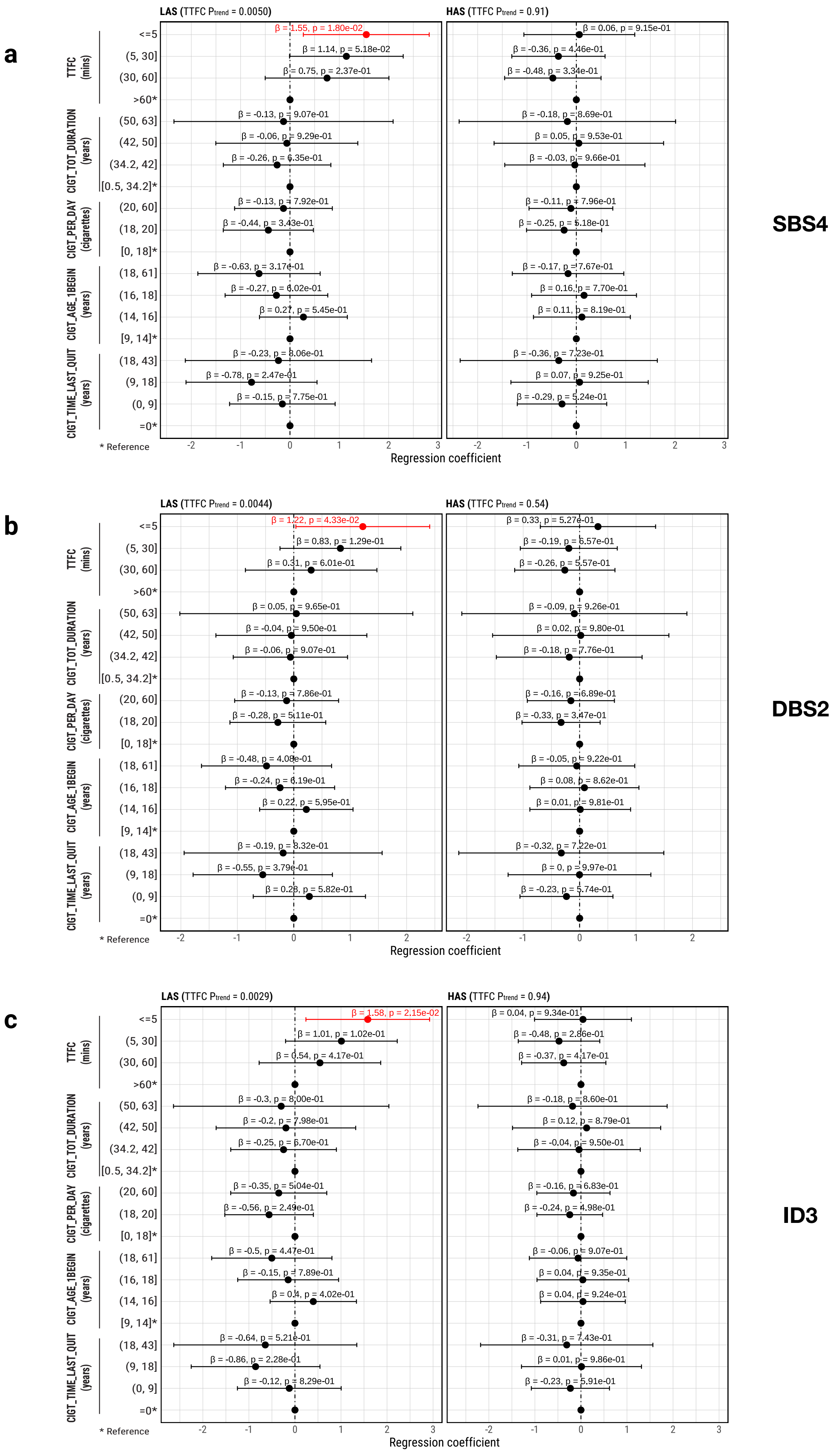

#### Extended Data Fig. 7

**a**

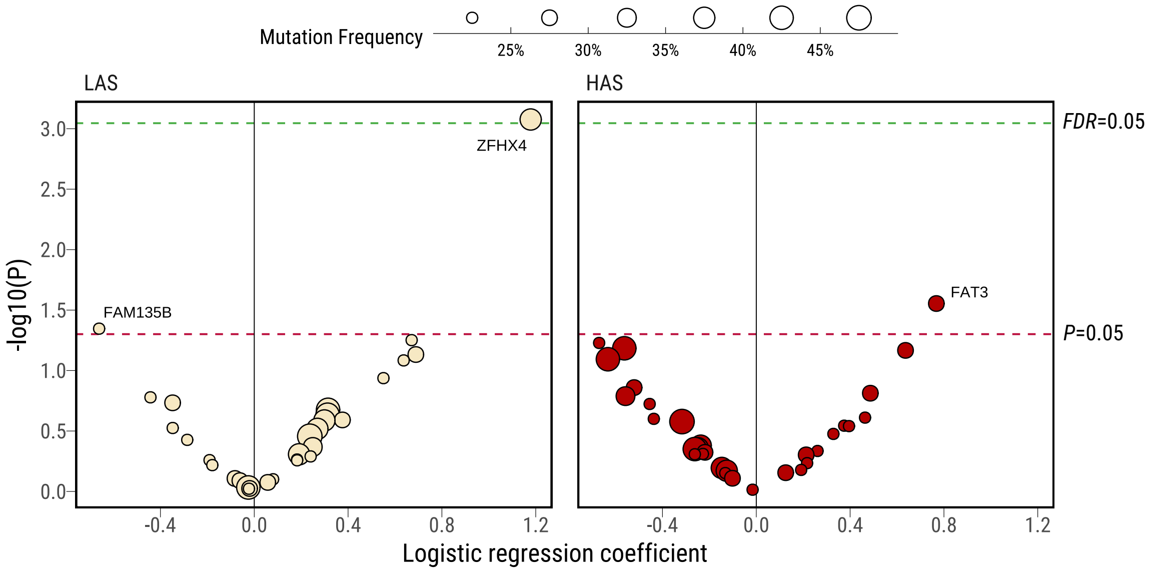


**b**

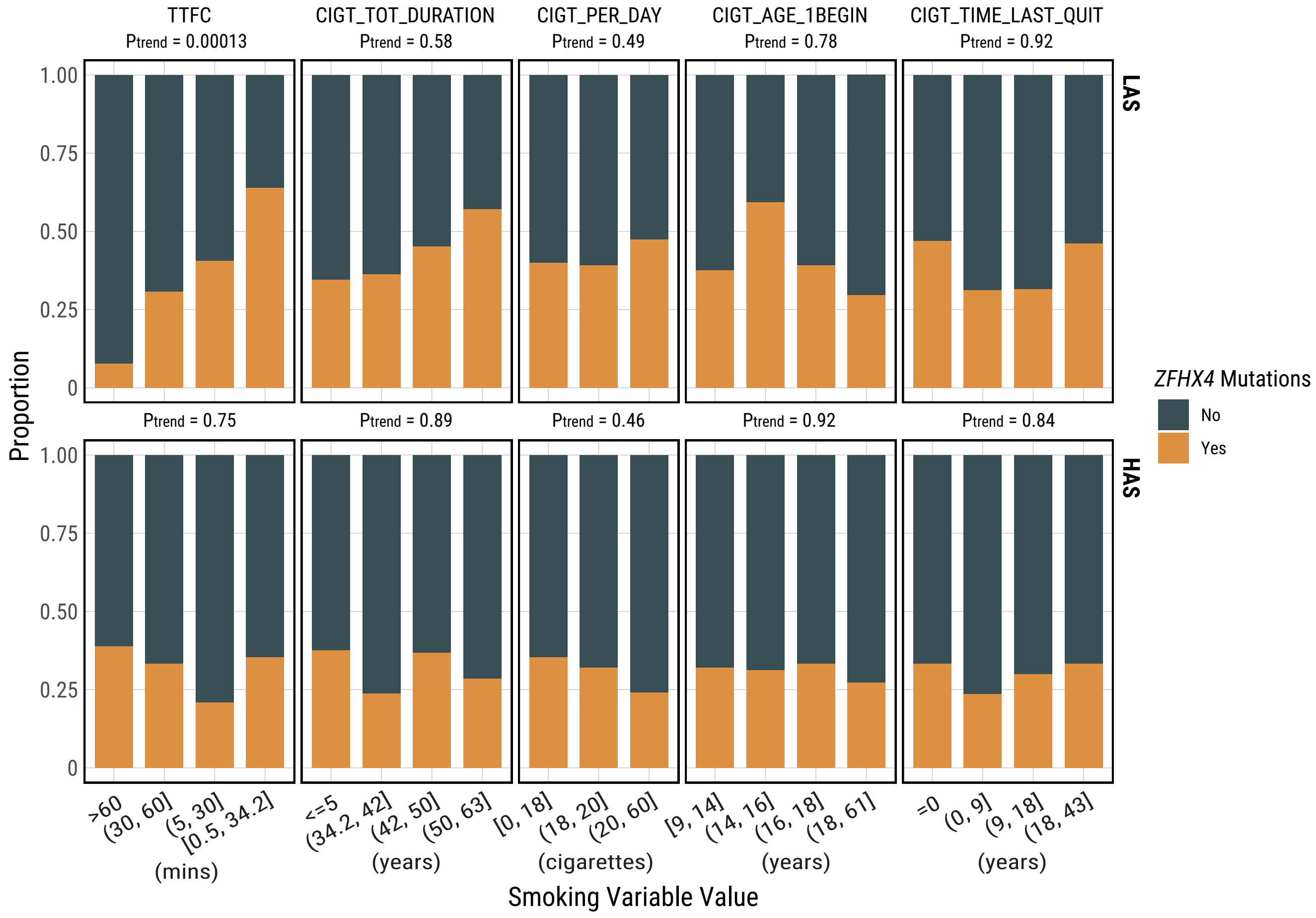

#### Extended Data Fig. 8

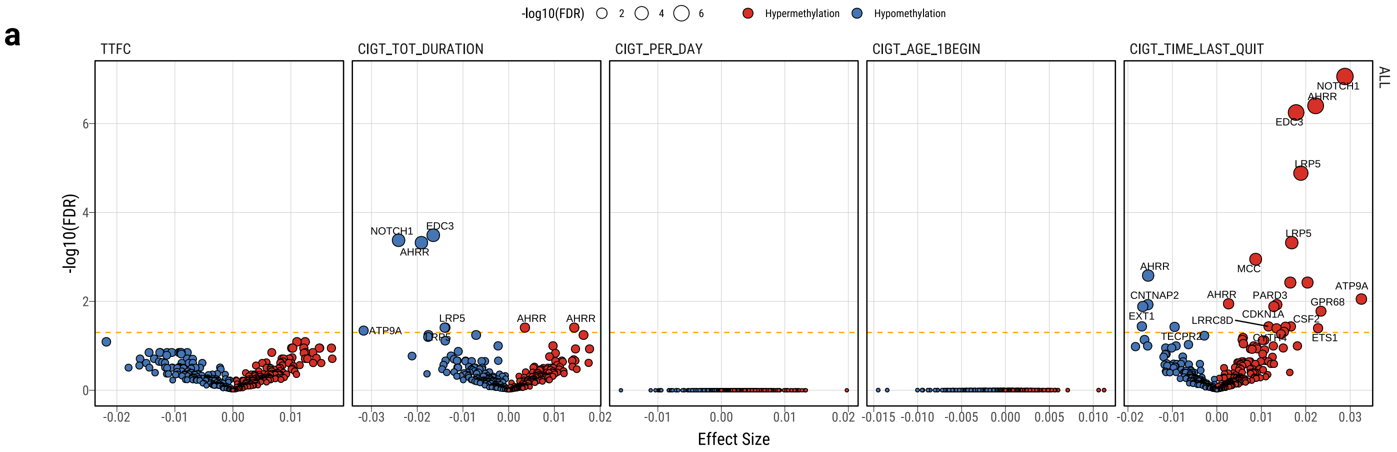


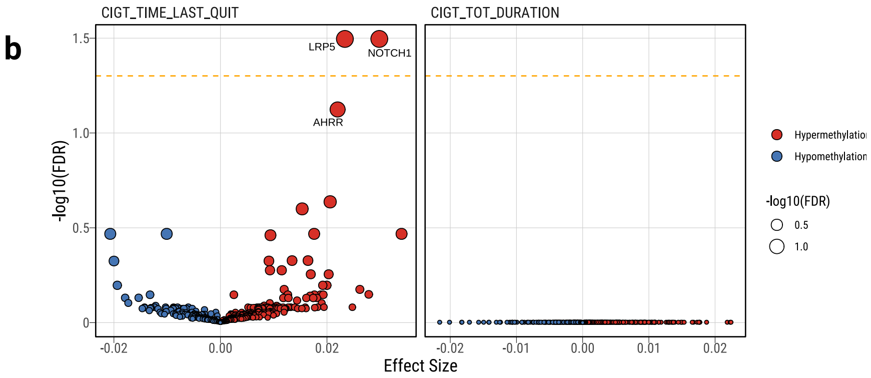
