## Supplementary_Figures for "APOBEC shapes tumor evolution and age at onset of lung cancer in smokers"

### Supplementary Fig. 1

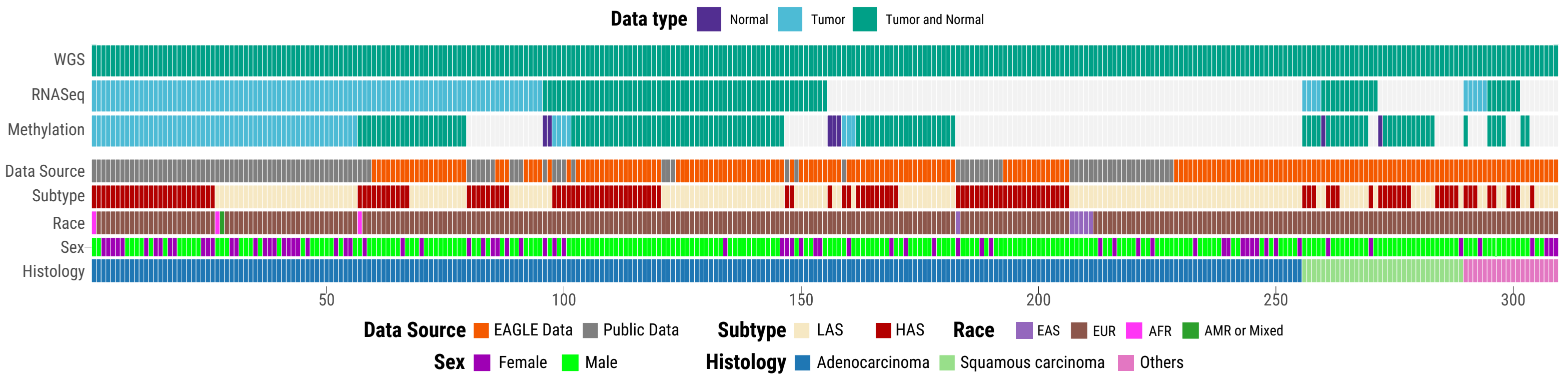

#### Supplementary Fig. 2

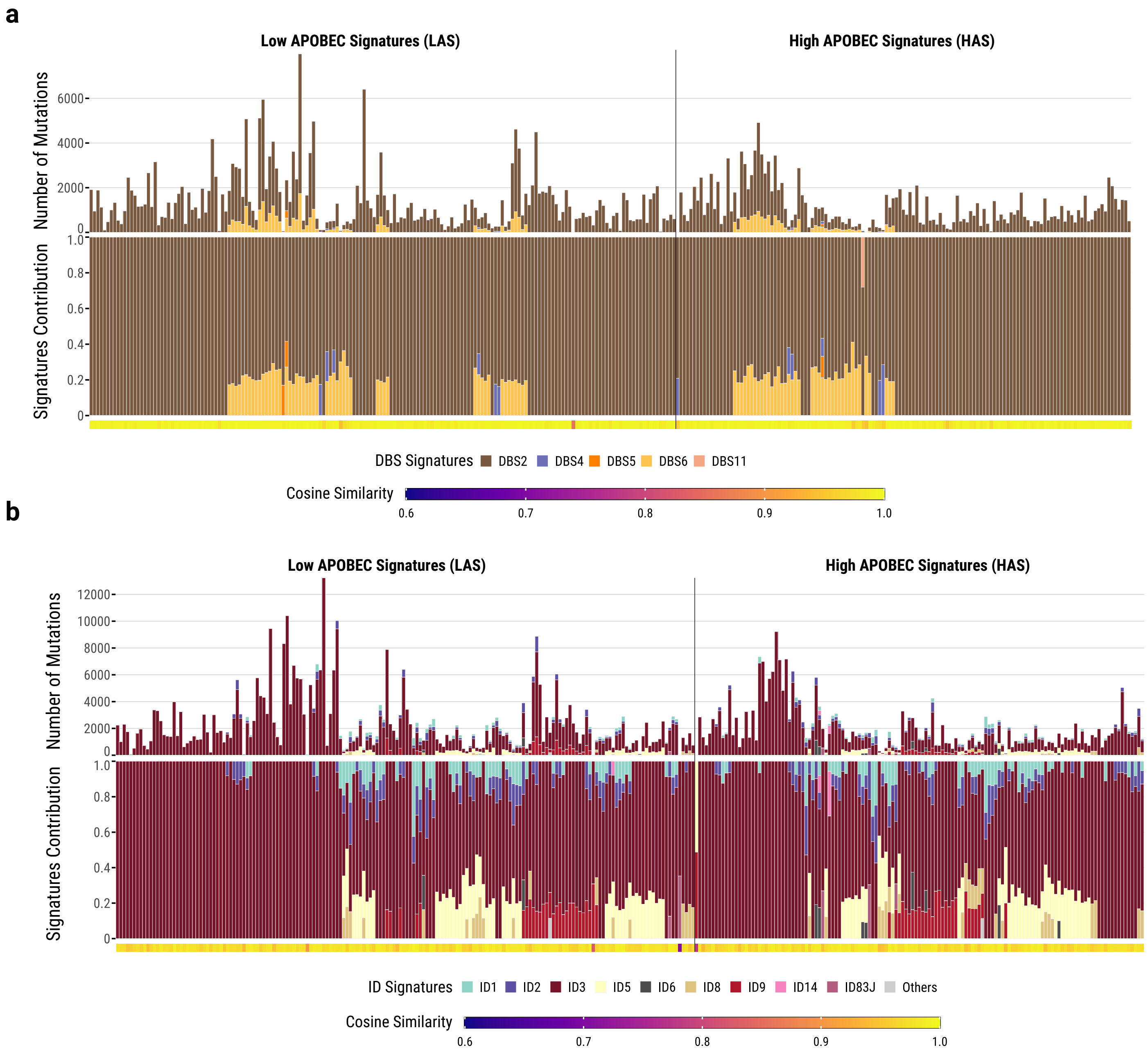

**Supplementary Fig. 3**

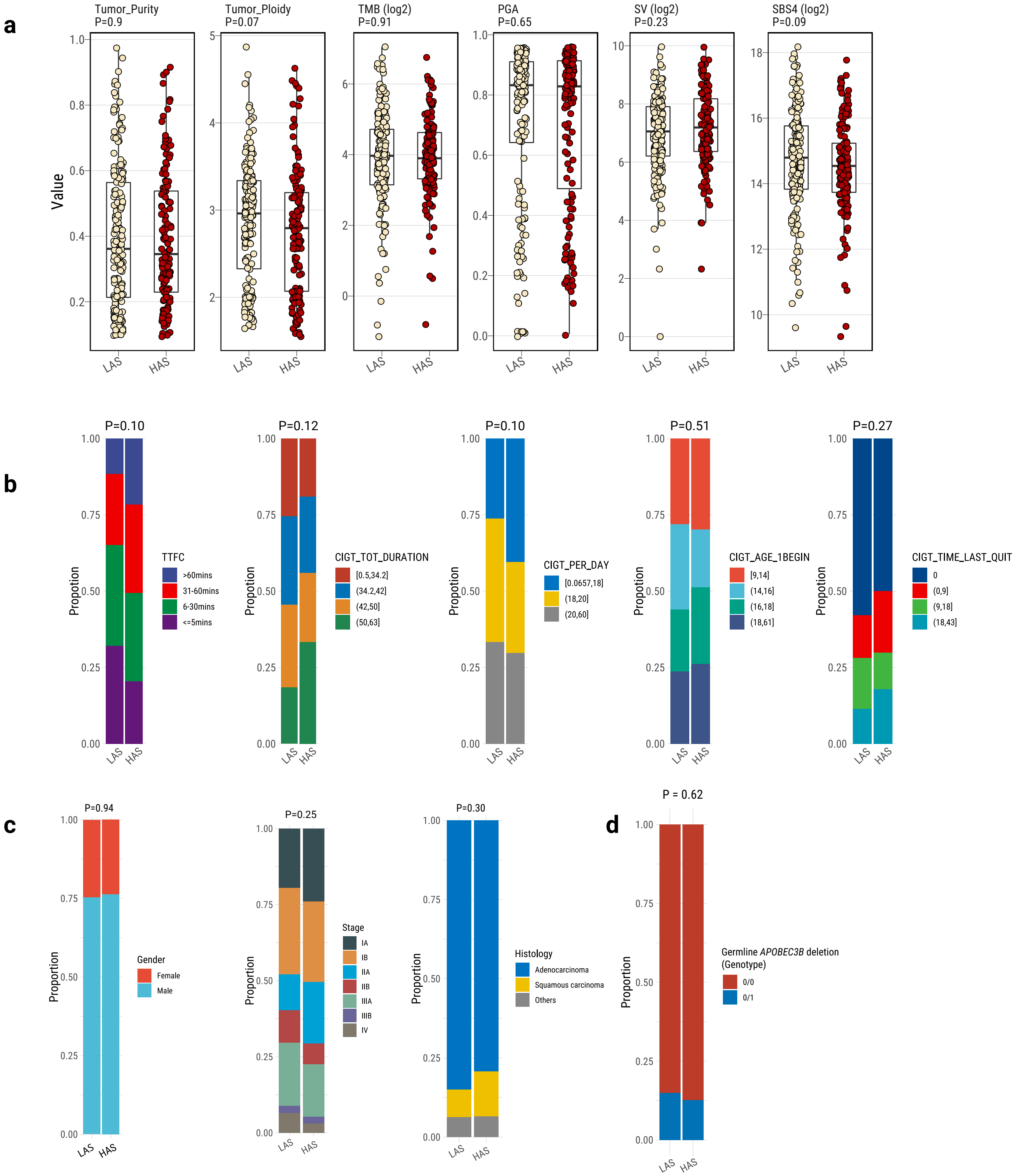

#### Supplementary Fig. 4

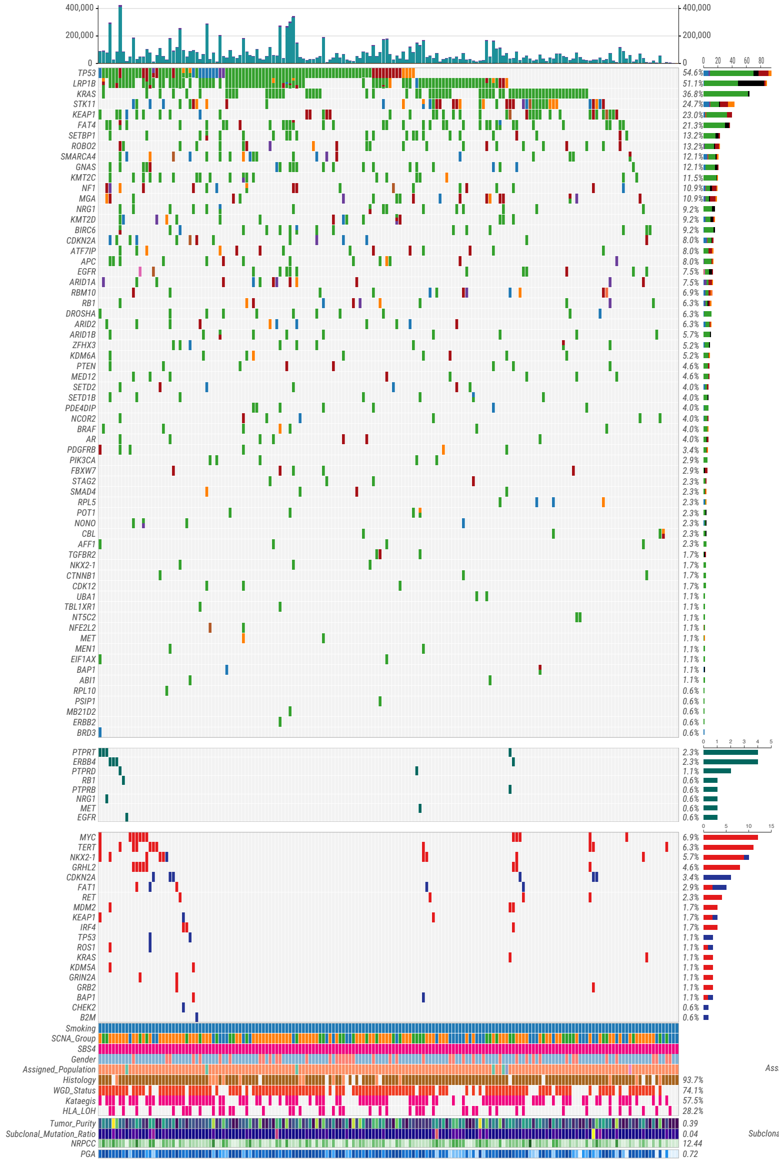


**LAS**

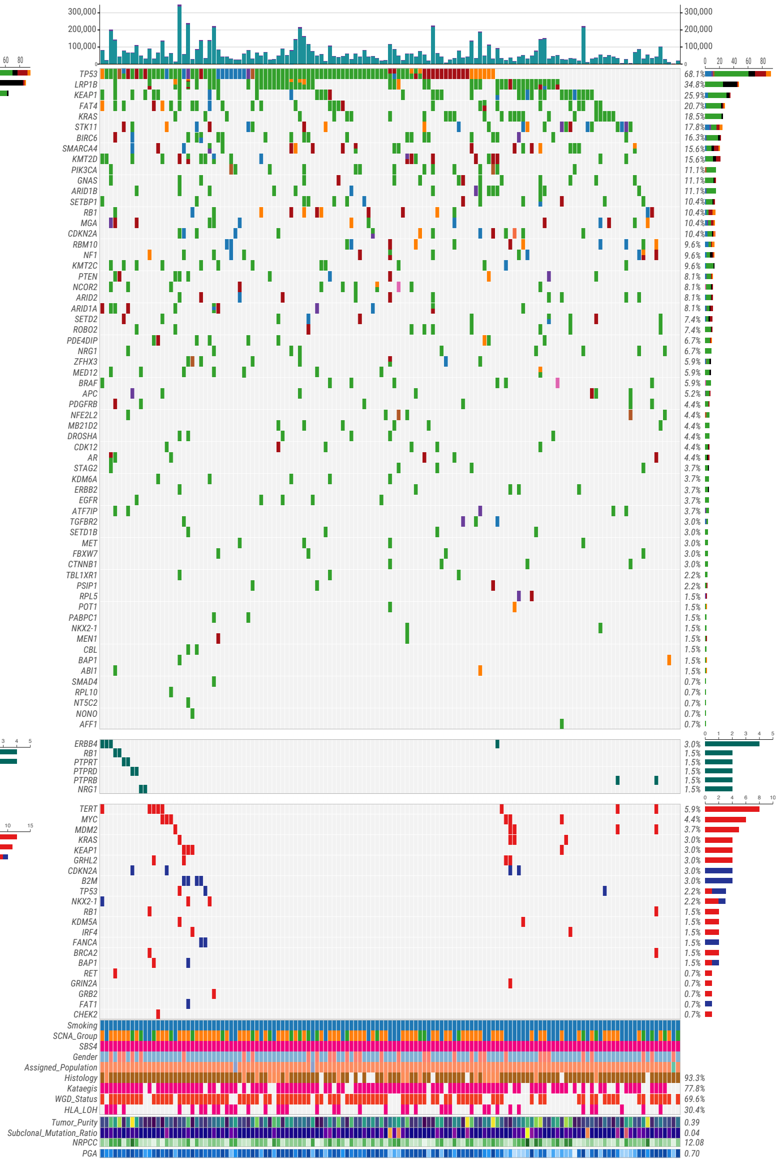


**HAS**

**Supplementary Fig. 5**

a

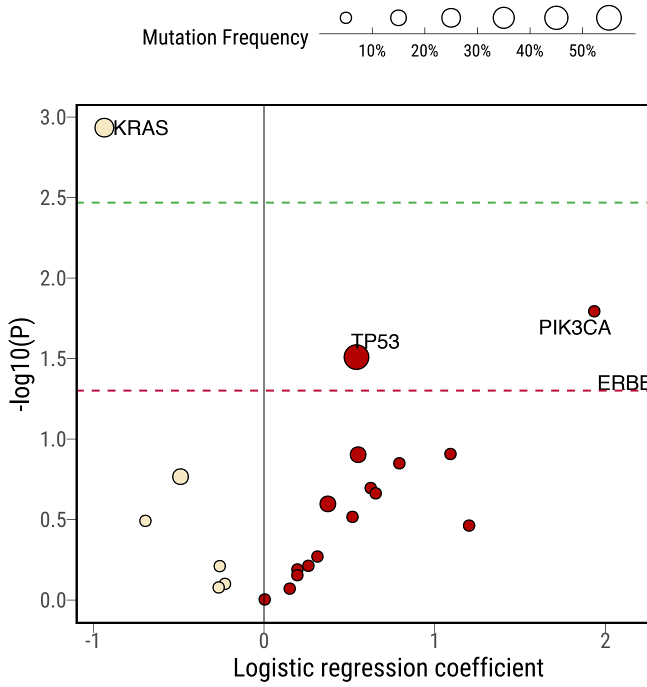


b

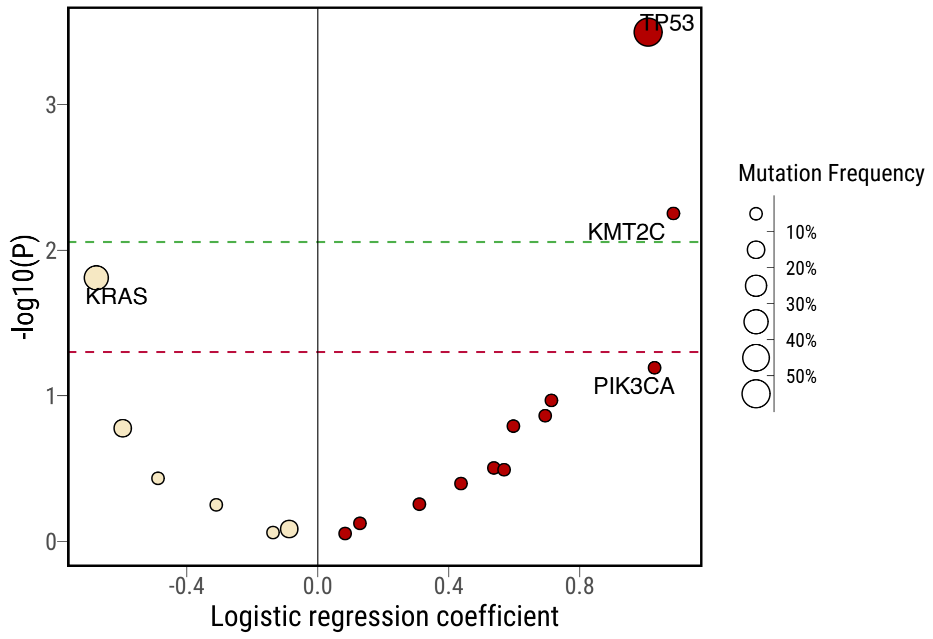

**Supplementary Fig. 6**

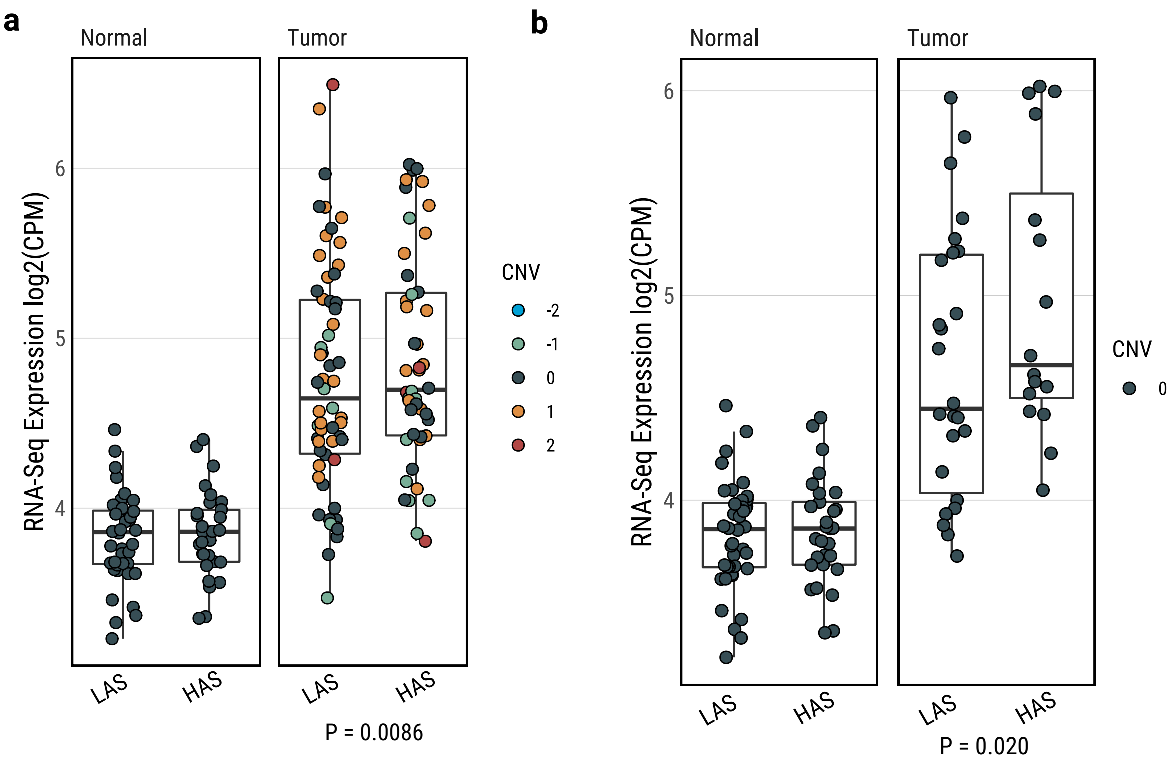

**Supplementary Fig. 7**

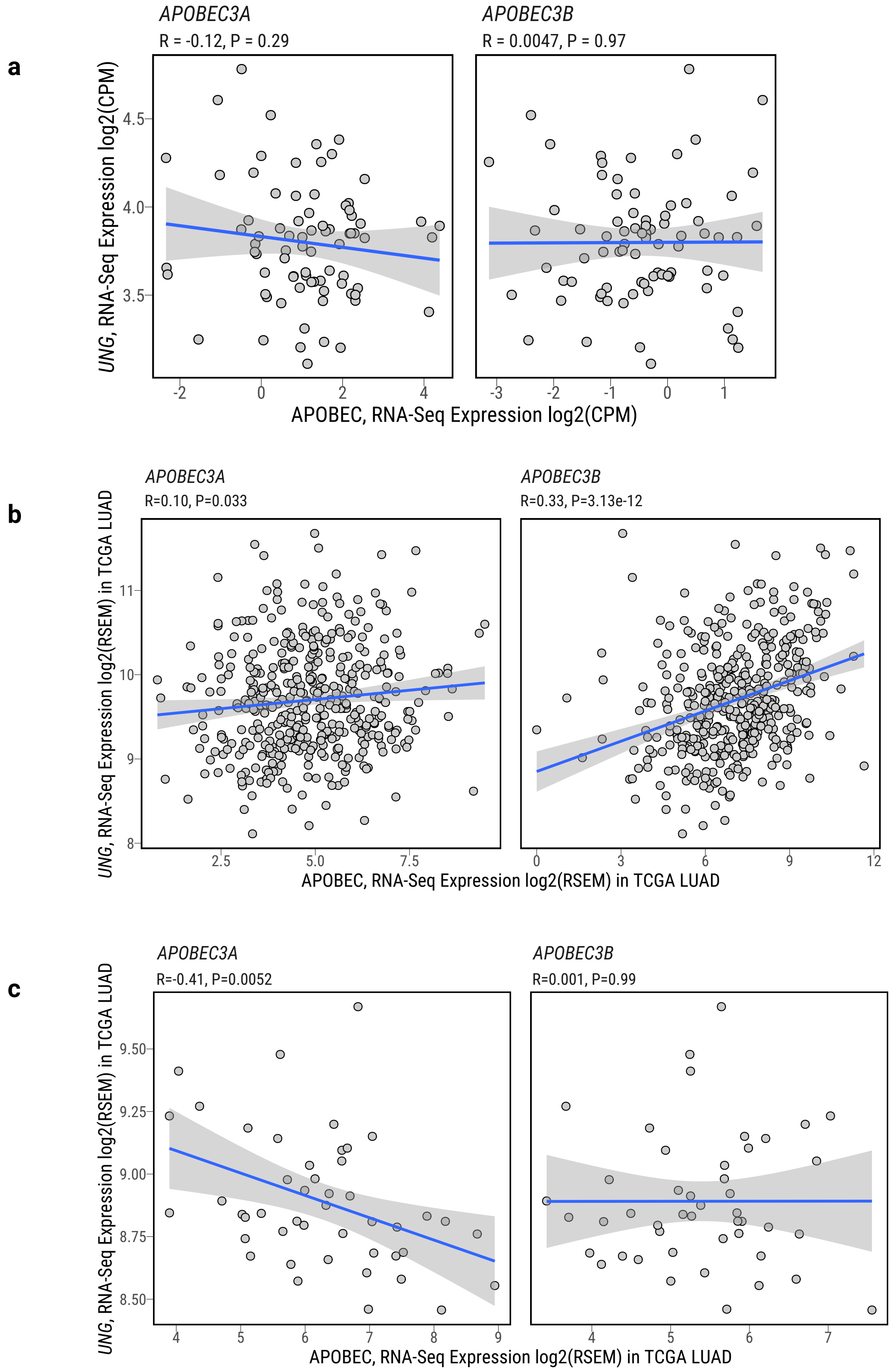

**Supplementary Fig. 8**

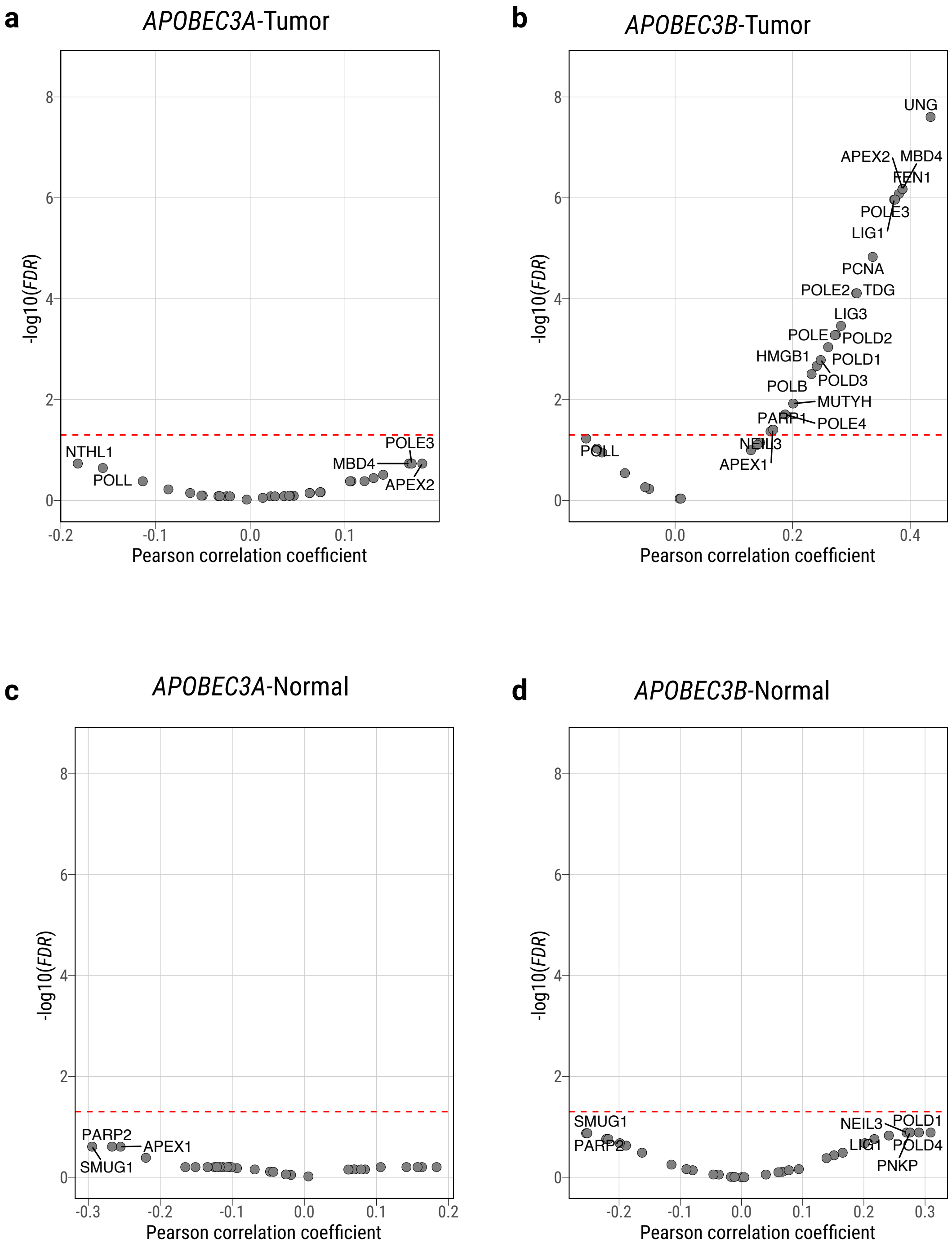

**Supplementary Fig. 9**

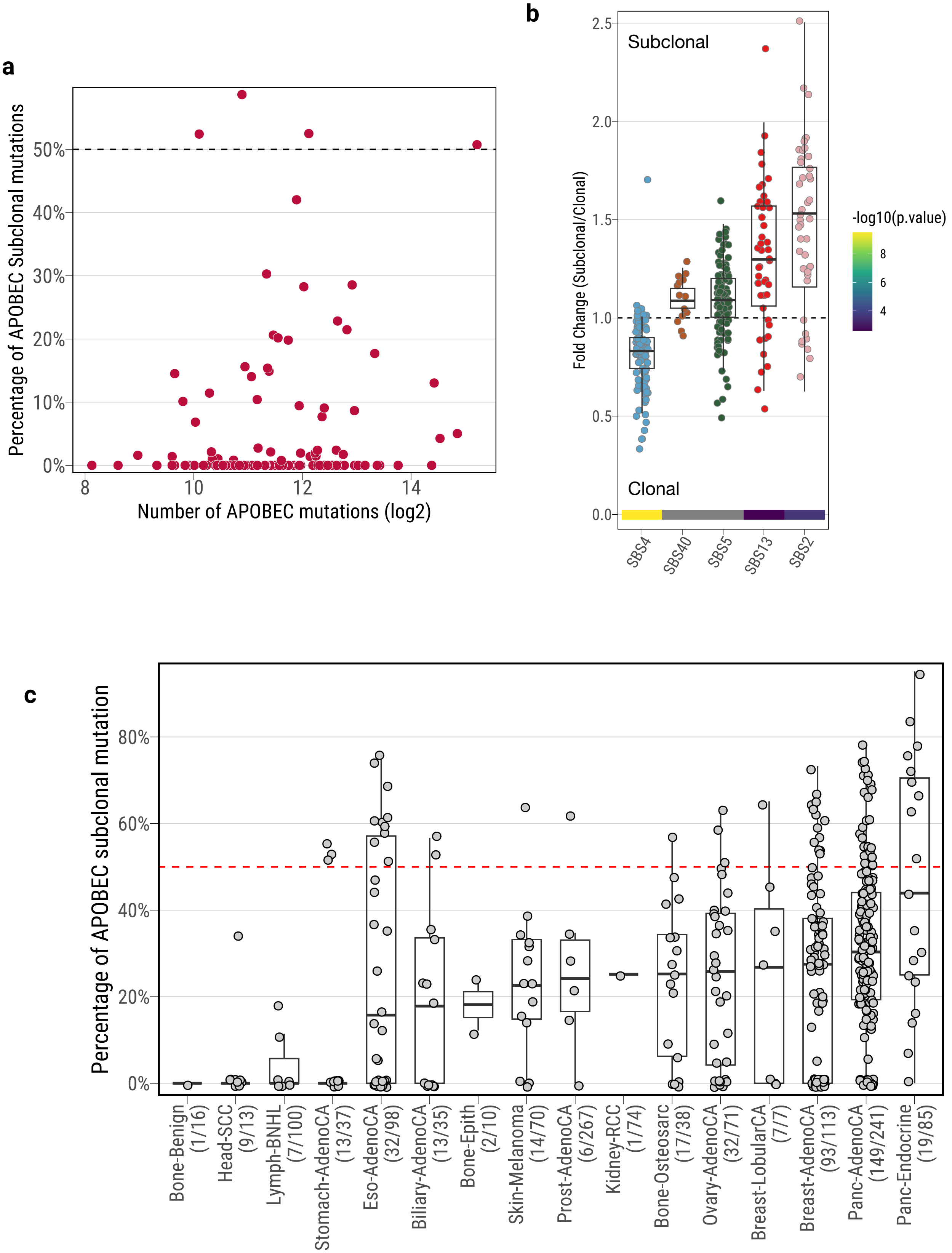

**Supplementary Fig. 10**

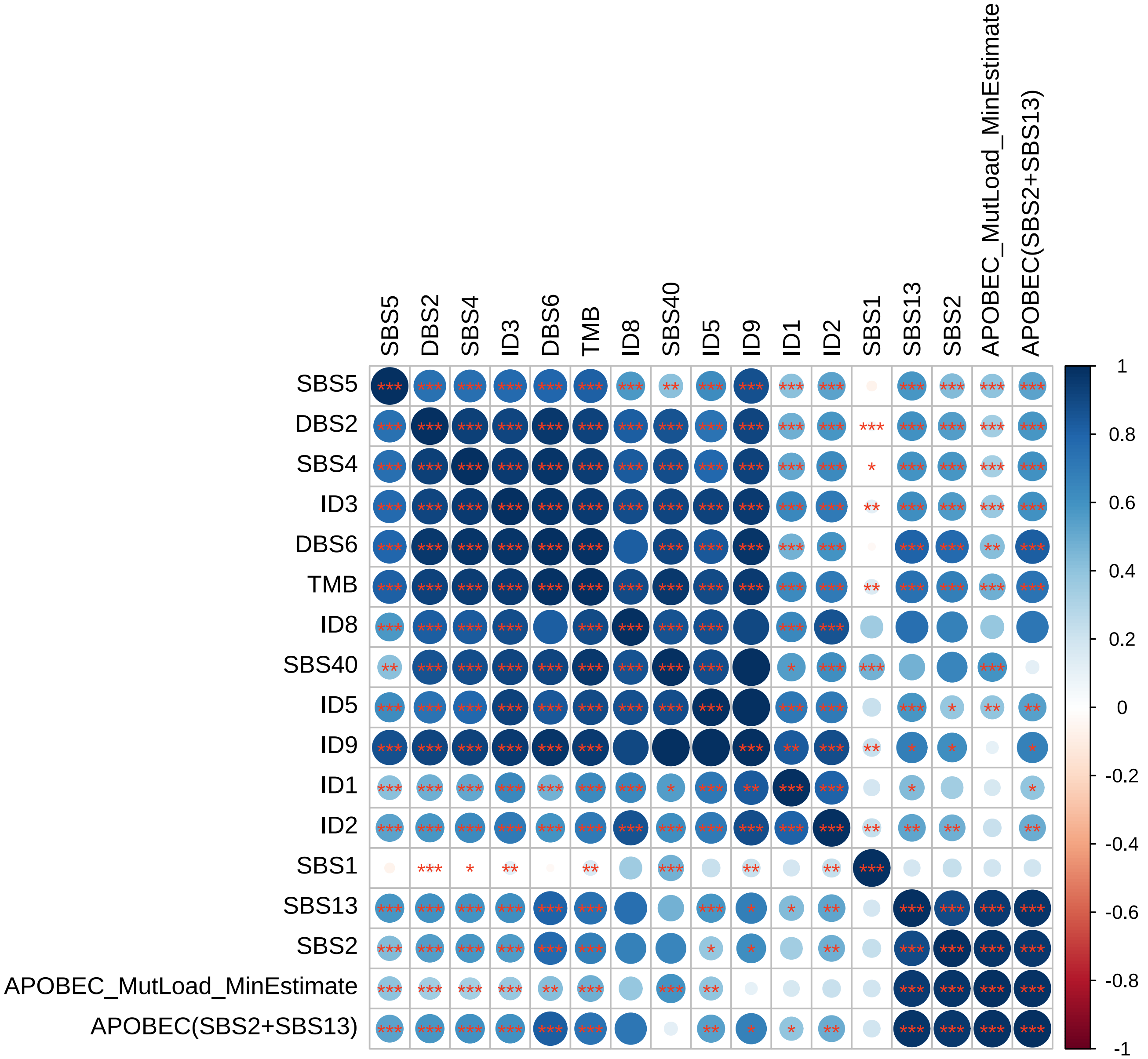

#### Supplementary Fig. 11

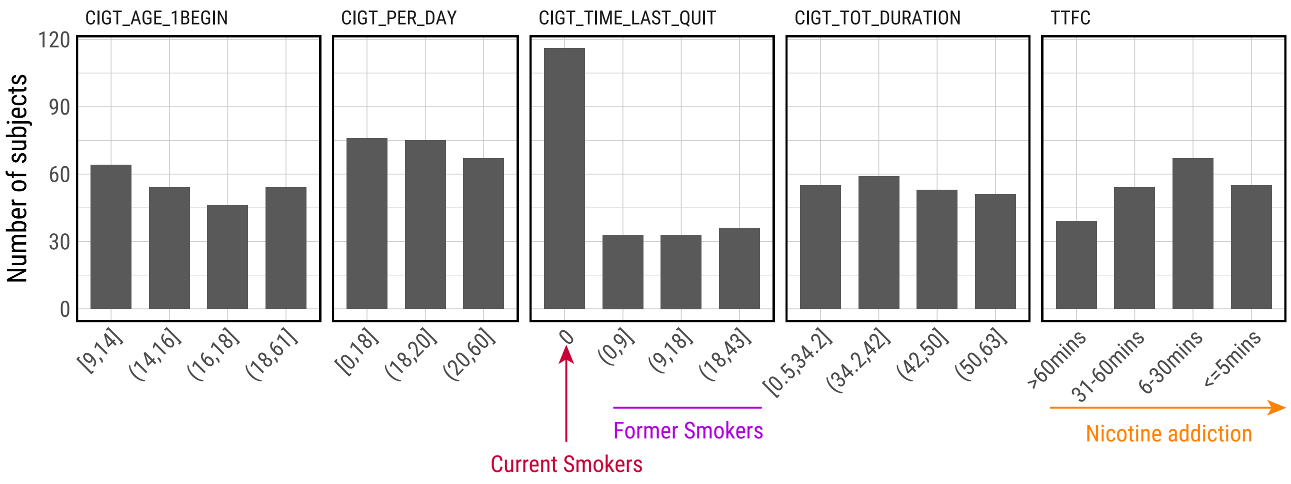


**CIGT_AGE_1BEGIN**: Age (in years) when subjects started smoking cigarettes regularly for the first time.
**CIGT_PER_DAY**: Average intensity of cigarette smoking, measured as the number of cigarettes per day.
**CIGT_TIME_LAST_QUIT**: Number of years since the subject quitted smoking cigarettes (0 means current smokers).
**CIGT_TOT_DURATION**: Total period (in years) during which the subject smoked cigarettes regularly.
**TTFC**: Time to first cigarette in the morning (from the first question of the Fagerstrom test for nicotine dependence: 'How soon after you wake up do you smoke your first cigarette?')

#### Supplementary Fig. 12

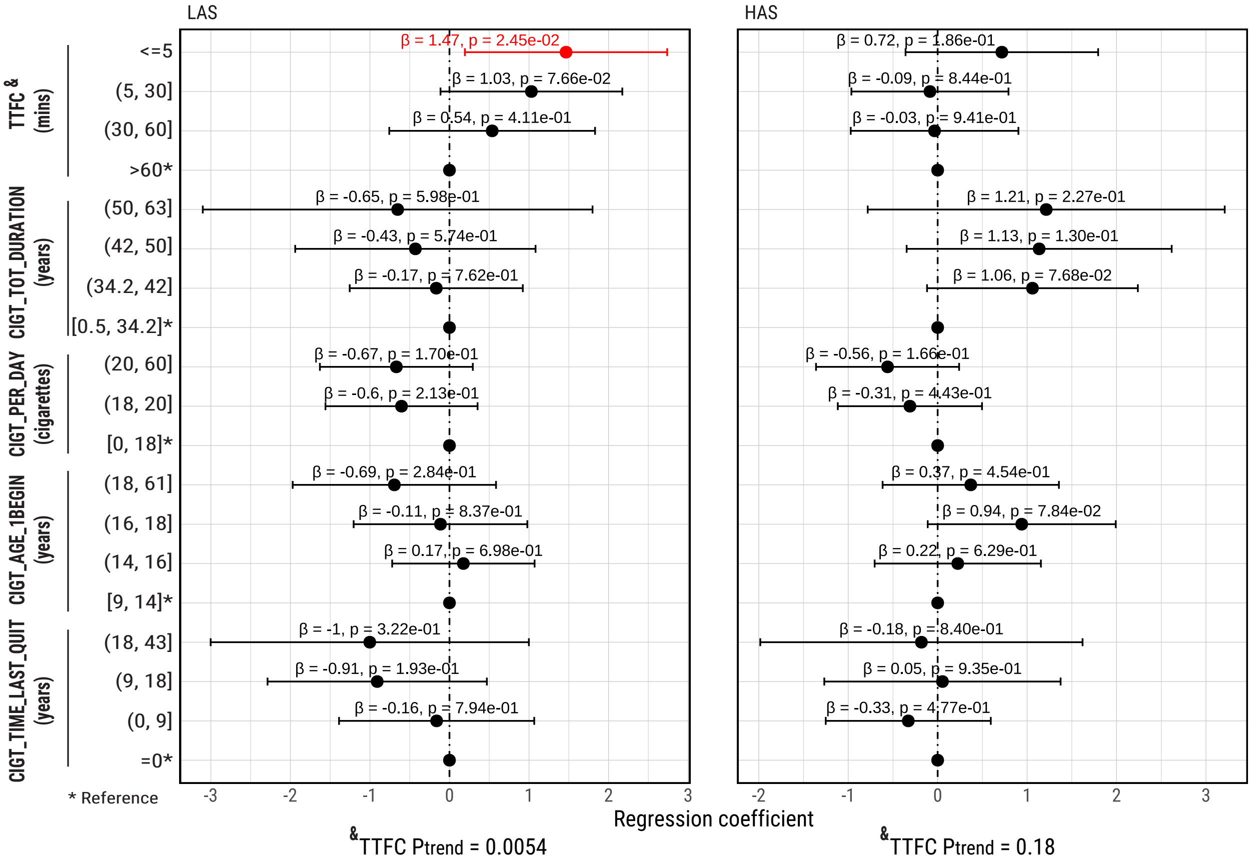

#### Supplementary Fig. 13

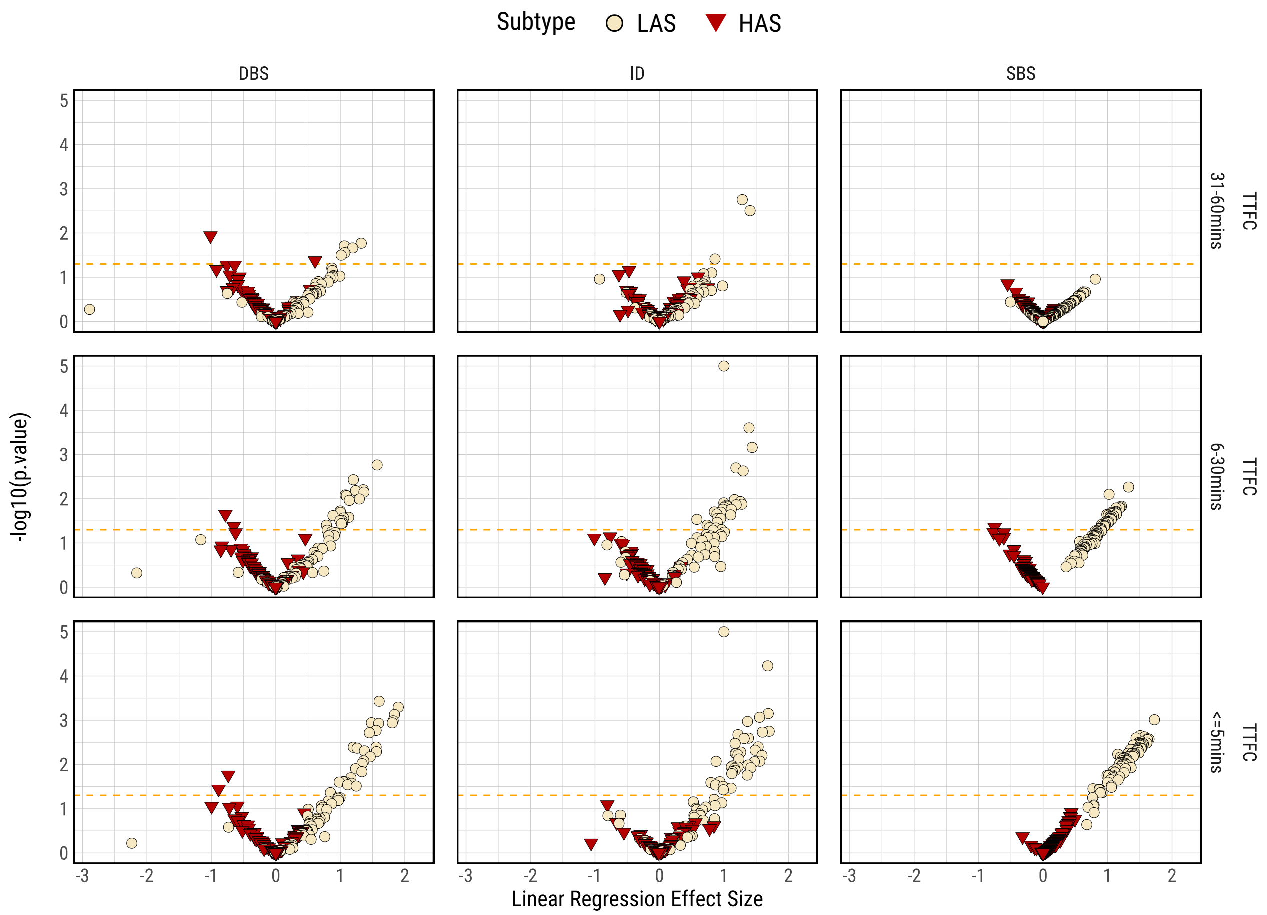

**Supplementary Fig. 14**

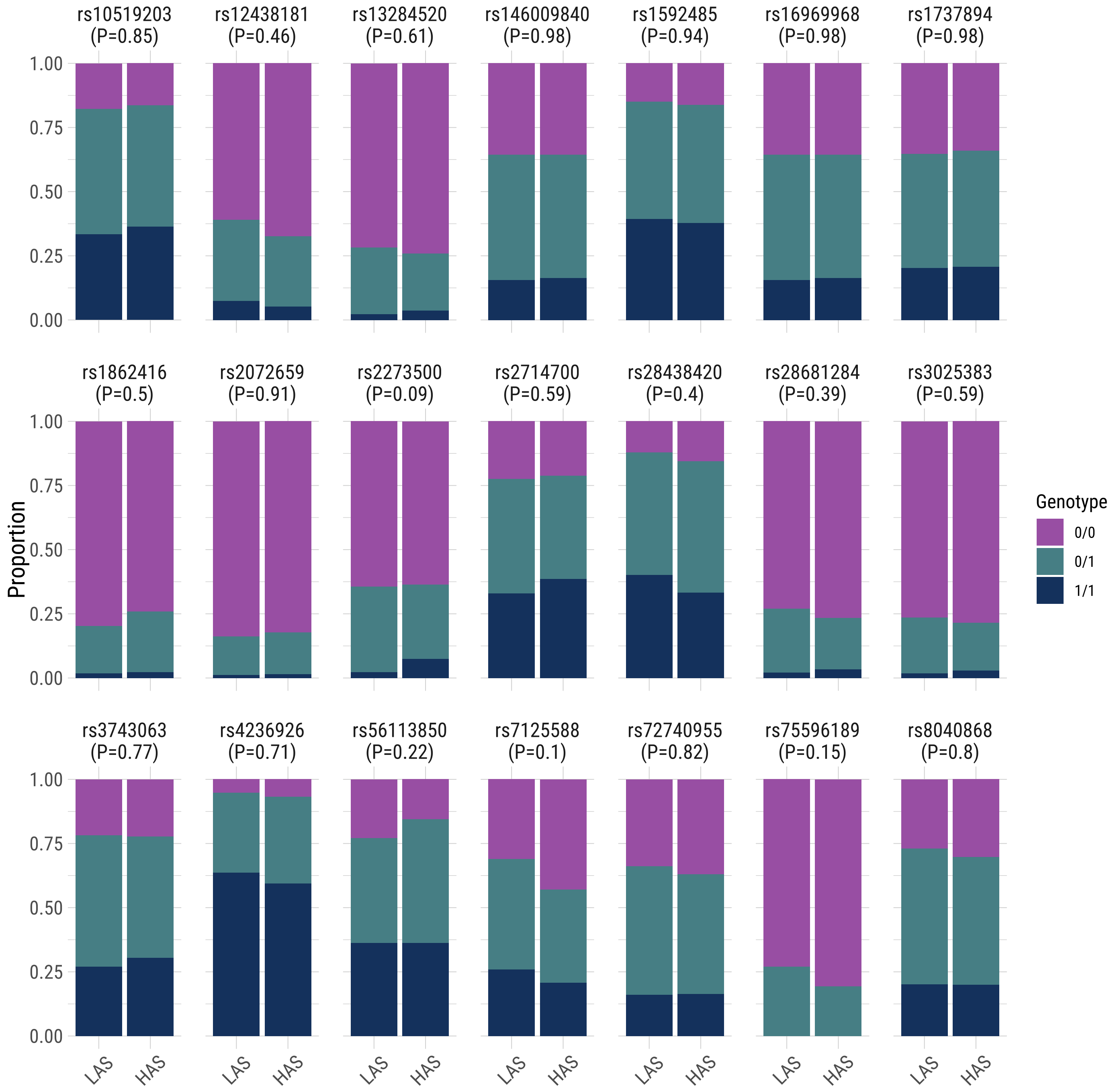

#### Supplementary Fig. 15

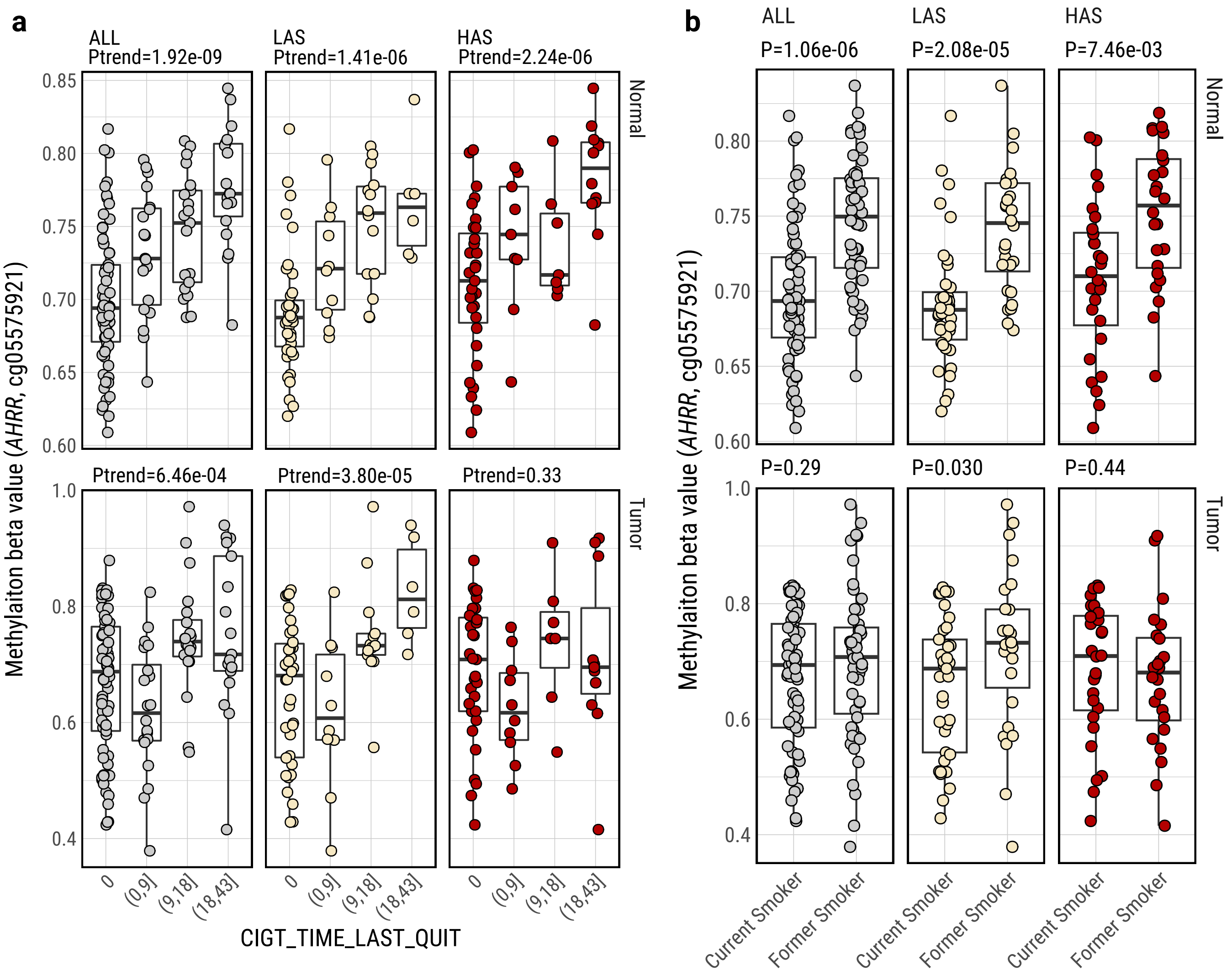

**Supplementary Fig. 16**

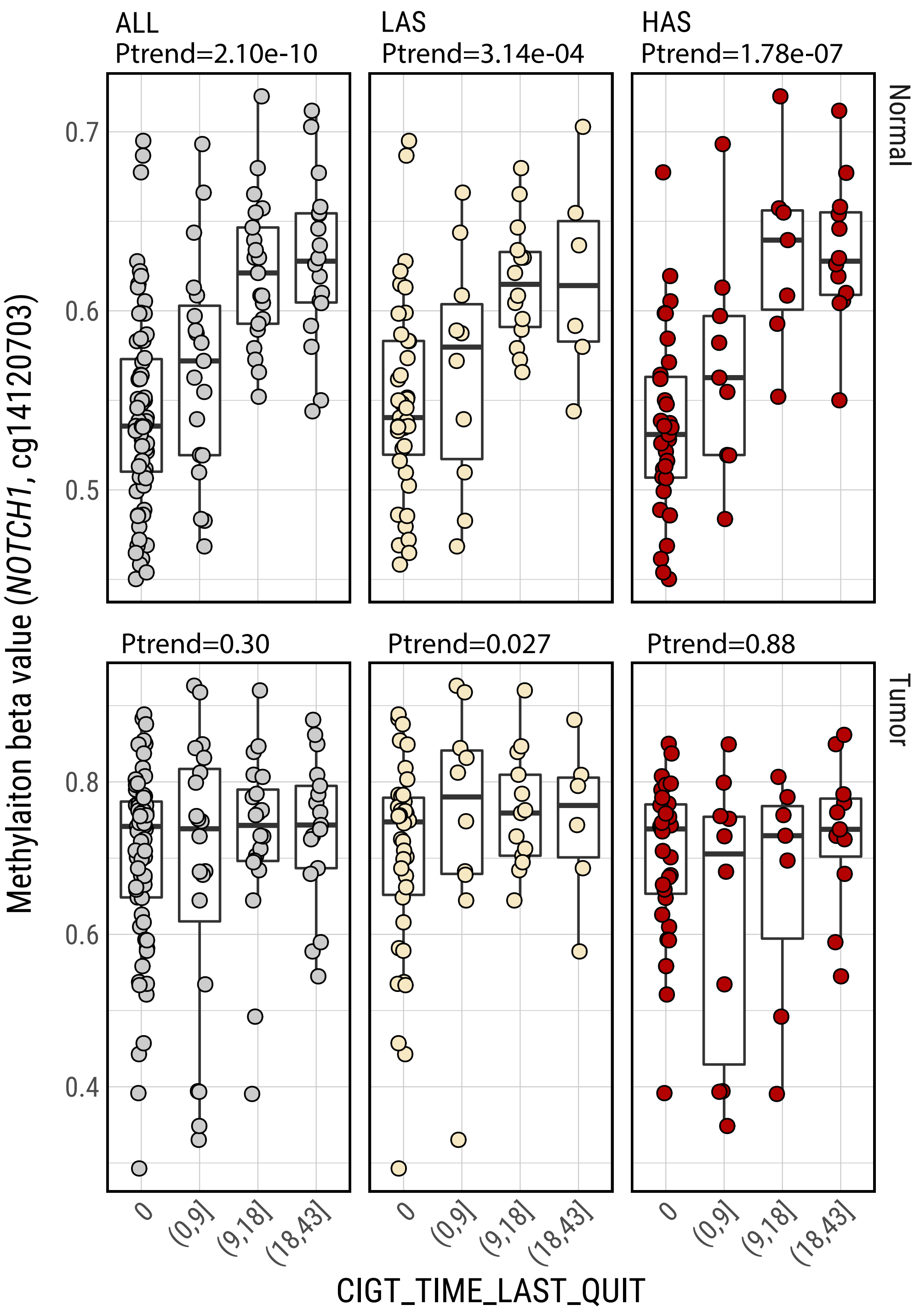

**Supplementary Fig. 17**

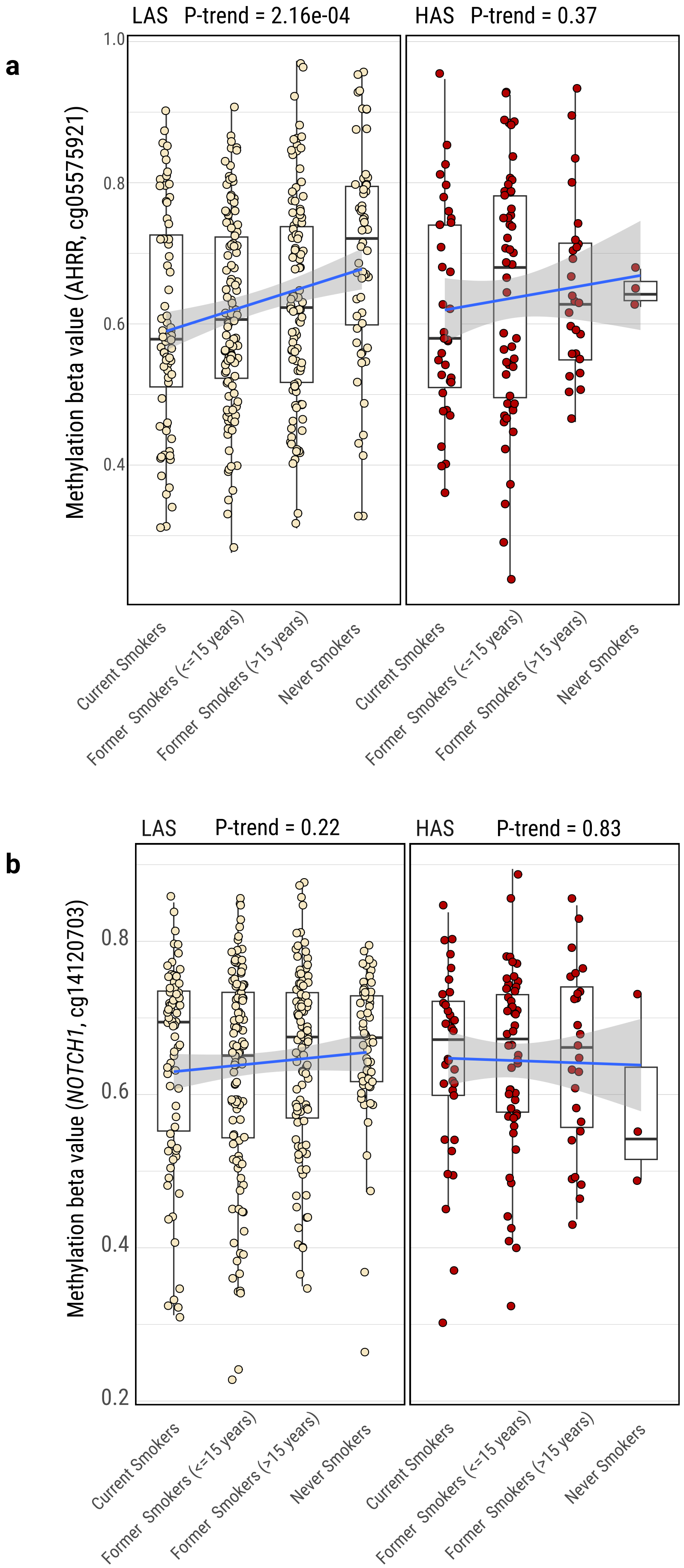

#### Supplementary Fig. 18

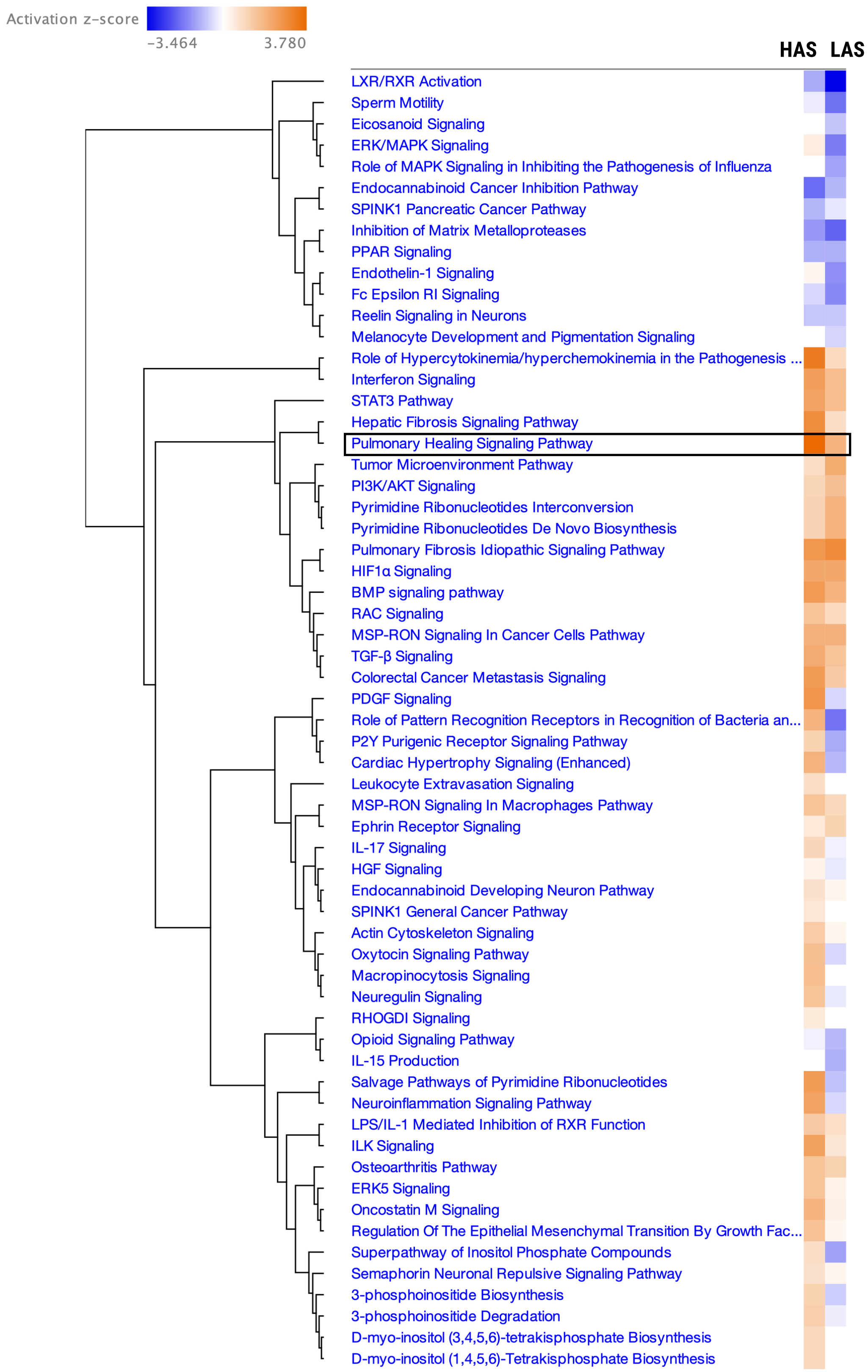

#### Supplementary Fig. 19

a

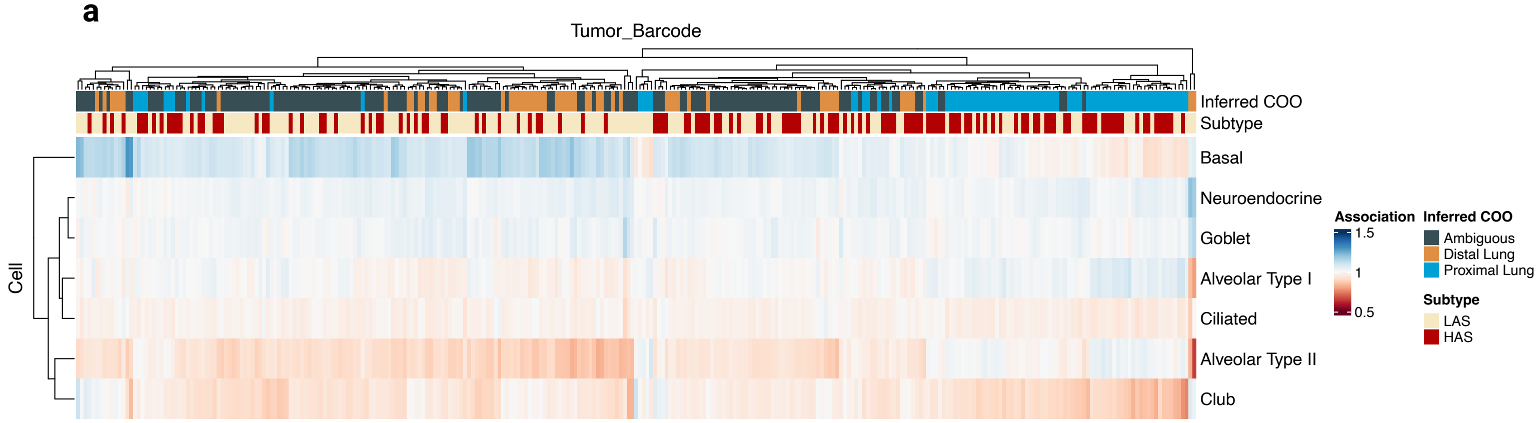


b

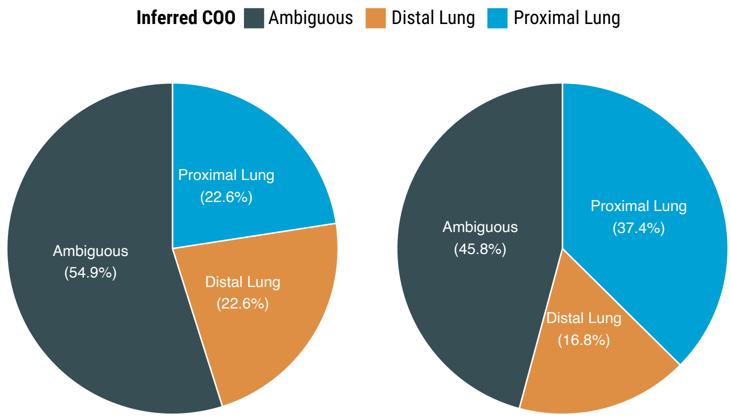

**Supplementary Fig. 20**

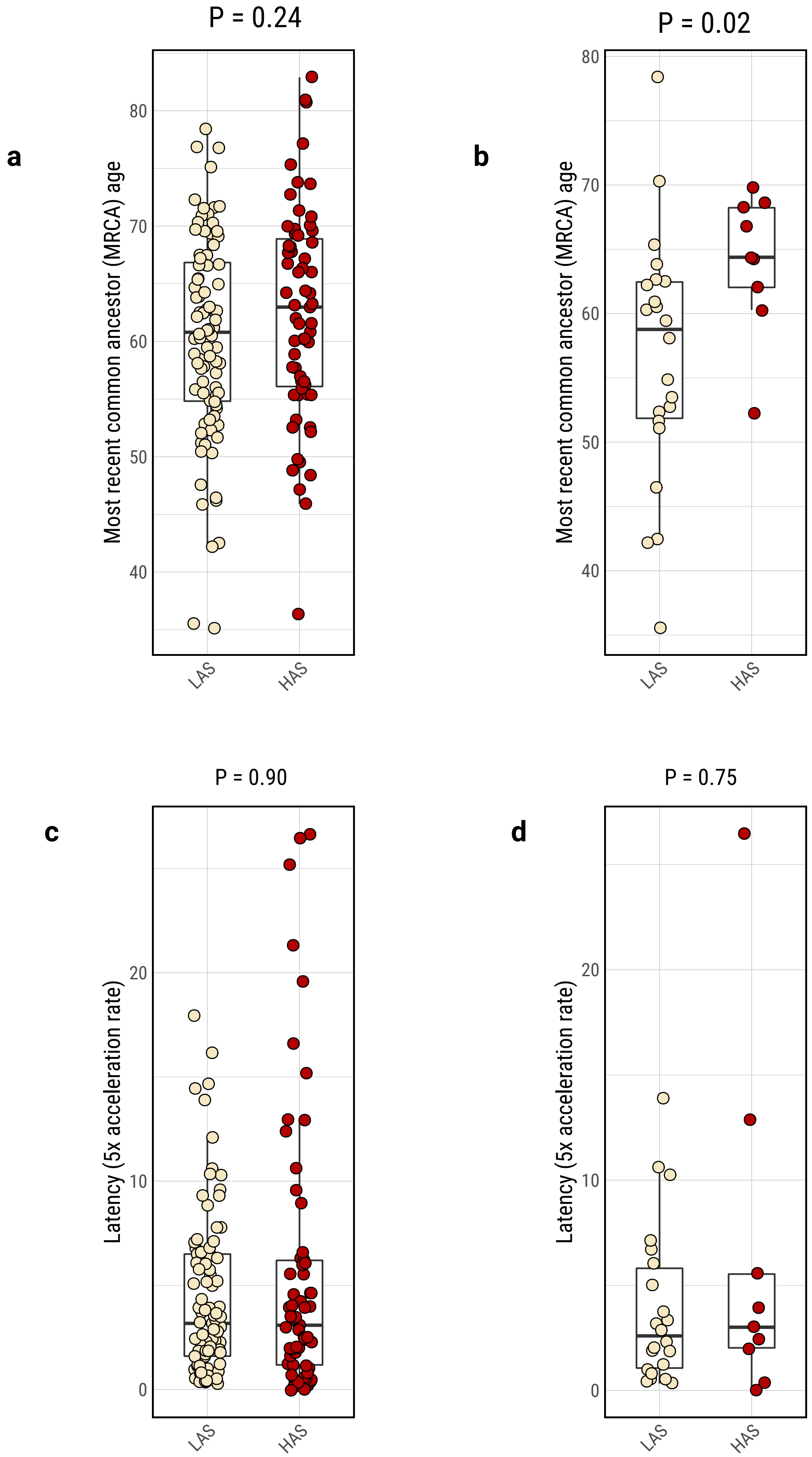

**Supplementary Fig. 21**

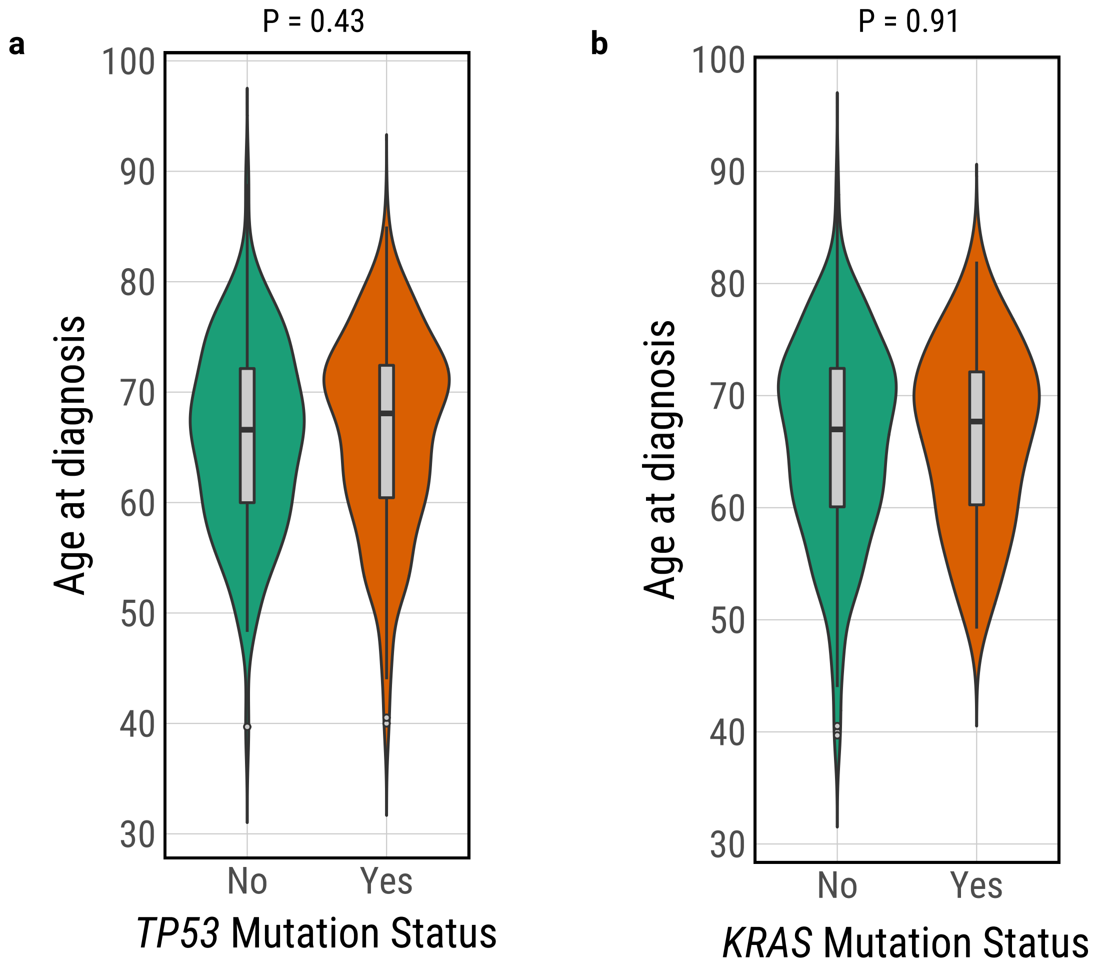

**Supplementary Fig. 22**

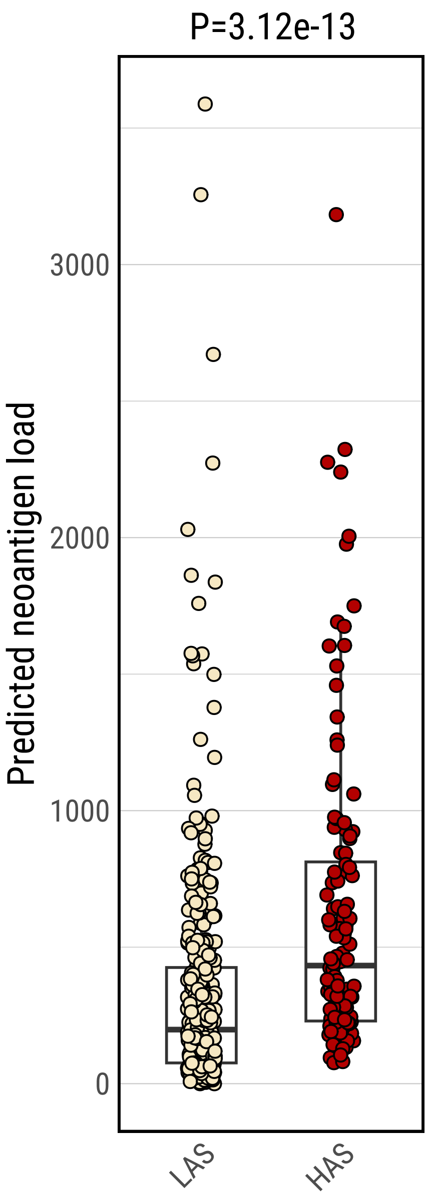

**Supplementary Fig. 23**

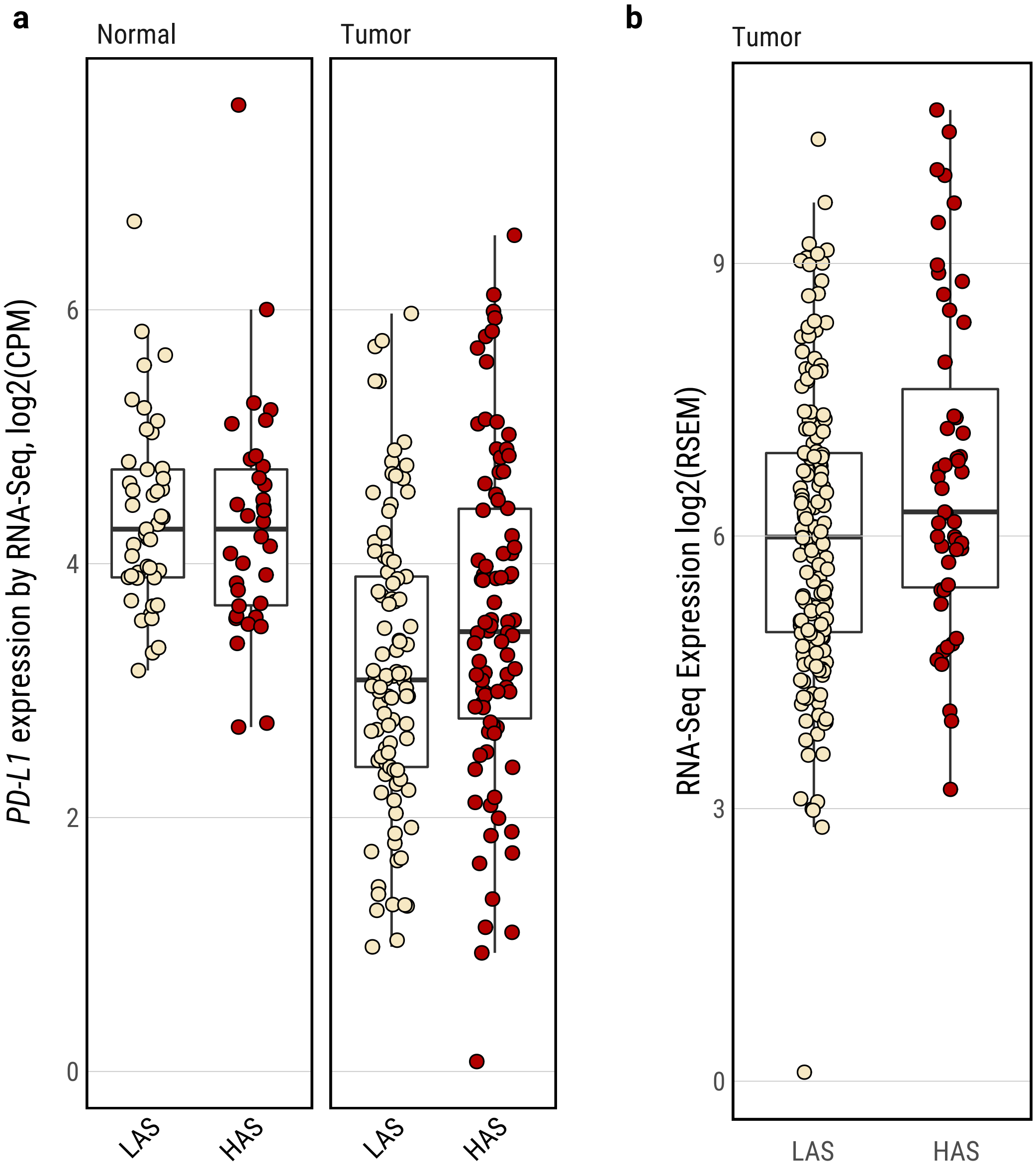

**Supplementary Fig. 24**

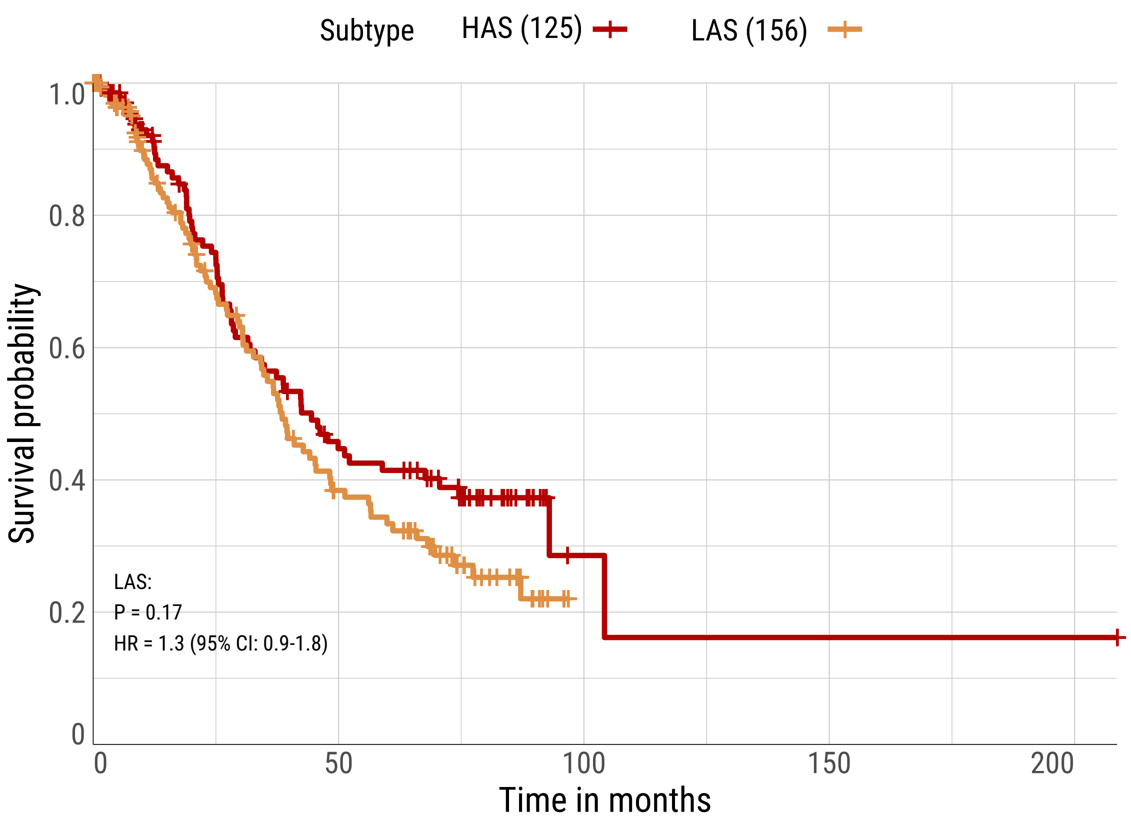

**Supplementary Fig. 25**

a

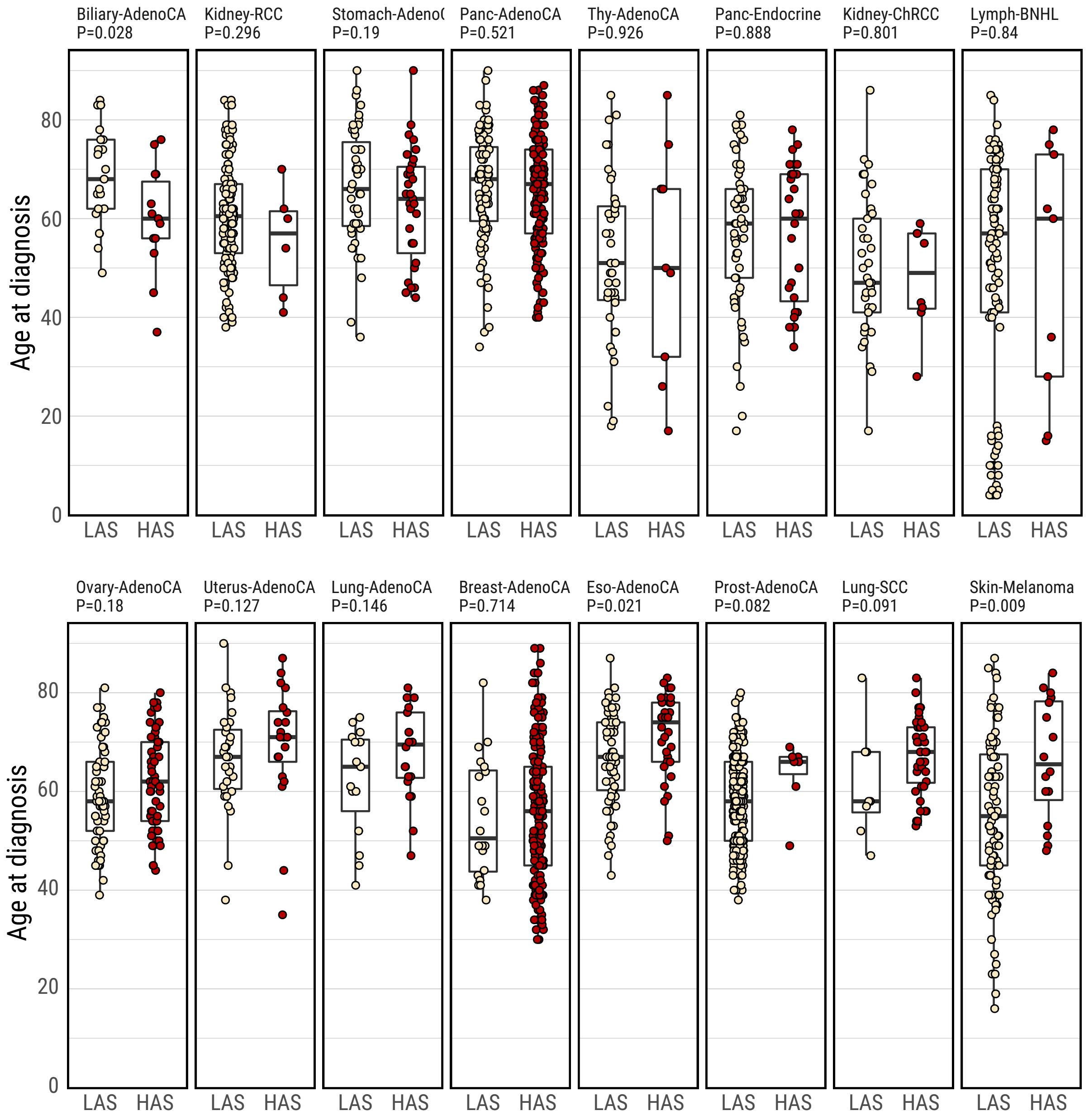


b

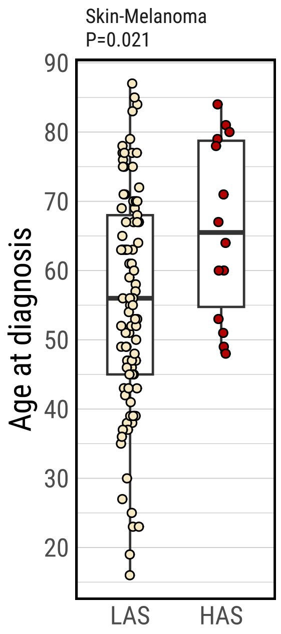

#### Supplementary Fig. 26

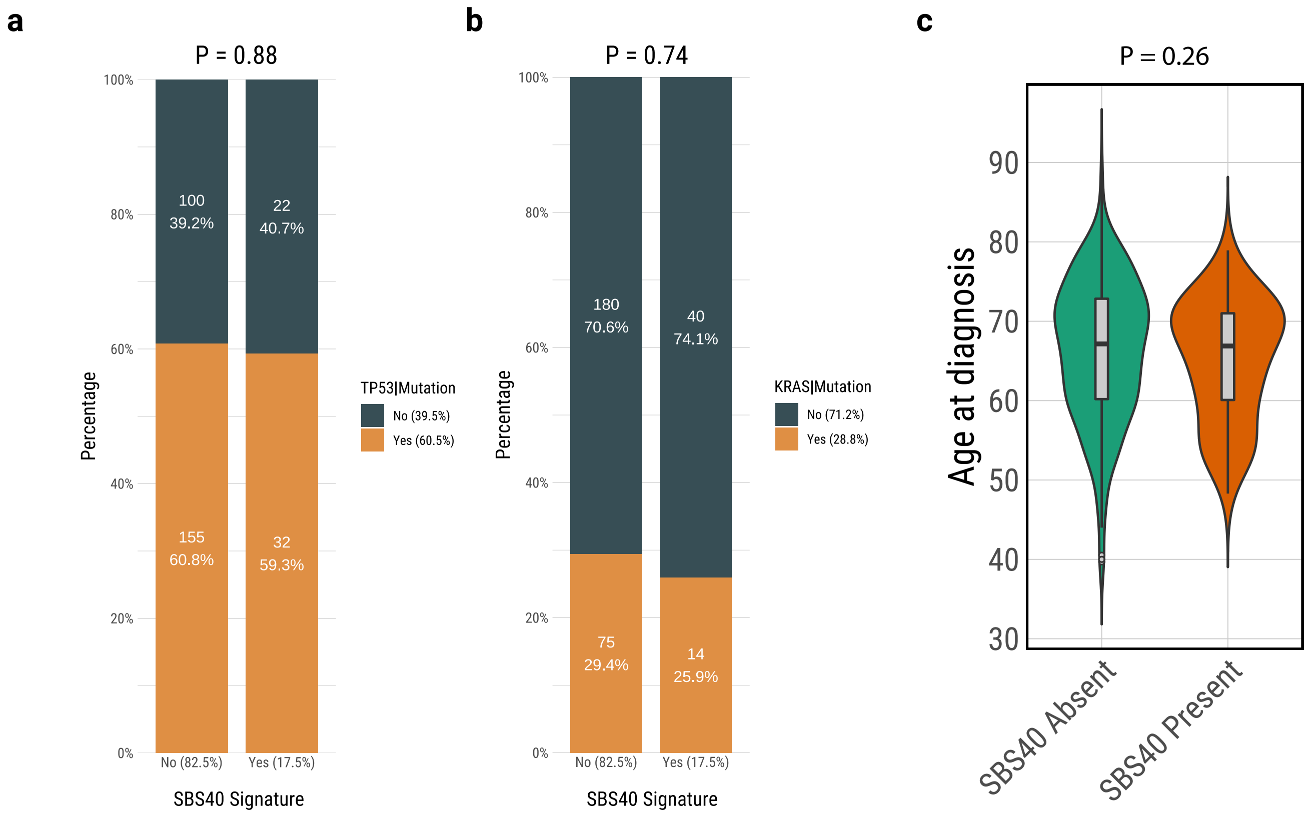
